# Proactive, Reactive, and Attentional Dynamics: An Integrative Model of Cognitive Control

**DOI:** 10.1101/2024.10.01.615613

**Authors:** Percy K. Mistry, Stacie L. Warren, Nicholas K. Branigan, Weidong Cai, Vinod Menon

## Abstract

We developed a novel Proactive Reactive and Attentional Dynamics (PRAD) computational model designed to dissect the latent mechanisms of inhibitory control in human cognition. Leveraging data from over 7,500 participants in the NIH Adolescent Brain Cognitive Development study, we demonstrate that PRAD surpasses traditional models by integrating proactive, reactive, and attentional components of inhibitory control. Employing a hierarchical Bayesian framework, PRAD offers a granular view of the dynamics underpinning action execution and inhibition, provides debiased estimates of stop-signal reaction times, and elucidates individual and temporal variability in cognitive control processes. Our findings reveal significant intra-individual variability, challenging conventional assumptions of random variability across trials. By addressing nonergodicity and systematically accounting for the multicomponential nature of cognitive control, PRAD advances our understanding of the cognitive mechanisms driving individual differences in cognitive control and provides a sophisticated computational framework for dissecting dynamic cognitive processes across diverse populations. Our integrative approach significantly advances psychological theory about the multiple neurocognitive processes underlying cognitive control. We also demonstrate the relevance of our theoretical and modeling framework to understand the cognitive, neural, clinical, and exposomic factors related to cognitive control.

## Main

Human cognition is a dynamic process, which relies in part, on goal-directed beliefs about task characteristics, context-dependent flexible action control, the capacity to learn from a history of decisions and consequences, leading to moment-to-moment adaptation of response strategies to optimize behavioral outcomes (A. R. Aron, 2011; T. S. Braver, 2012; Todd S Braver, Paxton, Locke, & Barch, 2009; Cai et al., 2016; Menon, Adleman, White, Glover, & Reiss, 2001). Impairments in cognitive systems that regulate such dynamic processes underlying everyday cognitive functioning are a hallmark of psychopathology (Casey, Tottenham, Liston, & Durston, 2005; Menon, 2011, 2013; Nigg, 2017; Toga, Thompson, & Sowell, 2006). Identifying latent cognitive factors that drive adaptive and maladaptive behavioral dynamics is critical for understanding individual differences in how cognitive processes unfold over time, and how their associated alterations in regulatory systems affect symptom presentation in neuropsychiatric disorders (Hall, Bernat, & Patrick, 2007; Ivanov, Schulz, London, & Newcorn, 2008; Korenblum, Chen, Manassis, & Schachar, 2007; Murray & Kochanska, 2002; Olvet & Hajcak, 2008; Oosterlaan & Sergeant, 1996; R. Schachar, Tannock, Marriott, & Logan, 1995; R. J. Schachar et al., 2004). However, conventional methods are often unable to reveal multicomponential latent constructs that govern dynamic cognitive control processes.

Here, we develop Proactive Reactive and Attentional Dynamics (PRAD), a novel computational model to characterize and measure latent cognitive constructs that govern behavioral dynamics of action execution and inhibition, whose deficits are often associated with multiple psychiatric disorders including attention deficit hyperactivity disorder, autism, substance abuse, and schizophrenia. This model was applied to a response inhibition paradigm in the large-scale (N > 7500) Adolescent Brain Cognitive Development (ABCD) (B. Casey et al., 2018) study to uncover the mechanisms by which dynamic cognitive processes involved in response initiation and inhibition are regulated, and the nature of individual differences in the latent cognitive constructs associated with such adaptive and maladaptive regulation.

This paper is structured as follows: We first motivate the need for PRAD by identifying that a large proportion of individuals representative of the general population demonstrate behavior that is known to cause issues in traditional measurements of response inhibition, or demonstrate behavior that highlights the needs greater process dissociation and mechanistic explanations of behavior. We then demonstrate the adequacy of the PRAD model in explaining aggregate patterns and individual level of behavior, show its added utility over cognitive models without these dynamic mechanisms, and importantly, provide theoretical advances on the role of proactive, reactive, and attentional mechanisms in cognitive control. We show how PRAD allows debiasing inferences, improves our understanding of the role of adaptivity, our understanding of individual differences, and ability to predict cognitive performance. We also show the relevance of PRAD to understanding clinical subgroups, to our understanding of the neural underpinnings of cognitive control, and to understanding the role of exposomic factors.

### Inhibitory Control and the Stop-Signal Paradigm

Inhibitory control, the ability to withhold or cancel undesirable action, thought, and emotion, is fundamental to goal-directed behaviors (Alderson, Rapport, & Kofler, 2007; Bonham, Shanley, Waters, & Elvin, 2021; Lavagnino, Arnone, Cao, Soares, & Selvaraj, 2016; Lipszyc & Schachar, 2010; McTeague et al., 2017; R. Schachar & Logan, 1990; Smith, Mattick, Jamadar, & Iredale, 2014). The stop-signal task (SST, **Figure 1**) is a widely used paradigm (Frederick Verbruggen et al., 2019; Verbruggen & Logan, 2008) to study inhibitory control mechanisms and their neural underpinnings. The SST involves making a response to a *Go* signal but inhibiting the prepared response when the *Go* signal is quickly followed by an infrequent *Stop* signal. The time interval between Go and Stop signals is called the stop-signal delay (SSD) and is experimentally manipulated. On stop signal trials with longer SSD, the prepotent Go response is cognitively further along, and more difficult to stop after detecting the Stop signal. The SST has been used in a variety of domains, including non-human primates (Ghasemian, Vardanjani, Sheibani, &

**Figure 1.**
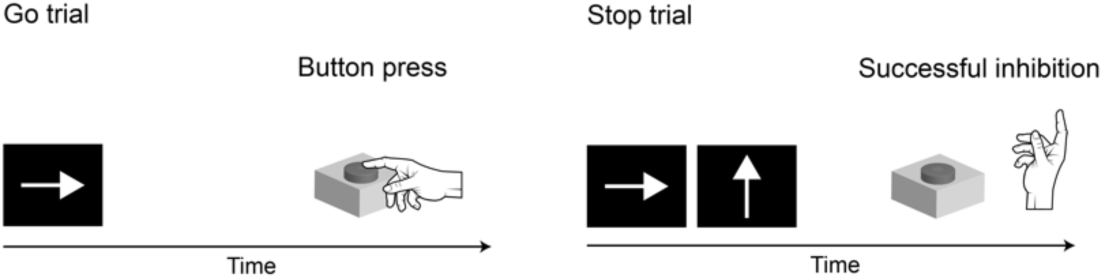
Overview of the stop signal task and analysis pipeline. Illustration of Go and Stop trials in the stop signal task. On Go trials, participants respond to indicate arrow direction; on Stop trials, they must inhibit their response when the stop signal appears.

Mansouri, 2021; Godlove et al., 2011; Liu, Heitz, & Bradberry, 2009), rodents (Bari et al., 2011; Caglayan, Stumpenhorst, & Winter, 2021; Eagle et al., 2009; Mayse, Nelson, Park, Gallagher, & Lin, 2014), during development in children (Cai et al., 2019; Carver, Livesey, & Charles, 2001; R. Schachar & Logan, 1990; Williams, Ponesse, Schachar, Logan, & Tannock, 1999), through the adult human life span (Hsu & Hsieh, 2022; Tsvetanov et al., 2018), in neurodiverse populations (Adams & Jarrold, 2012; Alderson et al., 2007; Bonham et al., 2021; Geurts, van den Bergh, & Ruzzano, 2014; Schmitt, White, Cook, Sweeney, & Mosconi, 2018; Senderecka, Grabowska, Szewczyk, Gerc, & Chmylak, 2012; Soreni, Crosbie, Ickowicz, & Schachar, 2009), psychiatric disorders (F.-F. Li, Chen, Zhang, Li, & Li, 2021; Lipszyc & Schachar, 2010; McTeague et al., 2017; Sjoerds, Van Den Brink, Beekman, Penninx, & Veltman, 2014; Yu et al., 2022), under the effect of medication (Kalanthroff et al., 2017; Sarkar et al., 2020) and intervention (Cai, George, Verbruggen, Chambers, & Aron, 2012; N. Swann et al., 2011; Wessel, Diesburg, Chalkley, & Greenlee, 2022), substance dependence (C.-s. R. Li et al., 2008; C.-s. R. Li, Milivojevic, Kemp, Hong, & Sinha, 2006; Smith et al., 2014), sleep disorders (Covassin et al., 2011; Zhao et al., 2018), learning difficulties (De Weerdt, Desoete, & Roeyers, 2013; Schmid, Labuhn, & Hasselhorn, 2011), eating disorders (Lavagnino et al., 2016; Svaldi, Naumann, Trentowska, & Schmitz, 2014), in studies of pregnancy related changes (Fiterman & Raz, 2019), and genetic basis of inhibitory control (Barnes, Dean, Nandam, O’Connell, & Bellgrove, 2011).

Understanding the dynamic cognitive mechanisms that underlie SST holds great promise for enhancing our knowledge of latent processes driving cognitive functioning. Despite its widespread use however, the SST and the traditional computational models applied to interpret it face significant challenges that limit their explanatory power and practical utility.

The most common approaches measure the efficiency of an individual’s inhibitory control by estimating the latent stop-signal reaction time (SSRT). SSRT cannot be measured directly and is typically estimated using a theoretical race model (Logan & Cowan, 1984; Logan, Van Zandt, Verbruggen, & Wagenmakers, 2014; Verbruggen & Logan, 2009a). Conventional methods for estimating the SSRT are the mean method and the integration method. In the mean method (mSSRT), the mean inhibition function or probability of responding given a stop signal during an SSD (for details, see Logan and Cowan (1984); Verbruggen and Logan (2009a)) is subtracted from the mean RT distribution. In the integration method (iSSRT), the completion of the stop process is estimated by integrating the RT distribution and determining the point at which the integral equals the probability of responding for a specific delay (Logan and Cowan (1984); Verbruggen and Logan (2009a)). The mean or integrated SSRT are the primary non-parametric methods used to index cognitive control, and the most popular measures used. Other measures include the slope of the inhibition function (Verbruggen & Logan, 2009a), or bespoke modelbased measures of SSRT.

### Need for a comprehensive integrative theoretically-motivated modeling framework

The need for a comprehensive, integrated, theoretically-motivated cognitive model stems from **(i)** Traditional models and related measures of SSRT being inappropriate for a significant proportion of individuals (**Figure 2**, green); **(ii)** Need for explanatory mechanisms – greater cognitive process dissociation and mechanistic explanations of behavior (**Figure 2**, blue). Figure 2 is based on the ABCD study where we examined 7,787 children who completed two runs of the SST task inside an fMRI scanner at baseline.

**Figure 2.**
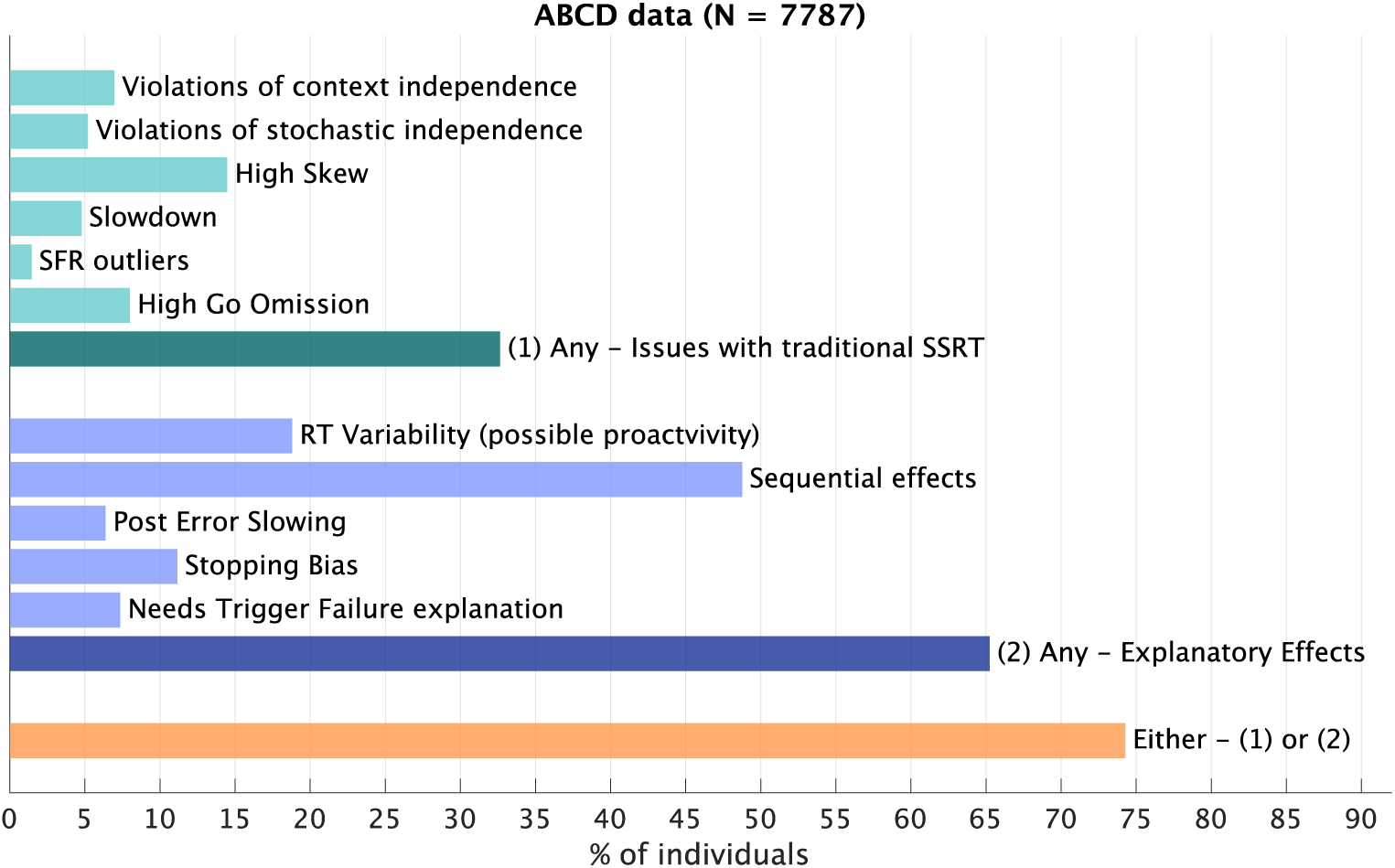
Proportion of participants for whom traditional SSRT measure may not be suitable (green) and for whom models with additional explanatory mechanisms may be required (blue).

### Traditional models and related measures of SSRT are not adequate or appropriate for a significant proportion of individuals

The validity of inferred SSRT measures become more questionable in the presence of specific behavioral patterns that confound SSRT measurement (R. A. Doekemeijer, Dewulf, Verbruggen, & Boehler, 2023; Verbruggen, Chambers, & Logan, 2013; Weigard, Heathcote, Matzke, & Huang-Pollock, 2019), potentially leading to systematic biases in conventional measures of SSRT and inhibitory control. There is a consensus that traditional and non-parametric (such as mean and integration methods) measures of SSRT should not be calculated or used when the assumptions of the conventional race model are violated, or when certain patterns of atypical or extreme behavior is observed. Recent research (Bissett, 2014; Bissett, Hagen, Jones, & Poldrack, 2021; Bissett, Jones, Poldrack, & Logan, 2021) has highlighted the severe implications of these violations, questioning the use of non-parametric SSRT as a definitive measure of inhibitory control (Bissett, Jones, et al., 2021; Verbruggen et al., 2013), although recent approaches have proposed model-based solutions to overcome some of these issues (Weigard, Matzke, Tanis, & Heathcote, 2021a, 2023). When non-parametric methods are used, the integration method (iSSRT) is considered more reliable and less biased than the mean method (Frederick Verbruggen et al., 2019). However, even iSSRT has been shown to have biases and underestimate true SSRT under multiple conditions.

### Violations of context or stochastic independence

Context independence refers to the assumption that go process reaction time distributions do not depend on whether or not a stop signal is observed, an assumption that is critical to non-parametric estimates of the SSRT. This assumption however has been shown to be consistently violated in a variety of stop signal tasks (Bissett, Jones, et al., 2021), including in the ABCD study (Bissett, Hagen, et al., 2021), where 7% of the participants show violations of context independence, as benchmarked by having stop RTs that are longer than go RTs (Frederick Verbruggen et al., 2019). Analysis of the ABCD data has shown that one important dimension of context dependence of the go process is the duration of the go stimulus, which varies significantly between trials, and also between go and stop trials on an average (Bissett, Hagen, et al., 2021), with stop trials having shorter stimulus display durations because of the onset of SSD. Bissett, Hagen, et al. (2021) recommend that parametric cognitive models of the stop signal task incorporate the sensitivity to stimulus duration into the go process. One such implementation proposed by (Weigard et al., 2021a), targets stop trials, such that the SSD affects the go process drift rate in a linear manner, until asymptotic performance is reached on par with the drift rate estimated on go trials. This addresses one of the issues raised, in terms of the large difference in stimulus presentation duration between stop and go trials but may not directly account for variability in stimulus duration within go trials.

Stochastic independence refers to the assumption that the finishing times of go and stop processes are independent on any trial, and violations of this assumption have been shown to lead to biases in the accurate estimation of SSRT (Frederick Verbruggen et al., 2019). Violations of stochastic independence might show up as high correlations between go process reaction times and SSRT times. In terms of model-free behavioral assessments, this has been shown to affect the slope of the inhibition function, that is, the rate of change in stop failure rate with increasing stop signal delays. We arrive at a conservative model-free estimate that in the current study, 5.2% of the participants may exhibit violations of stochastic independence, based on benchmarking against previous simulation studies (**Methods**).

### Higher skew and slowdown of reaction times

Higher right skew of the go process reaction times has been shown to bias estimates of non-parametric SSRT (Frederick Verbruggen et al., 2019; Verbruggen et al., 2013). Higher skew could be a sign of strategic or proactive slowing, or other sequential effects that produce greater variability in reaction times. In the current study, 14.5% of participants demonstrated higher levels of skew (**Methods**). Higher slowing of go process reaction times has also been shown to bias estimates of non-parametric SSRT (Frederick Verbruggen et al., 2019; Verbruggen et al., 2013). Higher slowdown could also be a sign of strategic or proactive slowing or other sequential effects. In the current study, 4.8% of participants demonstrated higher levels of RT slowdown (**Methods**).

### Deviations in stop success rates and go-omission rates

Higher go omission rates have been shown to affect accurate estimation of non-parametric SSRT (Frederick Verbruggen et al., 2019). Higher go omission rates may reflect some form of anticipatory preparation for inhibition before the observation of a stop stimulus. In the current study, 8% of participants demonstrated higher levels of go-omission rates (> 20%). Conventional non-parametric models also cannot capture patterns of inhibition functions that vary with SSD (Bissett, Hagen, et al., 2021), and SSRT estimates have also been shown to be unreliable when response inhibition (stop success rate) deviates sharply from 0.5, especially with higher stop success rates (Frederick Verbruggen et al., 2019). In the current study, 1.5% of participants demonstrated high deviations (SFR < 25% or > 75%).

### Need for explanatory effects – greater process dissociation and mechanistic explanations of behavior

Beyond just the ability to accurately measure SSRT for a diverse population with individual differences, a large body of work has shown the influence of multiple cognitive processes on inhibitory control, including proactive and strategic adjustments, sequential effects including error-monitoring and related adjustments, and attentional mechanisms such as trigger failure of the stop process. No existing model captures and measures all of these theoretically motivated aspects in a single quantitative framework that can provide a mechanistic explanation of how these components interact, and provide a measurement model for distinct cognitive process. The most important of these is the need for a cognitive model of the proactive inhibitory mechanisms.

Most previous measurements of proactive inhibition have relied on bespoke experimental manipulation, rather than a cognitive modeling approach that can be applied to standardized experimental paradigms.

#### Ability to identify, measure, and explain proactive mechanisms

Existing models of inhibitory control often fail to adequately differentiate between proactive and reactive control mechanisms. Proactive control refers to the anticipation and prevention of impulsive actions through the maintenance of goal-relevant information, whereas reactive control involves the suppression of an action in response to a stop signal (A. R. Aron, 2011; T. S. Braver, 2012; Cai, Oldenkamp, & Aron, 2011). The dichotomy between these processes and their interaction with both top-down and bottom-up regulatory mechanisms remains insufficiently explored in current research paradigms, especially in terms of quantitative measurements of such interacting processes (Adam R Aron, 2011; Elchlepp, Lavric, Chambers, & Verbruggen, 2016). In the current study, 18.9% participants demonstrated exceptionally high levels of variability in RT (**Methods**), suggesting a possibility for mechanisms that modulated within-individual RT.

Proactive inhibitory control has been assessed in terms of response slowing and higher variability (higher SD of RT) of the go process in blocks or cued designs when a stop signal is present compared to those that have no stop signals. However, such assessment requires a modified experimental setup and cannot be easily or directly assessed in a traditional stop signal task. Previous studies have modified the standard SST paradigm to probe the proactive component (Weidong Cai et al., 2023; Chikazoe et al., 2009; N. C. Swann et al., 2012; Verbruggen & Logan, 2009b; Vink, Kaldewaij, Zandbelt, Pas, & du Plessis, 2015), introducing variability in task design, and rendering replication more challenging. Such experimental setups have shown that a simple diffusion model fit of the reaction times resulted in an increase in the non-decision times for blocks that contain stop-signals (Verbruggen & Logan, 2009b). This has been attributed to either delayed-processing, where the processing of the go stimulus may be delayed, or response-suppression, where motor output is temporarily suppressed. Existing data are often compatible with both hypotheses. It might be expected that the ‘excess’ non-decision time inferred in such situations is independent of SSD and relatively stable in the latter hypothesis, but shows greater variability and depends on some learning-based expectancy of SSD in the delayed-response hypothesis. This need for SSD tracking of the delayed response hypothesis has led to some suggestions that it may be cognitively difficult to implement. It has however been suggested that SSD values could affect general strategy shifts in the stop signal task (Bissett, Hagen, et al., 2021), and that SSD and expected probability of stopping can affect reaction times (Ma & Yu, 2015). Some recent work (Elchlepp et al., 2016; Hu & Li, 2012; Perri, 2020) has also suggested that these response focused views should be expanded to include the role of dynamic attention in proactive control, including trial by trial modulation of attention and response settings, and a general attentional inhibitory account. This is also supported by evidence (Lee & Kang, 2020) that even proactive inhibition may employ distinct mechanisms that dissociate influence on slower reaction times and lower erroneous responses, suggesting that a simple adjusted-response threshold based explanation of proactive inhibition (Elchlepp et al., 2016) may not be sufficient. In addition, there is mixed evidence that proactive inhibition may support better reactive inhibition (Castro-Meneses, Johnson, & Sowman, 2015), suggesting some form of interaction between these processes.

Existing accumulator based cognitive models of the stop signal paradigm do not capture the possibility of variable SSD dependent delayed processing contributing to proactive inhibitory control or attention modulation (beyond rates of trigger failure, which are SSD independent), although there have been models proposed that capture similar dynamic belief effects within a Bayesian optimal control model (Ma & Yu, 2015). Such dynamic belief models have been used to study proactive control in the SST (Cai, Chen, Ide, Li, & Menon, 2017; Ide, Shenoy, Angela, & Chiang-Shan, 2013), but do not simultaneously incorporate other latent components, such as reactive control (Shenoy & Yu, 2011). They do not incorporate an evidence accumulation mechanism and have not typically been used as measurement models. They have been used primarily to showcase the influence of dynamic experimental contingencies (such as probability of stopping and SSD) on the variability of reaction times across subjects, without modeling any individual level parameterization that can capture or measure individual differences. Moreover, this class of models assumes an optimal Bayesian process, and are not ideal to model impaired processes or variations in clinical populations. This theoretical gap hinders a comprehensive understanding of the multiple dimensions of inhibitory control and their implications for behavior and cognition. There is thus, a need for cognitive models that capture processes in an integrated manner, the effect of SSD tracking on inhibitory modulation, possible mechanisms of proactive inhibition in the form of delayed response or response suppression, and the modulatory effects of preemptive mechanisms on reactive inhibition.

#### Sequential effects affecting the go process

We measured whether the reaction times or go-stop errors differed between trials that followed a correct or incorrect trial (incorrect trials included go omissions or stop failures). 48.8% of participants showed significant differences in at least one of these measures, and 33.8% showed significant differences specifically in reaction times, demonstrating the influence of error-based sequential effects on the go process. This sequential effect demonstrates another form of stochastic dependence, and motivates the need for mechanisms that explain systemic patterns of intra-individual variability without assuming that the go process is independent and identically distributed (IID) across trials. In addition, response thresholds have been shown to increase post errors and result in post error slowing in a variety of tasks (Schroder et al., 2020), but these mechanisms are not usually incorporated into cognitive models of the stop signal task. 6.4% of participants in this study demonstrated statistically significant post-error slowing.

#### Sequential effects affecting the stop process

Apart from the go process, stopping processes may also not be context independent, conditional stop trial probabilities may influence behavior, and SSRT may depend on SSD (Bissett, Hagen, et al., 2021). These aspects remain unaccounted for in most known cognitive models of the stop signal task that employ evidence accumulation models. Modulation of the conditional probability of stopping has been incorporated in some other models (Ma & Yu, 2015) but not explicitly as attentional mechanisms that depend on the number of trials that have elapsed since the last stop signal was encountered. 11.2% of participants showed a significant correlation between the number of trials since a stop signal was last observed, and the stop failure rate, demonstrating some sequential attentional mechanisms affecting dynamic modulation of the stop process.

Given the issues with non-parametric estimation of SSRT, alternate methods for estimation of SSRT are parametric models, which may be cognitive, statistical, or hybrid models that model the decision-making process within the SST paradigm. Some of these models are based on assumptions of a parametric distributional form of the SSRTs (Matzke et al., 2013; Weigard et al., 2019; Weigard, Matzke, Tanis, & Heathcote, 2021b). Others are based on the assumption of competing evidence accumulator processes reflecting plausible cognitive mechanisms (Heathcote & Love, 2012; Logan et al., 2014). However, the stop signal evidence accumulation process within such models has been shown to have poor identifiability and thus not be a suitable measurement model (Matzke, Logan, & Heathcote, 2020a). As an alternative, hybrid models have been suggested where the go process is modeled by a cognitive evidence accumulation process, and the stop process characterized by a parametric distribution such as the ex-Gaussian distribution (Weigard et al., 2021b). While such methods have produced more reliable and useful estimates of SSRT, they may not be the most suitable approach for accounting for how an integrated set of cognitive processes such as belief updating, error-based learning and modulation, and adjustment of attentional and response parameters affects the stop process in a dynamic manner.

#### Mechanisms underlying trigger-failures of the stop process

In previous literature it has been empirically observed that stop failure rate sometimes does not approach zero even on extremely short SSDs close to zero, and this has typically been explained with the assumption of trigger failures (R. Doekemeijer, Verbruggen, & Boehler, 2021), that is, a failure of the stopping process to initiate on a small subset of stop trials. Without the concept of trigger failures, it would be required that a stopping process that generates short enough SSRTs to work effectively at higher SSDs would also produce extremely long SSRTs that fail to successfully inhibit action at SSDs close to zero. Traditional models with fixed SSRTs, or even those that allow random variability in SSRT cannot produce the high level of trial-level variability required to consistently explain these findings at low and high SSDs. Some models (Matzke, Love, & Heathcote, 2017) have thus been augmented with a trigger-failure mechanism that assumes that the stop process fails to initiate on some percentage of stop trials. The inferred failures of inhibitory response processes have been associated with impulsivity (Skippen et al., 2019). It has also been proposed that there may be a confound between trigger failures and inference of longer SSRTs (Weigard et al., 2019). Allowing for a high-variability stopping process that can account for shorter and longer SSRTs, and thus provide an underlying systemic mechanism for when behavior that seems like a trigger-failure occurs, can provide a more powerful understanding of the dynamic mechanisms modulating the stopping process. In this study, 7.4% of the participants had stop failure rates in the lowest 10^th^ percentile (shortest or easiest SSDs) that were at least as high or higher than in the highest 10^th^ percentile (longest or hardest SSDs), representing participants for whom mechanisms similar to, or underlying trigger failures may be relevant.

## Summary

Typical recommendations (Bissett, Hagen, et al., 2021) to capture accurate inhibitory capacities based on the independent race model focus on excluding participants with extreme behaviors, or focus on experimental design that encourages standardized behavior – minimizing strategic adjustments, slowing, sequential effects, etc. These approaches reduce the elicitation of individual differences, and try to minimize the role of multiple cognitive and metacognitive systems that interact with core reactive inhibitory capabilities. Capturing individual differences in these multiple cognitive and metacognitive systems is critical for understanding behavior in ecologically relevant cognitive control situations. In the current study, using the above defined conservative benchmarks, at least 32.6% (**Figure 2**; green) participants show at least one of the behaviors outlined above that will affect accurate estimation of non-parametric SSRT.

Additionally, at least 65.2% (**Figure 2**; blue) of participants demonstrate the need for at least one of the dynamic measures of individual differences outlined above – in proactive mechanisms or sequential dependencies in go or stop processes, that interact with and complement basic reactive inhibitory control. Overall, 74.3% of participants fall under at least one of the above two categories. Note that some of these estimates are made based on subjective criteria (**Methods**). Nonetheless, it is not the specific proportions that are important, rather the fact that a substantial proportion of participants demonstrate these.

This critique underscores a critical gap in the current understanding and measurement of inhibitory mechanisms, pointing to the need for more quantitively precise models that can account for the complex dynamics of cognitive processes. To address this, we develop, implement, and validate a model of Proactive, Reactive, and Attentional Dynamics (PRAD), a novel computational cognitive model for inhibitory control that comprehensively accounts for the multi-componential dynamic processes that are not currently represented in extant models, and can address most of the above issues (see **SI Table S1** for a complete list of features).

### Proactive, Reactive, and Attentional Dynamics (PRAD) - An Integrated Approach

The PRAD model incorporates the dynamic modulation of behavior by multiple latent cognitive processes governing inhibitory control, allowing for multiple forms of sequential dependencies, both in the go and stop processes, and provides quantitative measurements for multiple cognitive components of reactive and proactive control. The dynamic modulation of behavior is based on theoretically motivated processes and includes dependencies on both endogenous (cognitively generated, such as being based on error-monitoring) and exogeneous (experimental contingencies, such as updating beliefs based on varying stop signal delays, or attentional modulation based on the number of trials since a stop signal was encountered) variables.

Importantly, it allows for measurement of individual differences in the cognitive variables that govern how these sequential dependencies are operationalized. PRAD does not require assumptions about context or stochastic independence, and it is not constrained by behavioral patterns such as high skewness of slowdown in reaction times, or extreme error rates that impact robust measurement of SSRT in traditional approaches. It provides a novel comprehensive account of dynamic reactive and proactive inhibition, providing more robust and dissociated measures of individual differences in inhibitory control. We demonstrate PRAD’s robustness across a wide range of measures and test its ability to overcome limitations of conventional race models.

One of our key objectives of implementing PRAD was to make theoretical advances in our understanding of cognitive control dynamics, by examining the distinct components of inhibitory control, including proactive delayed response (PDR) mechanisms (**Figure 3A**; green box) and attentional modulation of stopping (AMS) mechanisms (**Figure 3A**; yellow box), and their dynamic interactions, and by investigating systemic intra-individual variability in inhibitory control processes. A second key objective was to provide debiased estimates of SSRT, and debias inferences about cognitive control mechanisms especially for typically underrepresented or atypical populations. Since PRAD provides a multidimensional measurement system, this is expected to provide a stronger characterization of individual differences. Our third objective focused on evaluating the relevance of PRAD measures to a variety of cognitive task performance measures, its relevance to identifying differences in clinical subgroups, its ability to identify specific neural mechanisms underlying proactive and reactive control, and its relation to exposomic factors (Moore et al., 2022). These goals sought to validate PRAD’s effectiveness beyond its initial context and explore its potential as a more precise tool for understanding the latent substrates of cognitive variability. We leverage the large-scale, Adolescent Behavioral and Cognitive Development (ABCD) study data to develop and test the PRAD model in a developmentally homogenous (ages 9-10) group of children who completed the SST task.

**Figure 3.**
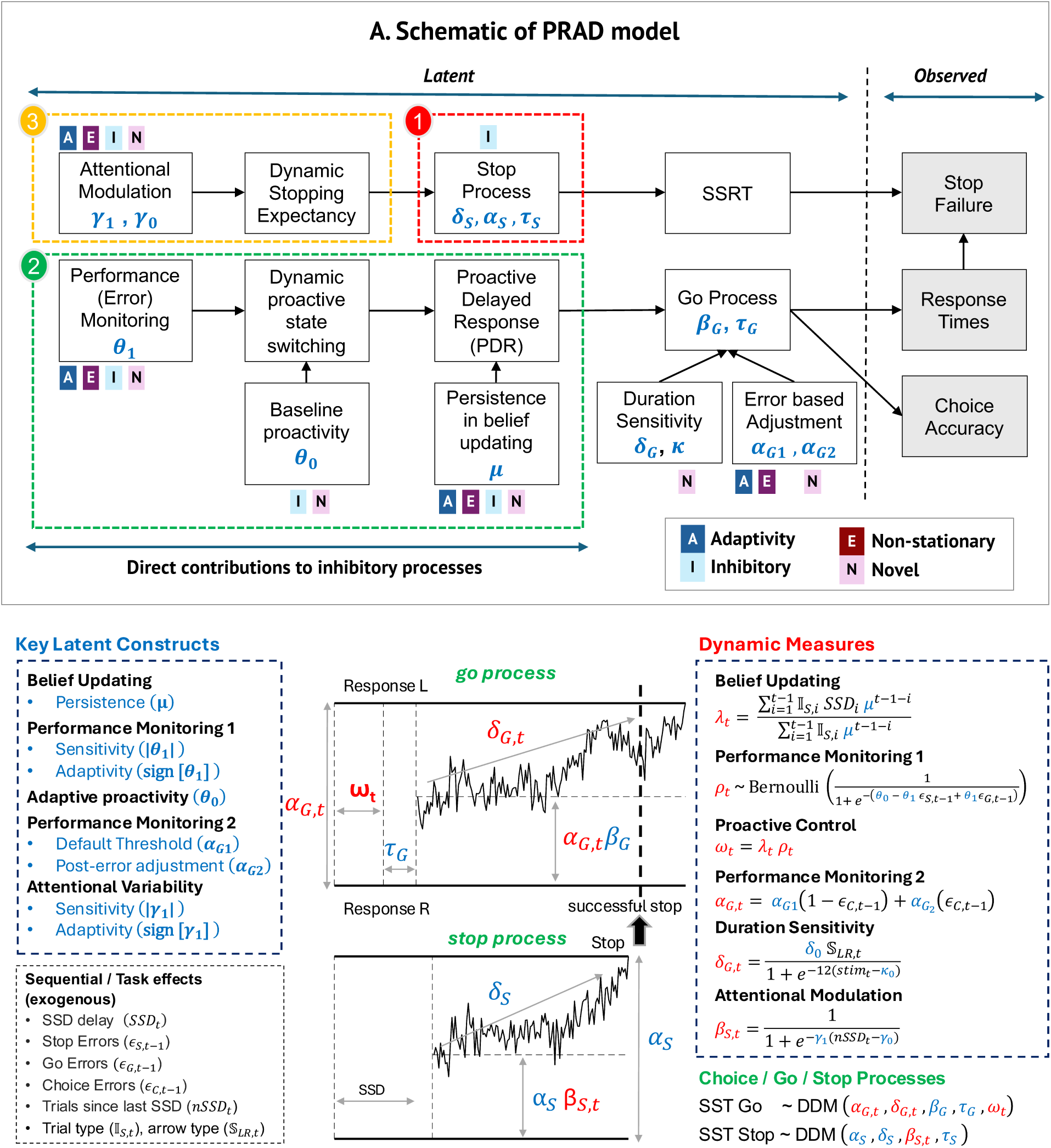
Schematic of the Proactive Reactive and Attentional Dynamics (PRAD) model. (A) Overview of how latent cognitive processes interact to produce observed behavior. The model infers latent variables related to three mechanisms of dynamic inhibitory control: basic reactive inhibition, proactive response delaying, and attentional modulation of stopping expectancy. (B) Mathematical details of the model, showing two drift diffusion processes (go, stop) augmented with dynamic parameters. Blue - individual-level parameters, Red - trial-level dynamic parameters.

### Specification of the PRAD model

The PRAD model was implemented within a hierarchical Bayesian framework, allowing for the estimation of both individual-level parameters and trial-level measures. This approach enables the identification and measurement of distinct components that characterize individual differences in inhibitory control, while also accounting for the temporal variability in cognitive processes. **Figure 3A** illustrates the model’s structure, incorporating proactive, reactive, and attentional mechanisms. **Figure 3B** and **SI Table S2** illustrate the mechanisms and summarize the key model parameters.

### Competing Go and Stop processes

Overall, the PRAD model incorporates separate evidence accumulation (drift-diffusion) processes for the go and stop processes, similar to a canonical horse-race model (Band, Van Der Molen, & Logan, 2003). However, in addition to typical driftdiffusion process parameters, PRAD includes additional individual trait-like and dynamic trial-level measures, which have been shown to improve model inferences in different cognitive tasks (Mistry, Chang, El-Said, & Menon, 2025; Mistry, Skewes, & Lee, 2018; Mistry & Trueblood, 2015).

### Dynamic modulation of the Go process (PDR)

The go process is modeled as a drift diffusion process, modulated by a probabilistic proactive delayed responding (PDR) mechanism, which in turn, is driven by a cognitive state switching mechanism and belief updating about stop signal delays. On proactive trials, the go process can be strategically delayed in anticipation of a possible stop signal. In PRAD, 𝜌_!_ is a binary variable representing the presence (𝜌_!_ = 1) or absence (𝜌_!_ = 0) of a proactive cognitive state on trial 𝑡. The proactive delayed responding is only initiated on proactive cognitive states. Proactive cognitive states are governed by a baseline proclivity for proactivity (𝜃_"_), and performance-monitoring based modulation (𝜃_#_).

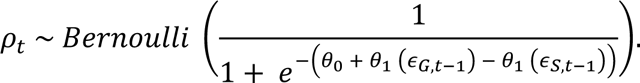

Here, 𝜖_-,!$#_ is an indicator of a go-omission (incorrectly stopping on a go trial) on the previous trial, and 𝜖_/,!$#_ is an indicator of a stopping error (not stopping on a stop trial). The PRAD model assumes that the correction in terms of increasing or decreasing the probability of a proactive cognitive state following these two types of trials will be in opposite directions. The sign of 𝜃_#_ is an indicator of adaptivity or maladaptivity of the performance monitoring mechanism – when 𝜃_#_ is negative, 𝜌_!_ adaptively increases after a stop-failure but reduces after a go-omission, and when 𝜃_#_ is positive, 𝜌_!_ maladaptively decreases after a stop-failure but increases after a go-omission. The absolute value of 𝜃_#_ is the sensitivity of the state-switching mechanism to errors and governs how large of an effect the errors have on the trial-level probability of proactivity. The probability of proactivity in the model is the posterior mean of 𝜌_!_, i.e., the posterior probability of 𝜌_!_ = 1.

When in a proactive cognitive state, the onset of the go process may be deliberately delayed in anticipation of a stop signal. This dynamic adaptation is modeled by adding a further delay 𝜔_!_ to the go process to reflect proactive delayed responding to the go stimulus, where:

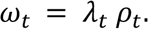

Here, 𝜆_!_ reflects a trial-level belief updating process, based on the history of stop signal delays (SSD) encountered, and is an internal noisy estimate of the prospective anticipated SSD. The parameter 𝜇 (0 < 𝜇 < 1) reflects persistence in belief updating, with high persistence implying a lower decay rate of older SSDs encountered. Further, letting SSD_!_ be the SSD on trial *t* and 𝕀_/,!_ be a stop trial indicator on trial *t*, the belief updating can be written as:

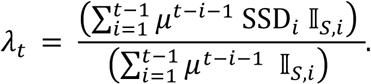

Proactive delaying (𝜔_!_ ) is the posterior mean of 𝜆_!_ on trials with probability of proactivity greater than 0.5, otherwise 0. Thus, the overall effective non-decision time, as compared to traditional models, will be:

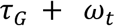

Here, 𝜏_-_ is the trial-invariant or core non-decision time component, while 𝜔_!_ measures the strategic adjustment to the non-decision time. Note that because of this mechanism, 𝜏_-_ cannot directly be compared to non-decision times from traditional models (see **Methods** for details).

In addition to the PDR mechanism, the go process incorporates a trial-varying decision threshold (𝛼_-,!_) and drift rate (𝛿_-,!_). The dynamic decision threshold is based on post-error adjustments:

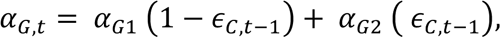

where (𝛼_-#_, 𝛼_-3_) are distinct threshold levels and 𝜖_2,!$#_ is an indicator of whether the left vs right choice on the previous trial was erroneous. Thus, the dynamic threshold implements a form of performance monitoring and varies between two levels based on the outcome of the previous trials, with 𝛼_-#_ being the default threshold and 𝛼_-3_ reflecting the threshold after post-error adjustments.

The dynamic drift rate is specified as:

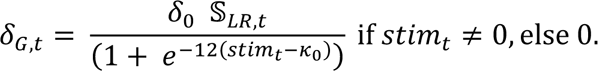

Here 𝛿_"_ is a measure of the maximum drift rate for an individual, with the actual drift rate depending on the duration of the go stimulus (𝑠𝑡𝑖𝑚_!_) and an individual parameter 𝜅_"_, which can be interpreted as the stimulus duration at which the drift rate is half the maximum. 𝕊_45,!_ assumes values 1 or -1 depending on the direction of the go stimulus (left or right). PRAD allows for trial level changes to the drift rate, overcoming the issues with variable go stimulus durations highlighted in previous work (Bissett, Hagen, et al., 2021), especially on stop trials where the go stimulus is overridden by the stop signal. Since the duration of the go-stimulus on go trials is confounded with the RT, making this a discriminative rather than generative model, we also test a fully generative model (PRAD-g; see **Methods** for details), where the dynamic drift rate is only applied on stop trials, and on go trials, it is retained as 𝛿_"_ 𝕊_45,!_, similar to previous work (Weigard et al., 2023).

The reaction times of the go process correspond to the reaction times for pressing the left or right buttons in response to the go stimulus.

### Dynamic modulation of the Stop process (AMS)

The stop process combines baseline reactive inhibition with attentional modulation of stopping (AMS). It is modeled as a drift diffusion process with a trial-invariant non-decision time (𝜏_/_), decision threshold (𝛼_/_), and drift rate (𝛿_/_), but a trial-varying bias (𝛽_/,!_). The stop process begins at the onset of the stop signal. The initial bias is

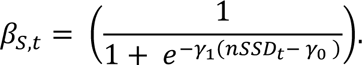

Here, 𝑛𝑆𝑆𝐷_!_ reflects the number of trials since a stop signal was last encountered. This reflects an attentional mechanism that modulates the stopping bias 𝛽_/,!_ (which varies from 0 to 1). Positive values of 𝛾_#_ result in an increase in stopping bias as 𝑛𝑆𝑆𝐷_!_increases and vice versa. Similarly, negative values of 𝛾_#_ result in a decrease in stopping bias as 𝑛𝑆𝑆𝐷_!_increases and vice versa. The absolute value of 𝛾_#_ measures the sensitivity to attentional modulation. The 𝛾_"_ parameter is a measure of the value of 𝑛𝑆𝑆𝐷_!_ when stopping bias is neutral (0.5).

The reaction times of the stop process correspond to the SSRT. The stop process is only initiated on stop trials after the appearance of the stop stimulus (which appears after a delay corresponding to the SSD). The PRAD model thus allows SSRT distributions to be dependent on the stopping bias (conditional probability of stopping), which are in turn inferred to be modulated based on attentional sensitivity to the number of trials that have elapsed since the last stop signal was encountered. The SSRT is not manifested as a behavioral action. Rather, if the SSRT, which is the duration of the stop process, plus the SSD on a stop trial is smaller than the go process reaction time, then the go action can be successfully inhibited (successful stop). The PRAD model enables obtaining the full posterior distributions of SSRT (Matzke et al., 2013), at a trial level, rather than a single individual level estimate obtained using non-parametric methods.

### Model implementation

We implemented PRAD within a hierarchical Bayesian modeling framework in JAGS (Plummer, 2003), which implements a Gibbs sampler for Markov Chain Monte Carlo (MCMC) simulations. Both the go and stop processes are modeled using the Wiener distribution (Wabersich & Vandekerckhove, 2014), which produces a joint distribution of the reaction times and the decision choice on each trial. The sampling hyperparameters, Bayesian priors, parameter recovery (**SI Figure S1**), and additional computational details are described in the **Methods** section. We applied PRAD to SST data from 7787 individuals from the ABCD dataset.

### Control Models and Sensitivity Analysis

We also compared PRAD versus two control models – fixed stopping model (FSM) and random variability model (RVM). Both of these include competing go and stop processes, but FSM does not incorporate any trial-level variability in SSRT, while RVM allows random variability in SSRT across trials. Neither incorporate PDR or AMS mechanisms. In addition, we conducted a sensitivity analysis with slightly different implementations of the PRAD model to test robustness to some modeling assumptions (PRAD-n, PRAD-g, as well as subsampling based on choice of MCMC hyperparameters). Details are provided in the **Methods**.

### Model Validation - Explaining Aggregate Behavioral Patterns

PRAD demonstrated robust performance across various measures and outperformed control models (**Figure 4A-F; SI Figure S2)**. **SI Table S3** shows key observed measures at an aggregate level, and the summarized PRAD model posterior values corresponding to these observed measures.

**Figure 4.**
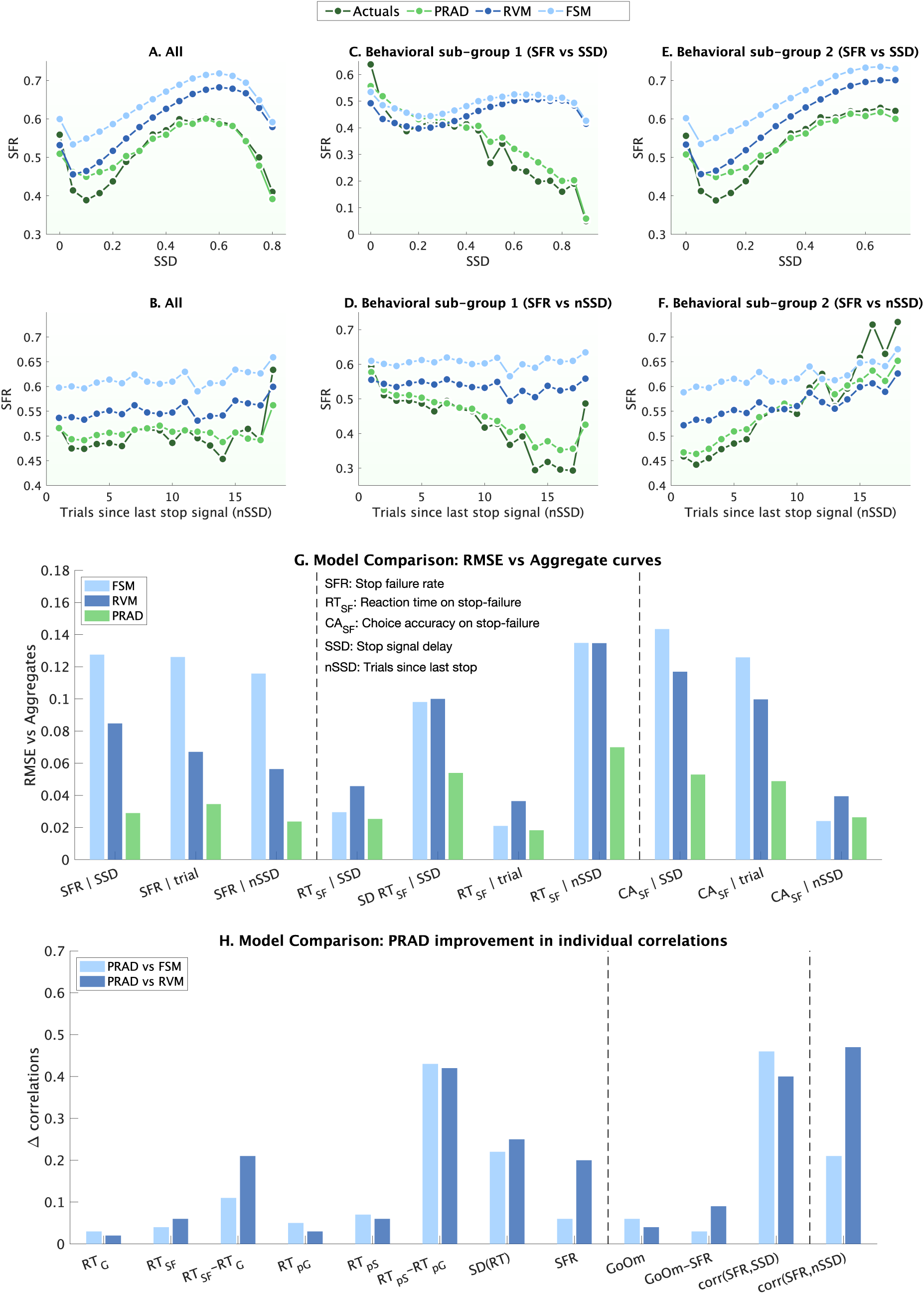
Model fit comparisons. (A-F) Aggregate behavior patterns for stop-failure rates (SFR) across different stop-signal delays (SSD) and number of trials since last stop signal (nSSD), comparing actual data with predictions from PRAD and control models (RVM and FSM). PRAD uniquely accounts for subgroup differences in SFR-SSD and SFR-nSSD relationships. (G) Root mean square error (RMSE) comparisons across models for various behavioral measures, showing PRAD’s superior fit. RMSE computed based on the distance between binned aggregate curves (actuals vs model posterior generated values). (H) Improvements in individual-level fit correlations using PRAD compared to control models across multiple behavioral measures. Dark green – actuals, Light green – PRAD, dark blue – RVM, light blue – FSM. RVM = Random Variability control model; FSM = Fixed SSRT control model.

At the group level, PRAD accurately captured trends in stop failure rates and reaction times across different experimental contingencies (SSD, nSSD). Specifically, the posterior predictives generated by the PRAD model explained aggregate behavioral patterns including non-linear Sshaped patterns of stop failure rate with increasing SSD (**Figure 4A**) and non-monotonicity at low SSD values, an indicator of violations of context independence (Bissett, Hagen, et al., 2021). Importantly, unlike control models, PRAD effectively captured the diverse patterns across different behavioral subgroups based on simple observed measures, such as whether SFR increased or decreased with SSD (**Figures 4C,E**). PRAD captured the increasing variability in stop failure rate with the numbers of trials since encountering the last stop signal (nSSD), with superior model fits for subgroup showing increasing and decreasing trends, both of which the control models struggled to fit well (**Figure 4B,D,F**). This demonstrates the fewer inductive biases within the PRAD model. PRAD also captured the linear increase in stop-failure RT with increasing SSD (**SI Figure S2A**), an indicator of the link between slower RT and better stopping performance, or the influence of SSD on RT, the increasing variability in RT with SSD (**SI Figure S2B**), and lower choice accuracies at low SSDs (**SI Figure S2C**).

Crucially, the PRAD model outperformed both control models (FSM, RVM) on all of these aggregate measures. RMSE distance from the observed aggregate curves (**Figure 4G, SI Table S4**) show that across trends based on different experimental contingencies, PRAD reduced RMSE of model fits by between 77% - 80% for SFR, 13% - 45% for RT on stop failures, and 48% - 61% for choice accuracies on stop failure, compared to the FSM model. Similarly, PRAD reduced RMSE by between 58% - 66% for SFR, 45% - 50% for RT on stop failures, and 48% - 51% for choice accuracies on stop failure, compared to the RVM model. These results suggest that incorporating sequential adaptive processes is crucial for characterizing behavioral dynamics on the SST, which PRAD achieves better than conventional approaches.

### Model Validation – Capturing individual differences and individual level behavior

We then examined PRAD robustness at the individual participant level. We assessed how well the model posterior values generated by PRAD explain behavioral patterns at an individual level (mean values per individual). Individual subject-level fits showed strong correlations between PRAD model fits and observed data for multiple behavioral measures (**SI Figures S3**). Notably, individual-level comparisons showed stronger correlations (**Figure 4H; SI Table S5)** and lower RMSE (**SI Table S6)** between observed and predicted values for PRAD across almost all key measures compared to the RVM and FSM control models. This included go reaction time (r = 0.91; p < 0.0001), stop-failure reaction time (r = 0.90; p < 0.0001), and stop-failure rate (r = 0.85; p < 0.0001), but importantly, also second order effects like post-go RTs (r = 0.92, p < 0.0001), post-stop RTs (r = 0.86, p < 0.0001), difference between stop and go RTs (r = 0.51, p < 0.0001), post-inhibitory differences in RTs (r = 0.45, p < 0.0001), and dynamic within-subject correlations like SFR vs SSD (r = 0.63, p < 0.0001) and SFR vs nSSD (r = 0.73, p < 0.0001).

PRAD’s ability to account for the wide range of variability and individual differences is also reflected in the Kullback-Leibler divergence between the distribution of observed measures and model fit posterior values of these measures. PRAD reduces the KL divergence (thus providing a closer match to the observed range of individual differences) for observed distributions by between 29%-89% compared to RVM and 4%-83% compared to FSM (**SI Table S7**). These results demonstrate that PRAD can accurately capture behavioral variability at both the individual-subject level and in terms of individual differences between subjects.

Additionally, we measured Deviance information criteria (DIC), which assesses model fit appropriately penalized for model complexity. In spite of the additional complexity, PRAD resulted in lowest (best) DIC values for 60% of the individuals, compared to 22% for the RVM, and 18% for the FSM model, suggesting that the additional complexity was necessary to explain behavior in a majority of individuals.

### Model Validation - Capturing intra-individual variability

For a significant proportion of participants, observed and latent measures show within-subject correlations with various sequential or experimental contingencies **(SI Table S8)** such as SSD, number of trials since the last stop signal was encountered (nSSD), the average SSD encountered so far (xSSD), whether a left versus right error was made on the previous trial (post_CE_), whether the previous trial was a successful go versus stop trial (post_GSE_), and whether the previous trial was an inhibitory trial (post_S_). 81% of the participants had a significant correlation of observed trial-level RT, 95% had a significant correlation of observed trial-level go-stop errors, and 82% had a significant correlation of latent trial-level SSRT with at least one or more of these exogeneous (SSD, nSSD, xSSD) or endogenous (post_CE_, post_GSE_, post_S_) variables. This analysis suggests that a good cognitive model should be able to at least partially capture structural sources of trial level intra-individual variability (IIV).

Analysis of IIV in Go RTs and SSRTs revealed that significant variance could be explained by model parameters, rather than random noise (**SI Figures S4**). See **SI Table S9** for summary statistics for PRAD latent dynamic measures, their coefficient of variation, and within-individual 95% CI ranges. We measured the partitioned variance of RTCV and SSRTCV into model parameters that were theoretically relevant. This revealed that 72% of RTCV was explained by the relevant cognitive model parameters. Of this, 26% was explained by standard drift diffusion model parameters and interactions between them, 46% was explained by the novel adaptive model parameters or their interactions with standard parameters, with 28% being residual, attributed to unexplained IIV, and possibly the contribution of noisy cognitive processes to IIV. Similarly, the partitioned variance of SSRTCV into relevant cognitive model parameters revealed that 81% of SSRTCV was explainable. Of this, 40% was explained by standard drift diffusion model parameters and interactions between them, 41% was explained by the novel adaptive model parameters or their interactions with standard parameters, with 19% being residual. These results point to the fact that a large percent of IIV can be explained structurally by the cognitive model parameters, without resorting to random effects across trials.

### Examining Model Dynamics With An Example

To visualize the dynamic interplay of key PRAD components and their trial-by-trial variability at the individual subject level, we examined detailed time courses of model-derived measures for representative participants. **Figure 5** illustrates the dynamics involved in stop trials from a single participant, showing how stopping expectancy, SSRT, PDR, and RT interact on a trial-by-trial basis. Stopping expectancy, modulated by attentional regulation, fluctuated considerably from trial to trial and showed an inverse relationship with SSRT. This revealed how dynamic attentional processes can influence inhibitory performance on a moment-to-moment basis. We also observed that task difficulty (represented by SSD) and stopping efficiency (SSRT) varied substantially across trials, with their sum (SSD+SSRT) providing insight into the overall challenge of inhibition on each trial. Our analysis of stop-failure trials revealed complex relationships between PDR, SSD+SSRT, and RT, demonstrating how proactive and reactive mechanisms interact to determine inhibitory outcomes.

**Figure 5.**
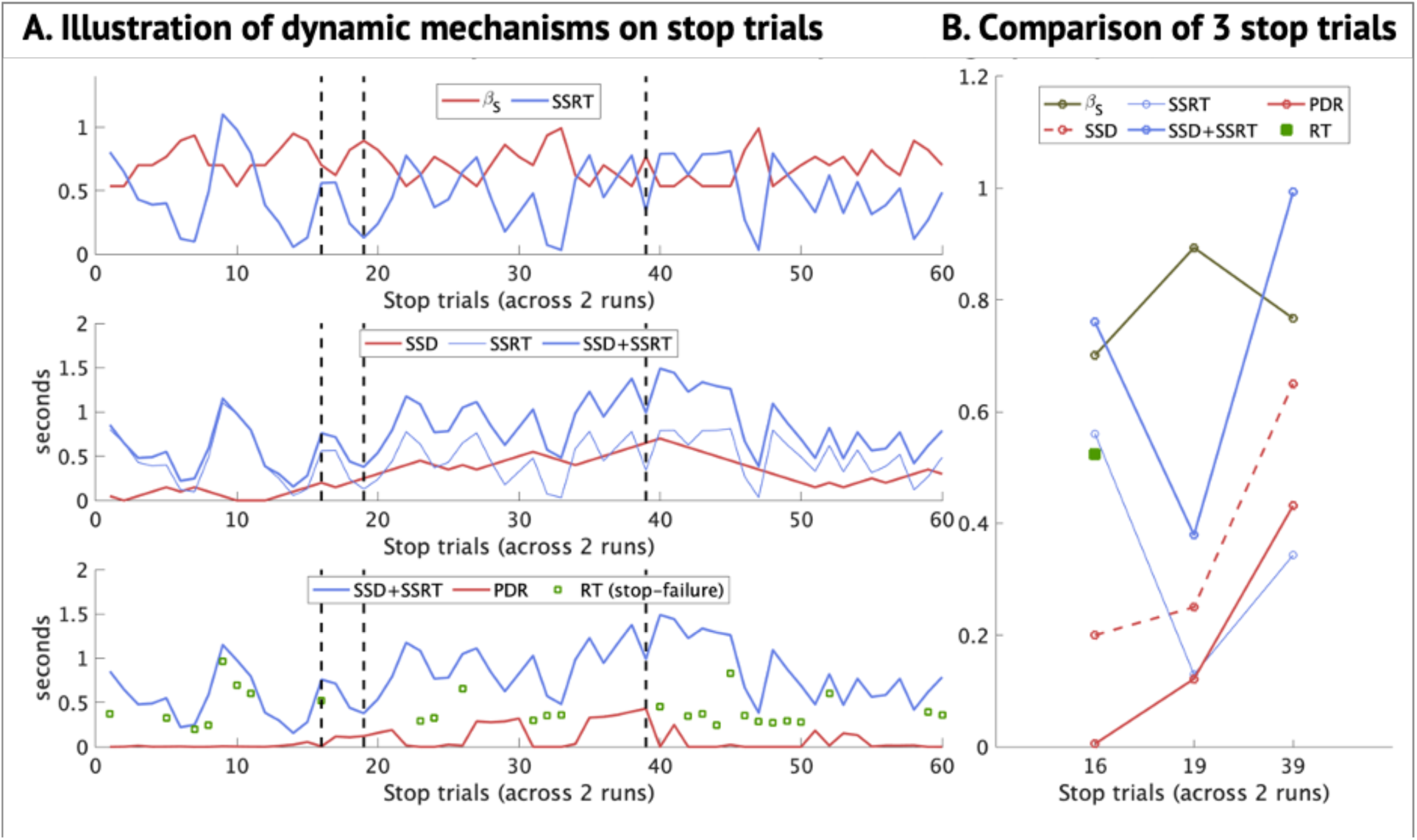
Illustrative single-participant dynamics on stop trials. (A) Trial-by-trial fluctuations in stopping expectancy, SSRT, SSD, PDR, and RT, illustrating the complex interplay of these variables. (B) Comparison of three specific stop trials (16, 19, and 39) demonstrating how different combinations of SSD, SSRT, and PDR lead to successful stops or failures, highlighting the model’s ability to capture nuanced trial-level dynamics. On stop trial 16, SSD is low (easy trial), but it still results in a stop-failure as there is no proactive delaying of the go process (PDR ∼ 0), and RT < (SSD+SSRT). On stop trial 19, SSD is higher (difficult trial), but SSRT is much lower because stopping expectancy is higher (attentional modulation of SSRT), resulting in a successful stop. On stop trial 39, SSD is even higher (very difficult), and stopping expectancy is not very high, resulting in a high SSRT. However, in spite of SSD+SSRT being higher than stop trial 16, this trial does not result in a stop-failure because of the influence of proactive delay of the go process (high PDR).

To further elucidate these dynamics, we closely examined three specific stop trials with varying levels of difficulty. We found that successful inhibition could occur even on more difficult trials (higher SSD) when compensatory mechanisms like increased stopping expectancy or heightened PDR were engaged. Conversely, easier trials could result in failures when these compensatory mechanisms were absent.

These observations highlight PRAD’s capacity to capture and explain the substantial intraindividual variability in inhibitory control processes, accounting for the complex interplay between task parameters, attentional modulation, and proactive control strategies that occur on a trial-by-trial basis.

### PRAD Advances Theoretical Perspectives of Cognitive Control

#### Distinct cognitively plausible components of inhibitory control

The model envisages a core reactive inhibitory process, modulated by attentional variations in stopping expectancies, and a dynamically adjusted proactive delayed response of the go process, all three of which affect observed inhibitory behavior (**Figure 6A**). Factor analysis of a subset of model parameters that are theoretically relevant to these three aspects of inhibitory control revealed the robustness of this three-factor structure (CFI 0.997, TLI 0.977, RMSEA 0.045; **Figure 6B, SI Table S10**). Factor 1, representing proactive control, loaded heavily on parameters (𝜃_"_, 𝜃_#_, 𝜇) that governed the trial level probability of engaging PDR mechanisms, and the adaptive belief updating about historical stop-signal delays that modulated the PDR duration. Factor 2, capturing basic reactive control (baseline SSRT), was dominated by parameters (δₛ, αₛ) governing the baseline stop process. Factor 3, reflecting attentional modulation that influences variations in trial-level SSRT, loaded strongly on parameters (𝛾_"_, 𝛾_#_) influencing the dynamic adjustment of stopping expectancy, which in turn affects dynamic SSRT. Factors 1 and 3 were not correlated, but factor 2 shows low correlations with the other 2 factors (|r| = 0.085, 0.16; both p < 0.0001), suggesting related but distinct processes. **SI Table S11** reports control analysis showing that 1- and 2-factor models were not adequate, and a 4-factor model was not identifiable.

**Figure 6.**
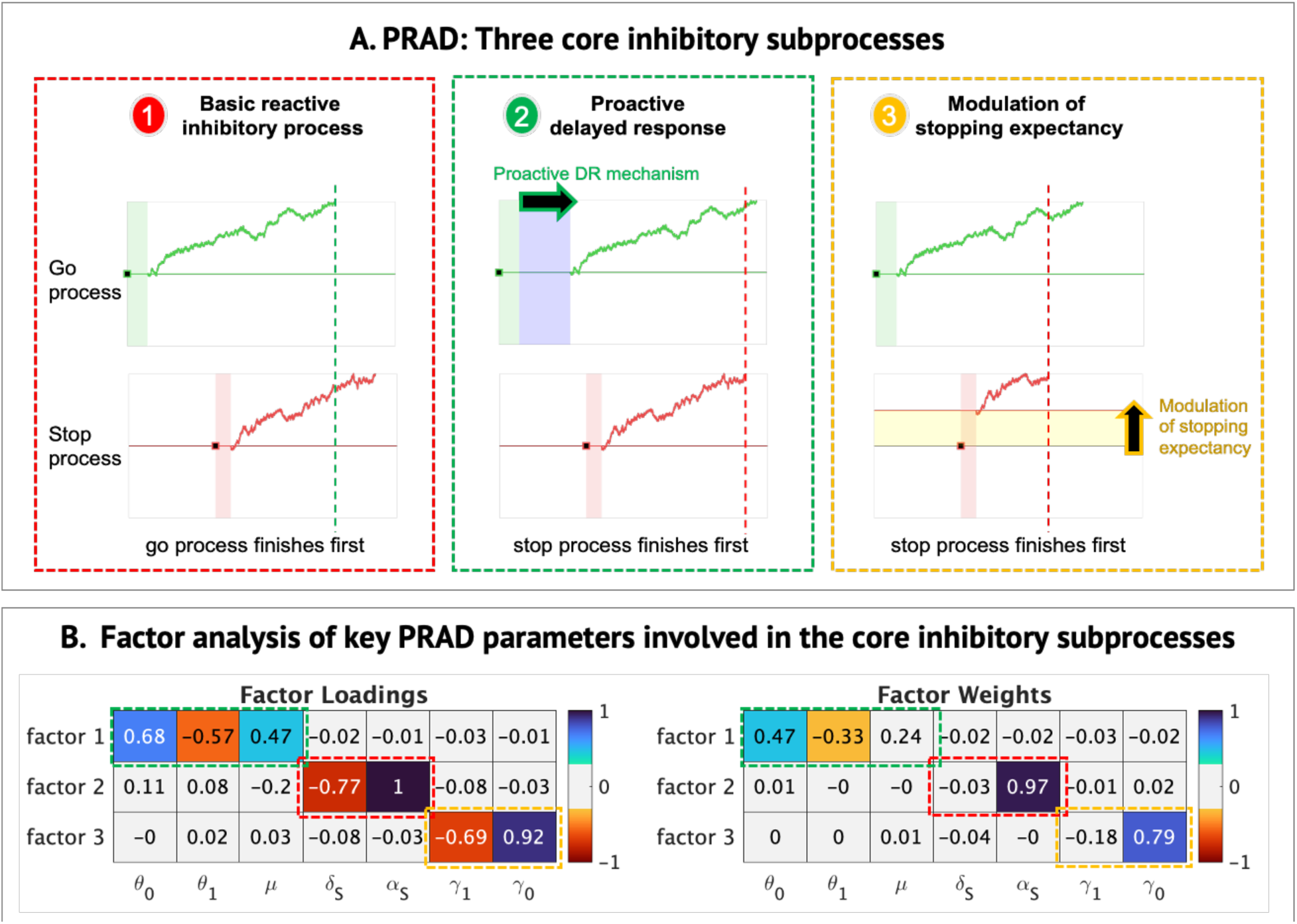
Core components of the PRAD model. (A) Illustration of three key inhibitory subprocesses: (1) basic reactive inhibition (red box), where competing Stop and Go processes determine trial outcome; (2) proactive delayed responding (PDR, green box), where anticipatory delaying of the Go process can lead to successful stopping; and (3) attentional modulation of stopping expectancy (yellow box), where top-down modulation can influence SSRT. (B) Factor analysis results showing how seven key PRAD parameters load onto three distinct factors corresponding to these subprocesses – proactive control, basic reactive control, and attentional modulation – demonstrating the model’s ability to dissociate different aspects of inhibitory control.

The three factors explained a significant portion of individual differences in inhibitory control performance. Regression analyses using all individual-level model parameters significantly predicted various measures of inhibitory control, including PRAD stop-signal reaction time (SSRT; 𝑅^3^ = 86%), observed stop-failure rate (SFR; 𝑅^3^ = 65%), observed mean experienced stop-signal delay (xSSD; 𝑅^3^ = 77%), SSRT coefficient of variation (SSRTCV; 𝑅^3^ = 89%), as well as observed RT (𝑅^3^ = 67%) and RT CV (𝑅^3^ = 68%), all p < 0.0001. Importantly, parameters from each of the three factors contributed uniquely and significantly to these predictions, with standardized beta coefficients ranging from -0.86 to 0.80 (see **SI Table S12**).

These results align with the theoretical constructs of proactive control, reactive control, and attentional modulation, and validate the model’s ability to dissociate different aspects of inhibitory control, thus representing distinct cognitive pathways that they can be dissociated into distinct components characterizing individual differences. They demonstrate that PRAD is a robust cognitive framework for distinguishing among three dissociable pathways of response inhibition beyond reactive control alone (Hu & Li, 2012; Lee & Kang, 2020; Perri, 2020), and contribute towards the development of a richer theoretical framework (Mirabella, 2023).

### Proactive Mechanisms

PRAD revealed the substantial contribution of PDR mechanisms to inhibitory control. Across individuals, proactive cognitive states occurred on an average of 78% of trials (2.5^th^ to 97.5^th^ percentile 3-98%), with PDR accounting for approximately 28% (2.5^th^ to 97.5^th^ percentile 251%) of average reaction times of 515ms (mean PDR = 150ms, 2.5^th^ to 97.5^th^ percentile 8-322ms; **Figure 7A**).

**Figure 7.**
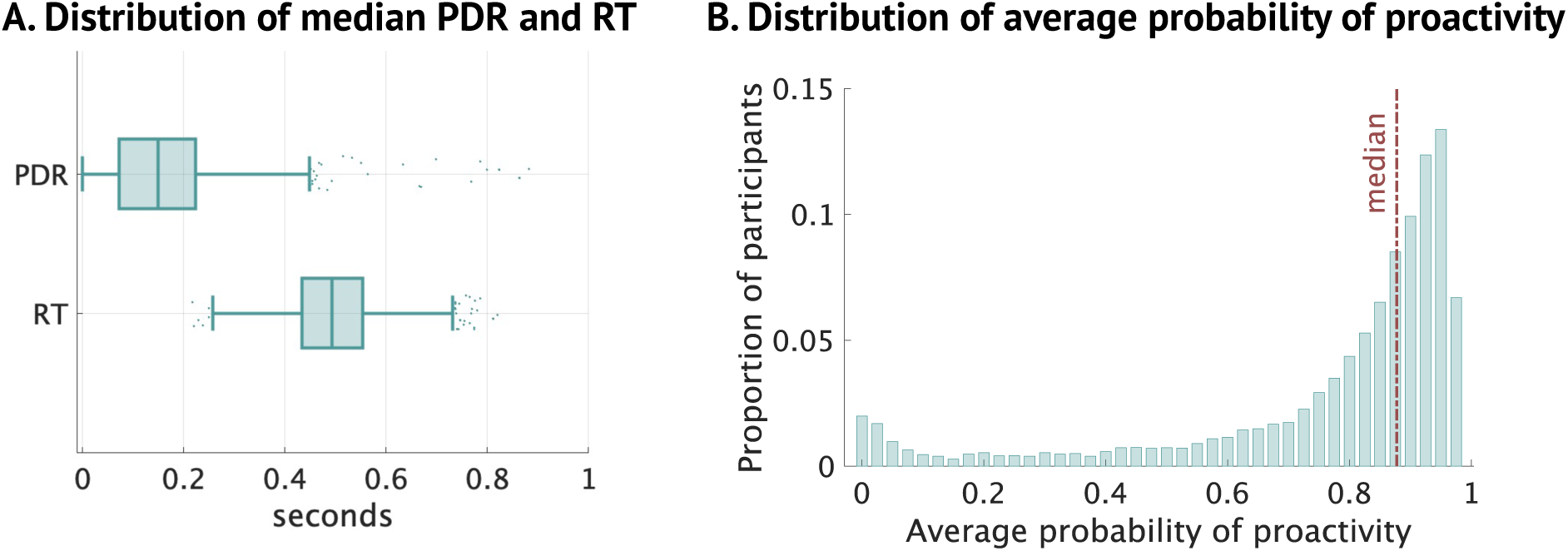
Proactive Delayed Response (PDR). (A) Distribution of median PDR and RT across individuals, showing the range of proactive control strategies. (B) Distribution of average probability of proactive states across participants.

Between subjects, the average PDR is associated with both the mean RT (r = 0.59, p < 0.0001) of responses as well as RTCV across trials (r = -0.51, p <0.0001), and is negatively correlated with SFR (r = -0.59, p < 0.0001), and positively correlated with xSSD (r = 0.61, p <0.0001). The variability in PDR (PDRCV) is correlated with RTCV (r = 0.36, p <0.0001). Linear regressions (**SI Table S13**) show that across individuals, PDR is significantly related to SFR (𝑅^3^ = 0.48, 𝛽 = −0.44, 𝑝 < 0.0001) and xSSD (𝑅^3^ = 0.46, 𝛽 = 0.48, 𝑝 < 0.0001) even after controlling for SSRT. The results demonstrate that individuals with longer delayed responding show more successful inhibitory control (lower SFR, higher xSSD) by appropriate modulation of RTs, even after controlling for the effect of SSRT on successful inhibition. Regression controlling for the influence of average experienced SSD and SSRT shows that the average probability of proactive states is significantly related to SFR (𝑅^3^ = 0.65, 𝛽 = −0.15, 𝑝 < 0.0001). Within-subject regressions also show that after controlling for SSRT, SSD, and variable drift rate across trials, the probability of proactivity, but not the length of proactive delay, is significantly related to stop-failures (**SI Figure S6**).

Higher baseline proactivity (θ_"_) was positively associated with increased PDR (r = 0.44, p < 0.0001) but not so with RT (r = -0.07, p<0.0001), suggesting that PDR mechanisms may be linked to core capabilities, and individuals with faster processing speeds may also have improved top-down PDR regulation. Baseline proclivity for proactive control (𝜃_"_) is also correlated to lower RT coefficient of variability (r = -0.29, p <0.0001).

Crucially, longer proactive delays were associated with more successful inhibition, underscoring the pivotal yet underappreciated role of proactive control in SST performance. This echoes a few previous findings that revealed negative association between proactive control and SSRT and suggested that better proactive control is related to faster stopping speed (Cai, Mizuno, Tomoda, & Menon, 2023; Chikazoe et al., 2009). It is noteworthy that these previous studies relied on additional experimental manipulation to probe proactive control whereas PRAD can identify proactive control components from the standard SST. Such studies with experimentally manipulated versions of the SST task have estimated go response delays in the range of 100-140ms (Castro-Meneses et al., 2015; Chikazoe et al., 2009; Jahfari, Stinear, Claffey, Verbruggen, & Aron, 2010), similar to the PRAD inferred average of 150ms

These findings have important implications. Previous models largely focused on reactive control, which is important for responding to unexpected stimuli in the environment. However, many daily life situations require proactive control to minimize the need for reactive inhibition and reduce impulsivity. As proactive control is generated by an individual’s goals, explicit measurements of it may have greater ecological validity in characterizing everyday response tendencies and psychopathology.

### Error-monitoring and adaptive or maladaptive modulation of proactive states

PRAD reveals selective proactive slowing (delayed responding) of the go process on some trials governed on a baseline propensity for proactivity that is adaptively (or maladaptively) modulated at a trial-level based on error monitoring. Adaptive modulation would involve increasing the probability for proactive slowing after failing to inhibit responses on a stop signal trial, and reducing it after an omission on a trial with no stop signal. Maladaptive modulation would work in the opposite direction. The direction and magnitude of error-monitoring based modulation of this probability based on whether the previous trial was a stop-failure or go-omission is operationalized by the 𝜃_#_ parameter. A higher negative value is optimal, a value close to zero implies no modulation based on error-monitoring, whereas a positive value implies maladaptive modulation. In the latter case, the probability for proactive slowing, would counter-intuitively, be increased after a go-omission and decreased after a stop-failure.

Figure 8A illustrates two actual participants with different values of 𝜃_#_. The first, with a high baseline propensity for proactivity, shows how adaptive modulation (𝜃_#_ = -2.42) helps shift towards non-proactive states after go-omissions. The second, with a low baseline propensity for proactivity, shows how maladaptive modulation (𝜃_#_ = 2.02) helps shift towards proactive states after go-omissions. Figure 8B shows how individual differences in participants’ error-based modulation affect changes in the probability of proactive. For individuals with 𝜃_#_ < 0 (62% of individuals) the average probability of proactive response states post stop-failure increases adaptively to 90% compared to 73% post go-omission (Figure 8C). For individuals with 𝜃_#_ > 0, this maladaptively reduces to 54%, compared to 77% post go-omission. The resulting differences manifest as a difference in reaction times (since proactive slowing increases RT), between post stop-failure and post go-omission trials. This difference depends on whether individuals demonstrate adaptive (𝜃_#_ < 0; mean RT increase of 37ms) or maladaptive (𝜃_#_ > 0; mean RT decrease of 25ms) error-monitoring. Higher values of 𝜃_#_ lead to higher go-omission rates (r = 0.527, p < 0.0001), and lower values of 𝜃_#_lead to higher post-inhibitory error effects, with a negative correlation between 𝜃_#_ and the difference between post-stop error and post-go error RTs (r = -0.38, p < 0.0001).

**Figure 8.**
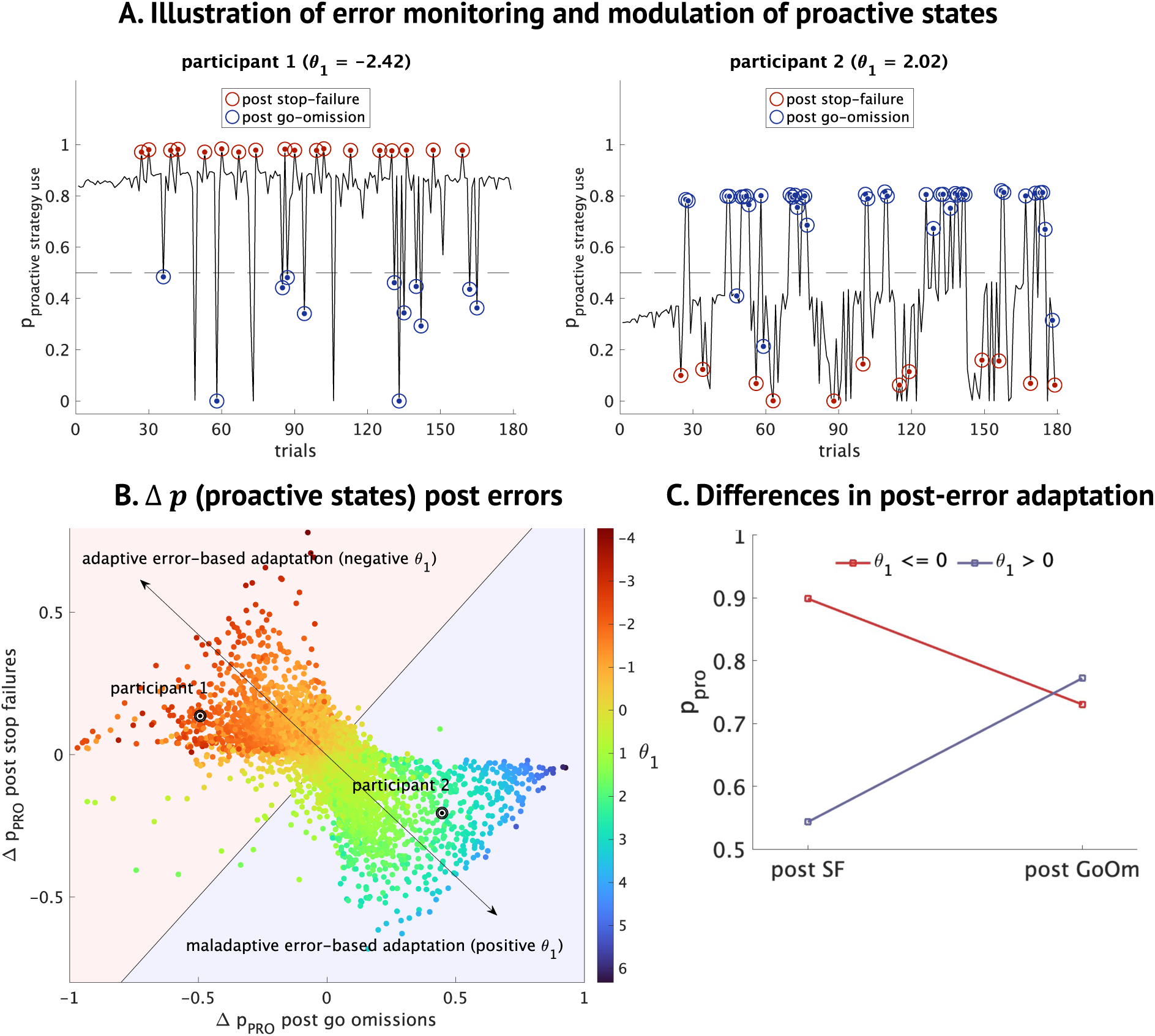
PDR dynamics. (A) Illustration of error monitoring and modulation of proactive states. Trial-level dynamics (probability of proactivity) from two participants with high negative (adaptive) and positive (maladaptive) θ# values. (B) Distribution of error monitoring related sequential effects on probability of proactivity. Distribution of average changes in trial level probability of proactivity after stop failures versus after go omissions. The colors reflect θ#values. Participants with negative values (red) tend to show more adaptive error-based modulation while those with positive values (blue) tend to show more maladaptive error-based modulation of proactivity. (C) Differences in post stop-failure proactivity between θ# subgroups.

This component of PRAD can be construed as engaging error-monitoring and executive function systems, and effecting top-down changes in cognitive states that result in dynamic adjustments in cognitive control. These results also showcase how the model captures post-inhibitory effects (Rieger & Gauggel, 1999) by adaptive or maladaptive modulation of the proactive inhibitory channels.

### Belief Updating processes driving proactive states

The proactive slowing of the go process is based on dynamically updated beliefs about prospective SSDs, which are based on tracking previously experienced SSDs. In PRAD, this belief updating is governed by the 𝜇 (persistence) parameter, where (1 - 𝜇) can be interpreted as the rate of decay. For 𝜇 = 1, the tracked SSD is equal to the average value of all observed SSDs up to that trial. As 𝜇 decreases, a higher recency bias is introduced, with SSD tracking being biased towards more recent values. Thus, while the tracked SSD generally follows the mean experienced values, lower values of 𝜇 track weigh recent observation more heavily. Figure 9A illustrates three actual participants with different values of 𝜇, demonstrating how high decay (low 𝜇 = 0.85) enables tracking of the most recent observed values of SSD while low decay (high 𝜇 = 0.99) corresponds to tracking the mean observed values over all past trials, and intermediate decay (𝜇 = 0.93) values balance between these two extremes. On trials when participants decide to implement a proactive strategy, the go process is delayed by this tracked belief about SSDs. Figure 9B shows how individual differences in participants’ decay rates biases tracked beliefs towards more recent or mean observed SSDs. Persistence in belief updating (μ) affected the absolute error between tracked and current stop-signal delays (r = 0.42, p < 0.0001), with lower μ values associated with higher recency bias, and more accurate dynamic SSD tracking, and hence more well-calibrated PDR (Figure 9C).

**Figure 9.**
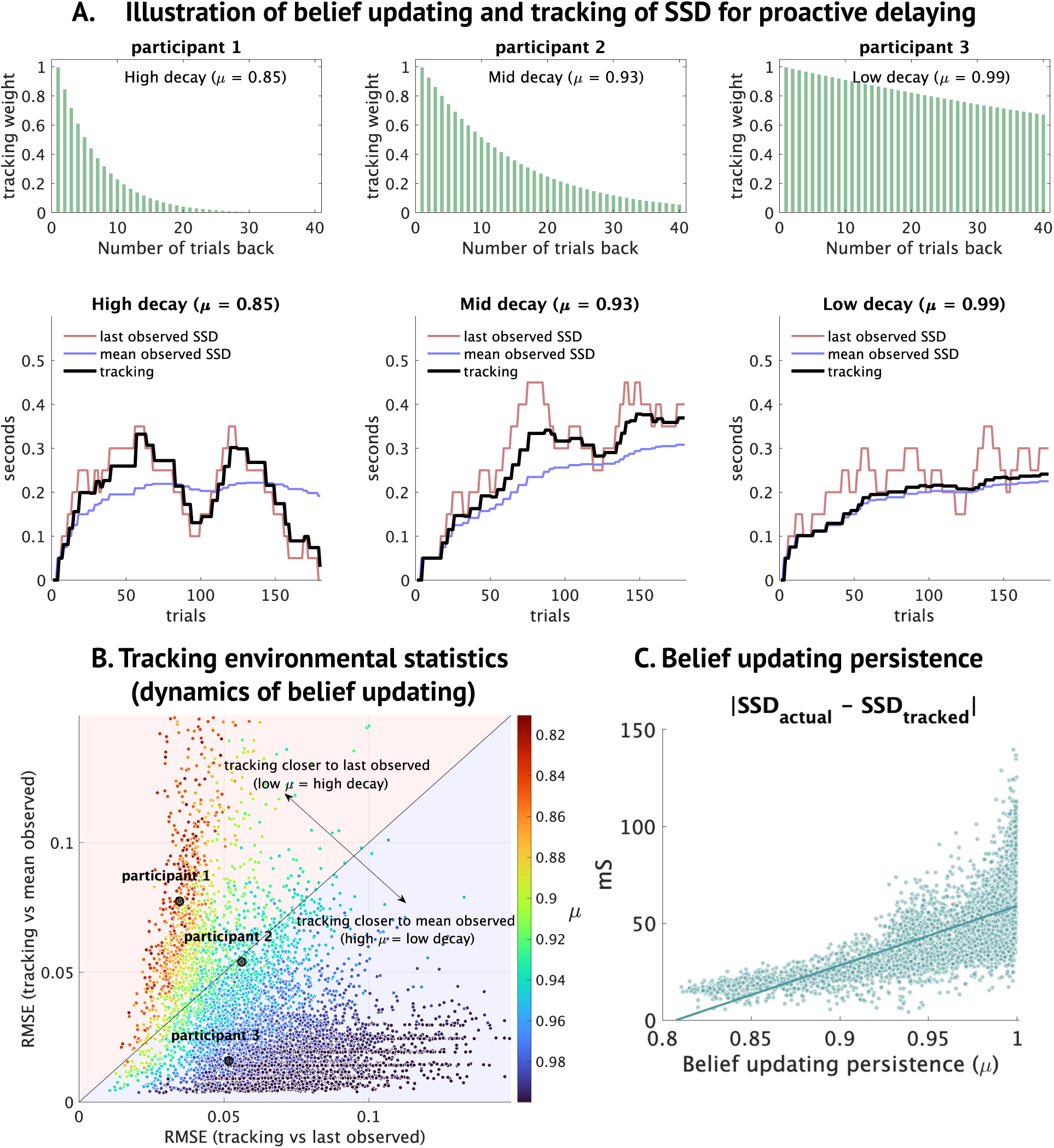
PDR dynamics. (A) Illustration of belief updating and tracking SSD for proactive delaying. Trial-level dynamics including tracking weights (bar chart), and how noisy tracking of SSD (black lines) is related to last observed (red) and mean observed (blue) SSD, depending on high, medium, or low decay rates for belief updating in the tracking process. Participants with higher decay (low persistence 𝜇) are closer to last observed SSD, while those with lower decay (high persistence 𝜇) are closer to mean observed SSD. (B, C) Effect of decay rate / persistence on tracking errors.

This component of PRAD can be construed as engaging both working memory as well as metacognitive awareness about what level of decay might be ideal in this task, and how this ties into dynamic adjustments in cognitive control.

### Attentional Modulation of Stopping

Attentional Modulation of Stopping (AMS) emerged as a crucial component of inhibitory control in the PRAD model. AMS is governed by two key parameters: attention-based adaptivity (γ₁) and attention length (γ0). These parameters modulate the stopping expectancy which in turn affects the variability of SSRT on a trial-by-trial basis.

We observed that within-individual correlation between the number of trials since a stop signal has last been encountered (nSSD) and (a) the observed SFR average at each nSSD (95% CI of correlation coefficient ranging from -0.69 to +0.70), (b) the trial level SSRT (95% CI of correlation coefficient ranging from -0.89 to +0.90) show a wide range of correlations across individuals. Across participants, the average stopping expectancy ranged from 0.43 to 0.66 (2.5^th^ to 97.5^th^ percentile). Across all trials and participants, the stopping expectancy ranged from 0.38 to 0.71 (2.5^th^ to 97.5^th^ percentile). Within individuals, the 95% CI width of stopping expectancy ranges from 0.0 (no modulation) to 0.68, with an average 95% CI of 0.16. This stopping expectancy is the implicit expectation about the probability of encountering a stop signal; it is the initial bias of the stopping drift diffusion process, modulating the baseline reactive stopping process. The within-individual correlation between SFR and trials since stop signal was last encountered is correlated positively with the average stopping expectancy (r = 0.48, p <0.0001) and weakly negatively (r = -0.135, p <0.0001) with the variability in stopping expectancy (measured by coefficient of variability). The absolute value of this within-individual correlation is correlated weakly with average stopping expectancy (r = 0.052, p <0.0001) and positively (r = 0.30, p <0.0001) with the variability in stopping expectancy. These trial-level changes in stopping expectancy form the basis of the AMS process.

The distribution of SSRT and RT (**Figure 10A**) across the sample revealed substantial individual differences, with SSRT showing greater between-subject variability (CV = 0.44) compared to RT (average CV = 0.16). We found that the log ratio of SSRT/RT (**Figure 10B, SI Table S14**) was significantly influenced by both stop process drift rate (β = -0.27, p < 0.0001) and threshold (β = 0.63, p < 0.0001), accounting for 74% of the variance in this measure. Linear regression **(SI Table S14**) reveals that after controlling for stop process drift, response thresholds, and nondecision time (i.e., the remaining parameters affecting SSRT), the AMS driven stopping expectancy still has an influence on SSRT (𝑅^3^ = 0.84, 𝛽 = −0.21, 𝑝 < 0.0001).

**Figure 10.**
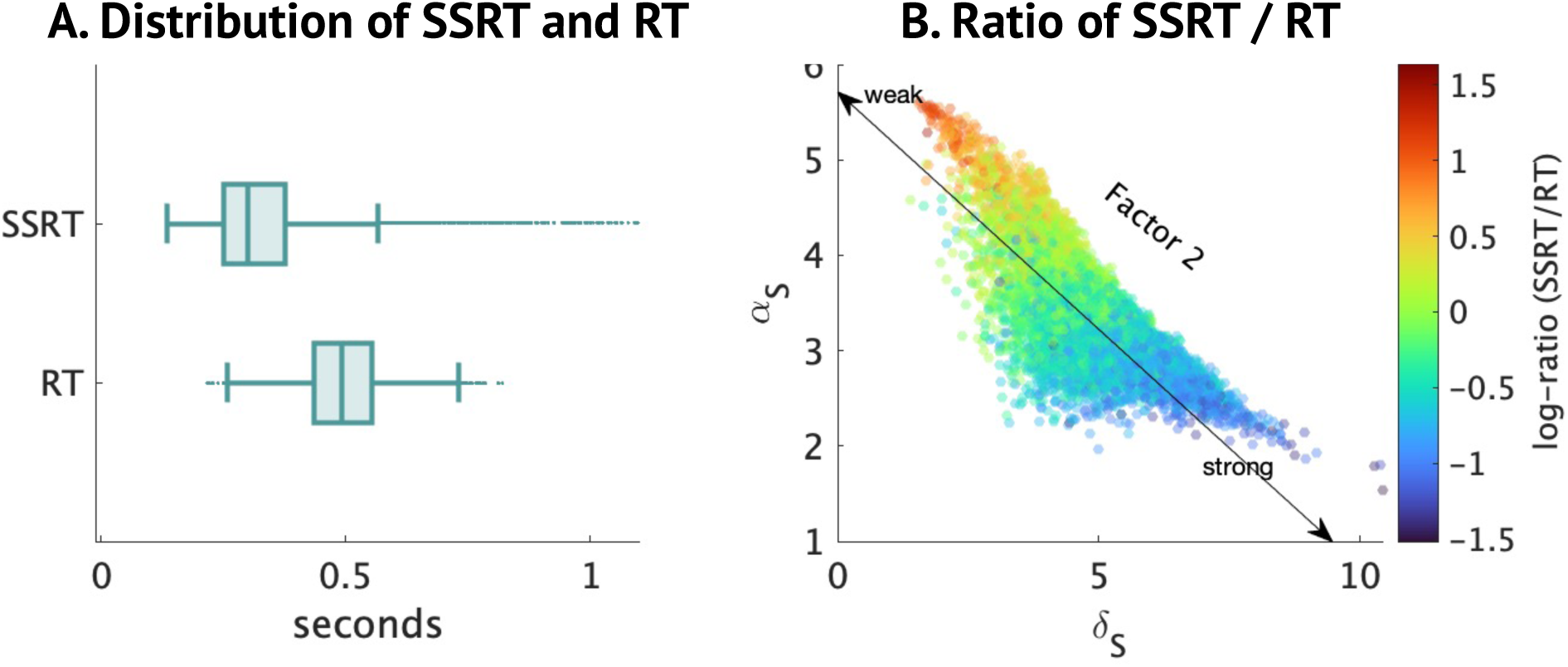
Attentional Modulation of Stopping (AMS). (A) Distribution of median SSRT and RT across individuals. (B) Distribution of the log ratio of SSRT to RT across participants, and link to stopping process parameters.

The variability in stopping expectancy represents the strength of the AMS effect, and the coefficient of variation of stopping expectancy ranged from 0.005 (2.5^th^ percentile; weak AMS effect) to 0.229 (97.5^th^ percentile; moderate AMS effect). The SSRTCV is positively correlated with variability in stopping expectancy (r = 0.55, p <0.0001). Linear regression (**SI Table S14,** 𝑅^3^ = 0.83) 𝑠hows that average stopping expectancy (𝛽 = 0.36, 𝑝 < 0.0001) and coefficient of variation of stopping expectancy (𝛽 = 0.55, 𝑝 < 0.0001) have a significant influence on SSRTCV even after controlling for the effects of stop process drift rate and stop process decision threshold.

These findings demonstrate the pervasive influence of attentional modulation on stop process dynamics across our sample, highlighting its role in explaining individual differences in inhibitory control performance.

### Differential patterns of attentional modulation of stopping

To further quantify the impact of attentional modulation on stop process dynamics across our sample, we analyzed subgroups based on their attentional modulation parameter (γ₁). Analysis revealed that 58% of participants showed decreasing attentional control (γ₁ < 0) as the number of trials since the last stop signal increased, while 42% showed increasing control (γ₁ > 0). Here increasing attentional control refers to increasing bias of the stopping process, which on the presentation of a stop signal stimulus would result in faster SSRTs.

We found significant differences between these subgroups in several key measures as a function of the number of trials since the last stop signal (nSSD). Changes in stopping expectancy with nSSD (**Figure 11A**) showed a divergent trend between subgroups (t(7785) = 839, p < 0.0001), with the γ₁ > 0 group maintaining higher expectancy as nSSD increased (average r = 0.96), while the γ₁ < 0 group showed decreasing expectancy (average r = -0.97). This translated to significant differences in the correlation of SSRT **(Figure 11B)** and nSSD between subgroups (t(7785) = 157, p < 0.0001), and correlation of observed SFR **(Figure 11C)** and nSSD between subgroups (t(7785) = -52, p < 0.0001), with the γ₁ > 0 group showing more stable performance, in terms of reducing SFR with nSSD (average correlation -0.21 vs 0.16). Thus, AMS parameters significantly influenced patterns of stop-failure rates.

**Figure 11.**
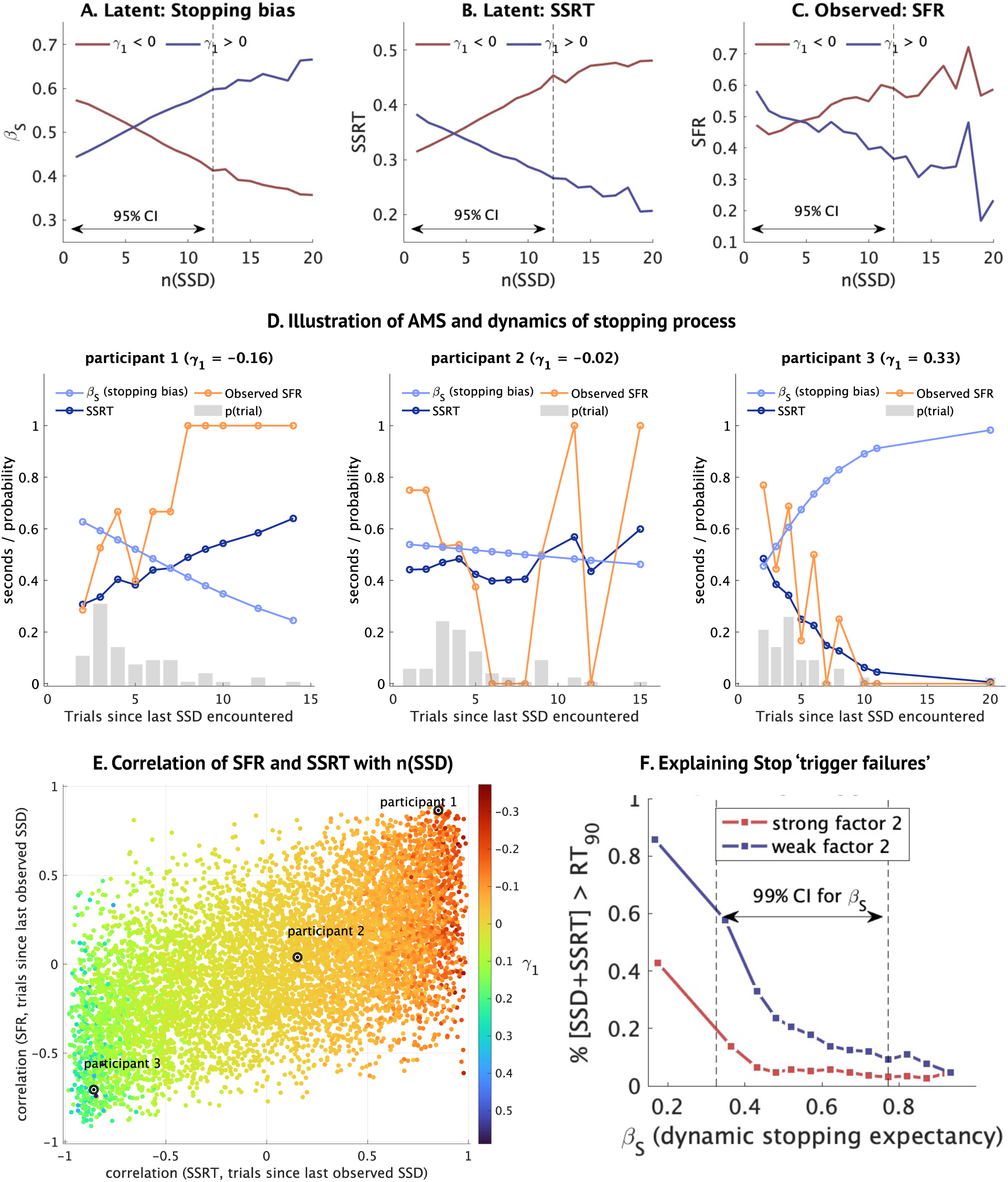
Attentional Modulation of Stopping (AMS) dynamics. (A-C) Evolution of stopping dynamics (expectancy, SSRT, and SFR) with increasing trials since last stop signal (nSSD) for different attentional modulation subgroups (γ#> 0 vs γ#< 0), showing adaptive and maladaptive patterns. (D) Illustration of AMS and dynamics of stopping process. Trial-level dynamics including stopping expectancy or bias (light blue), the resulting SSRT (dark blue), and the subsequently resulting SFR (orange), as a function of n(SSD), the number of trials since a stop signal was last encountered. The participants with different positive and negative values of γ# show different patterns of positive and negative change. (E) Distribution of how SSRT and SFR are related to n(SSD) across participants, based on the underlying γ# values. (F) Explanatory mechanisms for stop ‘trigger failures’. Proportion of stop trials where the SSRT+SSD was higher than the 90th percentile RT (at an individual level), as a function of trial level dynamic stopping expectancy.

Figure 11D illustrates three actual participants with different values of γ_#_. The first, with a negative value, shows how stopping bias (implicit expectation of a stop signal) reduces with increasing trials since a stop signal was last encountered (nSSD), leading to a corresponding increase in SSRT as well as SFR at high nSSD. The second, with γ_#_ close to zero, show no specific trend with nSSD, while the third, with positive γ_#_ shows increasing stopping bias, along with reducing SSRT and SFR with higher nSSD. This behavior would be compatible with an explanation of stronger sustained attention. **Figure 11E** shows how this generalizes across participants as the value of γ_#_varies.

These results show that the AMS process has a significant effect on SSRT and variability of SSRT within individuals, as well as on the divergence of observed patterns of reducing or increasing stopping capacity as the time since the last stop signal was encountered increases. Thus, the AMS effect seems to be measuring the capacity for sustained attention and its impact on inhibitory capabilities.

### Mechanism for trigger failures

AMS processes also provided a continuous process explanation for what have previously been termed trigger failures (R. Doekemeijer et al., 2021; Matzke et al., 2017; Skippen et al., 2019), or assumed failures to initiate the stopping process **(Figure 11F)**.

The total SSD + SSRT is higher than the 90^th^ percentile individual RT for 14% of stop trials across individuals (95% CI from 0% to 82%; average 5% for stronger factor 2 and 22% for weaker factor 2). The AMS mechanism in the PRAD model allowed the distribution of SSRTs to include significantly higher valued SSRTs on some trials, including SSRTs that were higher than the typical RTs, especially for participants low on factor 2 (weak reactive inhibition), as seen in **Figure 10B**. Such trials, in the absence of the PRAD model, which allows for higher variability in SSRT, would be difficult to explain and be classified as trigger failures within the traditional account. Compared to the PRAD model, the control RVM model had significantly lower within-individual SSRT variability (SD) (*t*(7784) = -58, p<0.0001; 89% of the individuals had higher inferred within-individual SD of SSRT across trials in the PRAD compared to RVM). The mean SSRT SD/CV was 0.04/0.119 in the control model compared to 0.060/0.173 in PRAD.

The model identifies not only participants who are more likely to be identified as having a higher proportion of trigger failures, but also what type of trials these are more likely to be encountered on – demonstrating that the inferred trigger failure mechanisms may be exceedingly long SSRTs that are predicated on low stopping expectancy, which in turn is predicated mostly on decreasing expectancy with increasing trials since a stop signal was last encountered. This account is compatible with attentional accounts of trigger failures but provides a strong basis for these attentional lapses based on continuous variability in the stopping process modulated by AMS.

### PDR and AMS uncover district sources of intra-individual variability

PRAD uncovered complex interaction dynamics between proactive and reactive control mechanisms. Figure 12 demonstrates the variability in PDR and AMS processes across 4 individuals, illustrating how different combinations of these processes can lead to similar overall performance but through distinct cognitive mechanisms. Figure 12 compares inferences from four actual participants: The first two participants have a high variability in PDRs (row 1), whereas the latter two have a low variability in PDR states, with participant three having almost no PDRs, and participant four having a consistently high level of PDRs. This manifests as high variability in RT in the first two, but low variability of RT in the latter two participants (row 2). Participants one and four have a high variability in of stopping expectancy based on the AMS mechanism, whereas participants two and three have a stable stopping expectancy across trials (row 3), representing minimal attentional modulation. This manifests as highly variable SSRT for the former two, and low variability in SSRT for the latter (row 4). This illustrates how for instance, very different underlying behavioral patterns between participants 3 and 4 may result in similar iSSRT estimates based on traditional parametric methods. It also illustrates how the presence or absence of a strong proactive components (participant 2 vs 3) can reverse inferences about SSRT between the PRAD model and traditional parametric estimates. Participants two and four have very similar SFR and similar model inferred median SSRT, although the source of trial level variability is very different between these participants.

**Figure 12.**
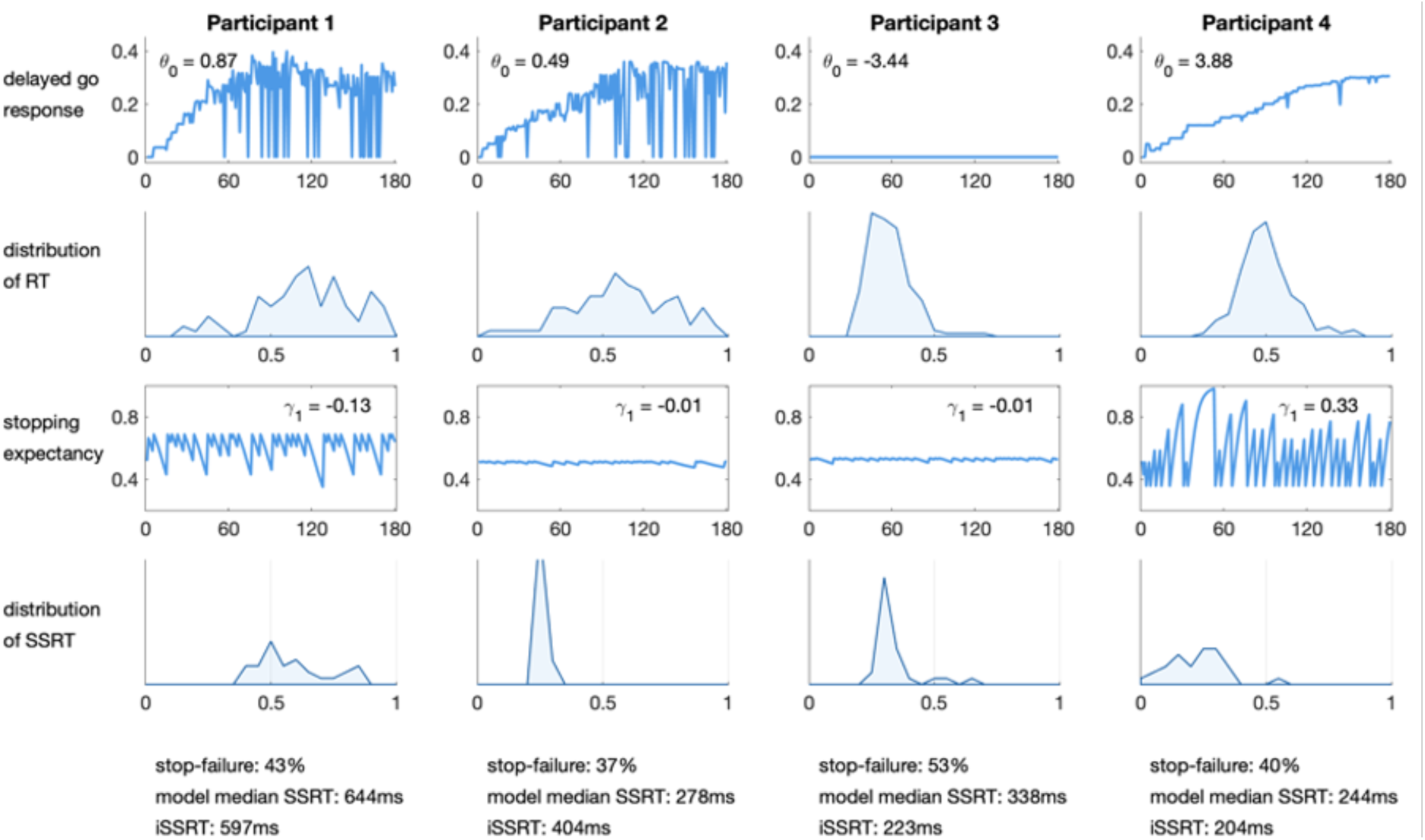
Illustration of variability in PDR and AMS processes across four representative participants. Comparison of trial-by-trial fluctuations in PDR, RT, stopping expectancy, and SSRT, demonstrating how similar overall performance can arise from different underlying cognitive dynamics. This figure highlights how the PRAD model can dissociate different sources of intra-individual variability that may be indistinguishable using traditional measures.

These illustrative results highlight how the model can dissociate different sources of IIV while observed summary level behavior may be similar. These results also highlight the importance of considering the dynamic interplay between different control mechanisms when characterizing individual differences in inhibitory control.

### PDR and AMS as compensatory mechanisms for baseline reactive processes

We ran regression models evaluating the effect of factor 2 scores (representing baseline reactive inhibitory process), average proactive delayed response times, and the interaction between these measures, on stop failure rate (SFR). The interaction model is significant (𝑅^3^ = 0.62, p < 0.0001), with significant influence of factor 2 scores (standardized 𝛽 = 0.2, p < 0.0001), proactive delays (standardized 𝛽 = -0.51, p < 0.0001), and interaction (standardized 𝛽 = -0.37, p < 0.0001). This reveals that longer average proactive delays reduce stop failure rates as expected, but more importantly, the interaction shows that baseline reactive processes show sharp differences in SFR between low and high basic reactive processes when proactive delayed responses were low, but these differences were minimal when proactive delayed responses were high. Thus, proactive delayed responses could compensate for weaker reactive baseline processes.

Next, we ran a regression model evaluating the effect of factor 2 scores (representing baseline reactive inhibitory process), average stopping expectancies, and the interaction between these measures, on stop failure rate. The interaction model is significant (𝑅^3^ = 0.35, p < 0.0001), with significant influence of factor 2 scores (standardized 𝛽 = 0.58, p < 0.0001), stopping expectancies (standardized 𝛽 = -0.15, p < 0.0001), and interaction (standardized 𝛽 = -0.10, p < 0.0001). This reveals that higher stopping expectancies could compensate for weaker reactive baseline processes.

### Dynamic interactions between PDR and AMS

PRAD uncovered complex interaction dynamics between proactive and reactive control mechanisms. Figure 13 illustrates how individuals can be grouped based on their PDR and AMS parameters, revealing four distinct adaptivity profiles. These profiles demonstrate various compensatory relationships between proactive and reactive control.

**Figure 13.**
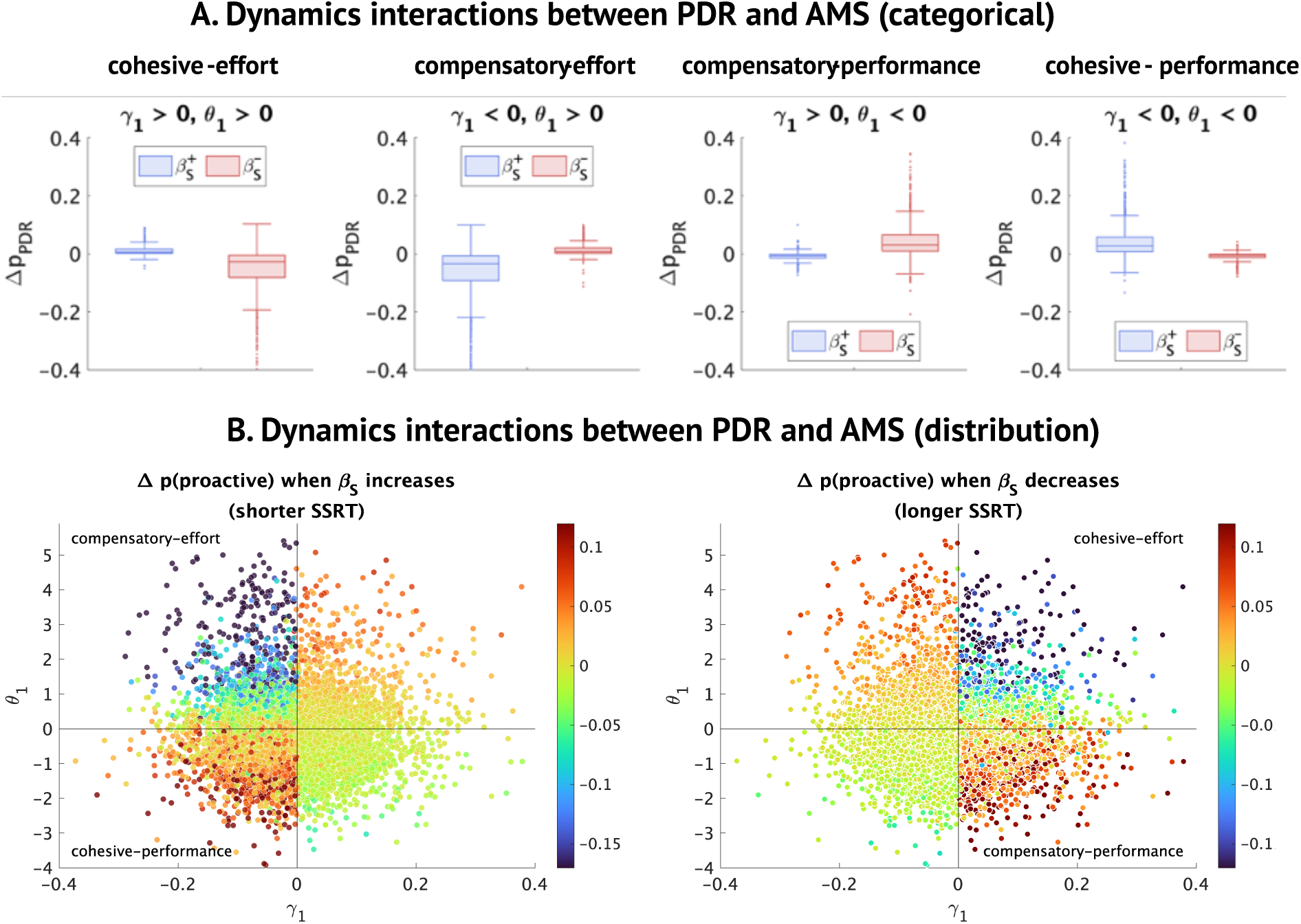
Dynamic interactions between PDR and AMS processes. (A) Four categories of interactions based on values of γ#and θ#, and the resulting changes in trial level probability of proactivity when stopping expectancy (bias) increases or decreases. (B) Distribution of changes in trial level probability of proactivity (color coded) when stopping expectancy (bias) increases (left) or decreases (right), as a function of γ#and θ#.

Specifically, individuals with θ_1_ < 0 and γ_1_ > 0 (25.7%) showed the most adaptive dynamic regulation of processes affecting inhibitory control. Here, when stopping expectancy decreased (worsening SSRT), the probability of proactivity increased, showing compensatory proactive mechanisms that worked towards maintaining performance levels (*compensatory – performance* in **Figures 13A,B**).

Conversely, individuals with θ_1_ > 0 and γ_1_ > 0 (16.8%) showed reduction of proactive probability when SSRT worsened, indicating maladaptive proactive regulation. Rather than compensating, both reactive and proactive modulation moved in a cohesive manner towards reducing effort (*cohesive - effort* in **Figures 13A,B**).

Individuals with 𝜃_#_ > 0 and 𝛾_#_ < 0 (21.6%) show reduction of proactive probability when SSRT improves, compatible with a compensatory mechanism that works towards effort-reduction by reducing proactivity when reactive control is stronger (*compensatory – effort* in **Figures 13A,B**).

Finally, individuals with 𝜃_#_ < 0 and 𝛾_#_ < 0 (36%) show an increase in proactive probability when SSRT improves, compatible with a cohesive internal system that improves both reactive and proactive processes dynamically, possibly in stronger anticipation of a stop trial, but may not result in strong compensatory action between proactive and reactive mechanisms (*cohesive – performance* in **Figures 13A,B**).

Thus, the PRAD model allows inferences about multiple trait-like parameters governing individual differences in dynamic regulation, which provide a multi-faceted view of adaptivity. The dissociated PDR and AMS processes identified under PRAD reveal distinct and complementary sources of within-individual adaptivity.

### Nonergodicity in cognitive control processes

Nonergodicity in a behavioral context occurs when statistics of a behavior over time (within-individual dynamics) do not converge to statistics of the behavior over individuals (between-individual dynamics) (Fisher, Medaglia, & Jeronimus, 2018). In other words, nonergodic processes exhibit different inferences when behavioral dynamics are analyzed at the within-individual level compared to the between-individual level. This distinction is crucial because within-subjects conclusions are often drawn from between-subjects inferences (Curran & Bauer, 2011). Yet, such generalizations are only valid for ergodic processes (Molenaar, 2004).

PRAD revealed substantial evidence for nonergodic processes in inhibitory control. Specifically, the model uncovered differences between within-subject and between-subject correlations for both latent and observed measures (Figures 14A-D), indicating nonergodicity.

**Figure 14.**
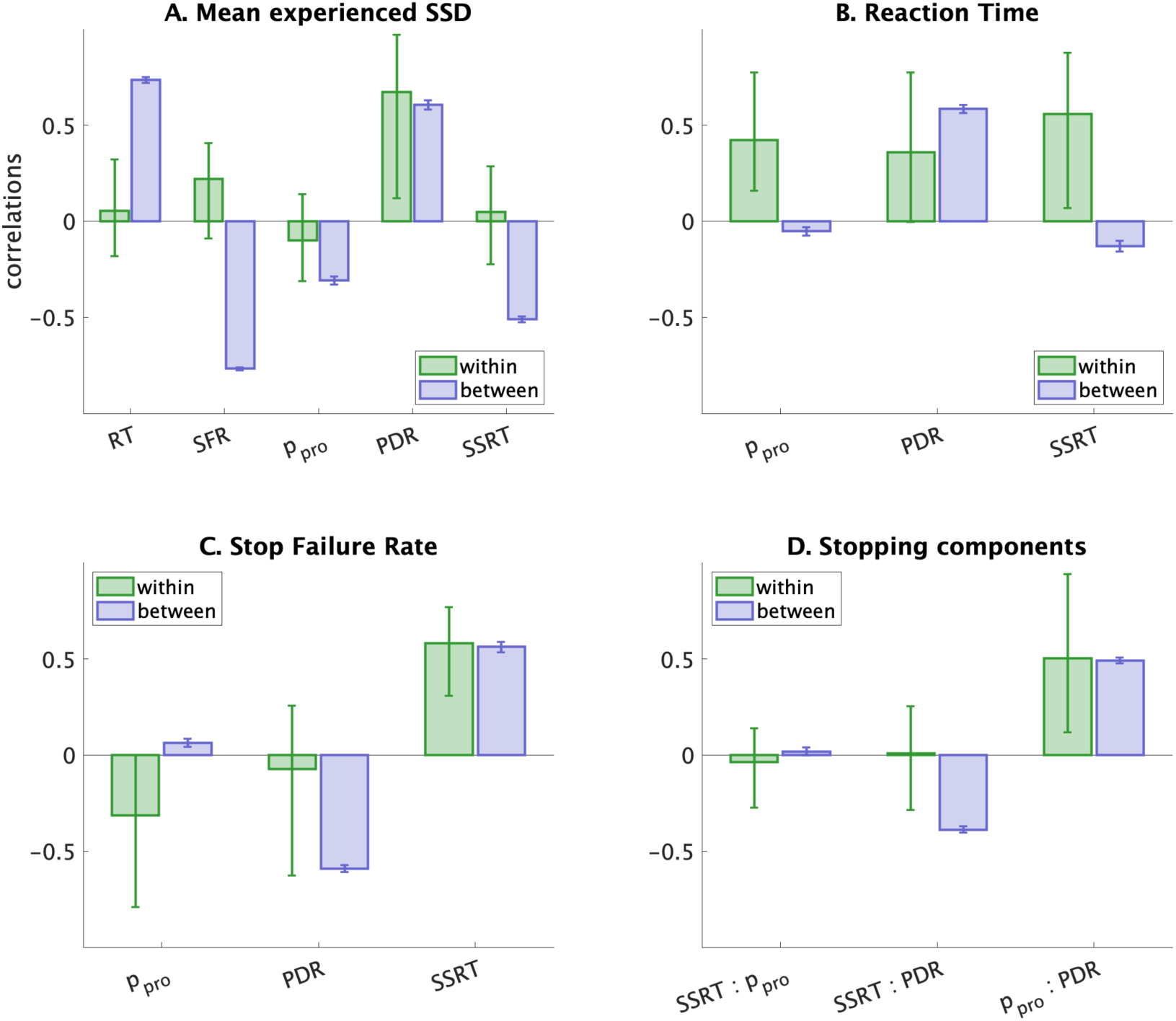
Evidence for nonergodicity in cognitive dynamics. Comparison of within-subject (green) and between-subject (blue) correlations for various pairs of observed and latent variables. Differences between these correlations, especially sign changes, indicate nonergodic processes. Notable nonergodic effects are observed in some of the relationships between mean experienced SSD, SFR, SSRT, RT, PDR, and probability of proactive state. The between-subjects values plotted are based on average values for individuals and bootstrapping the sample. SFR: Stop failure rate, SSRT: Stop-signal reaction time, RT: Reaction time, PDR: Proactive delayed responding, ppro: probability of proactive cognitive state. Error bars reflect 2.5th to 97.5th percentiles for within subject correlations and 2.5th to 97.5th percentiles based on bootstrapped samples for between subject correltions.

For example **(SI Table S15**), we observed opposite patterns of within-subject and between subjects correlations, in the associations between mean experienced stop-signal delay and stop-failure rate (average within r = 0.22, between = -0.77), RT and probability of proactive state (average within r = 0.42, between = -0.05), RT and SSRT (average within r = 0.56, between = - 0.13), stop-failure rate and probability of proactive state (average within r = -0.31, between = 0.06), SSRT and average proactive delay (average within r = 0.01, between = -0.39). These findings suggest that the interactions among cognitive processes underlying response inhibition are nonergodic, and that within-individual dynamics cannot be fully captured by between-individual analyses.

The acknowledgment of nonergodicity calls for dynamic, individualized approaches to analyzing cognitive data, challenging the conventional reliance on aggregated metrics and static models (Aristodemou, Rommelse, & Kievit, 2022; Cañigueral et al., 2022; Epstein et al., 2022; Karalunas, Geurts, Konrad, Bender, & Nigg, 2014; Kofler et al., 2013; Kuntsi, 2014; Ode, Robinson, & Hanson, 2011; Paraskevopoulou, Coon, Brunner, Miller, & Schalk, 2021; Tamm, Epstein, & Becker, 2019; Wiecki, Poland, & Frank, 2015; Yamashita et al., 2021). The implications of these findings are twofold. First, they highlight the importance of considering individual differences in cognitive dynamics when studying response inhibition and other cognitive processes. Group-level analyses may not adequately capture the complex, time-dependent relationships between cognitive processes within individuals. Second, the presence of nonergodicity along with individual differences in within-subject dynamics suggest that personalized approaches to understanding and modifying cognitive control deficits may be necessary. Interventions targeting specific cognitive processes, such as proactive control or attentional modulation, may have different effects depending on an individual’s unique cognitive dynamics.

### PRAD reduces model and measurement biases

#### Systematic compensation for known biases in traditional SSRT

PRAD-inferred SSRT demonstrated systematic compensation for known biases in traditional SSRT estimates (Bissett, Hagen, et al., 2021; Frederick Verbruggen et al., 2019; Verbruggen et al., 2013). Comparing model-based median SSRT (mean 340ms, SD 155ms) to non-parametric integration method SSRT (iSSRT, mean 302ms, SD 135ms) revealed that traditional iSSRT estimates were lower by 38ms on average (t(7786) = -32.3, p < 0.0001), and showed a smaller degree of individual differences (SD lower by 20ms) between individuals (F(7786,7786) = 0.76, p < 0.0001).

Crucially, PRAD SSRT estimates were significantly higher for conditions known to lead to underestimation of traditional non-parametric iSSRT (Figure 15A-E**; SI Table S16**). These included participants with higher right skew of RT (F(3,7783) = 21.9, p < 0.0001), larger RT slowdown (F(3, 7783) = 9.3, p < 0.0001), high stop success rates (F(2,7784) = 76.8, p < 0.0001), higher go-omission rates (t(7785) = 3.4, p < 0.0001) and those classified as race-model violators (t(7785) = 28.9, p < 0.0001).

**Figure 15.**
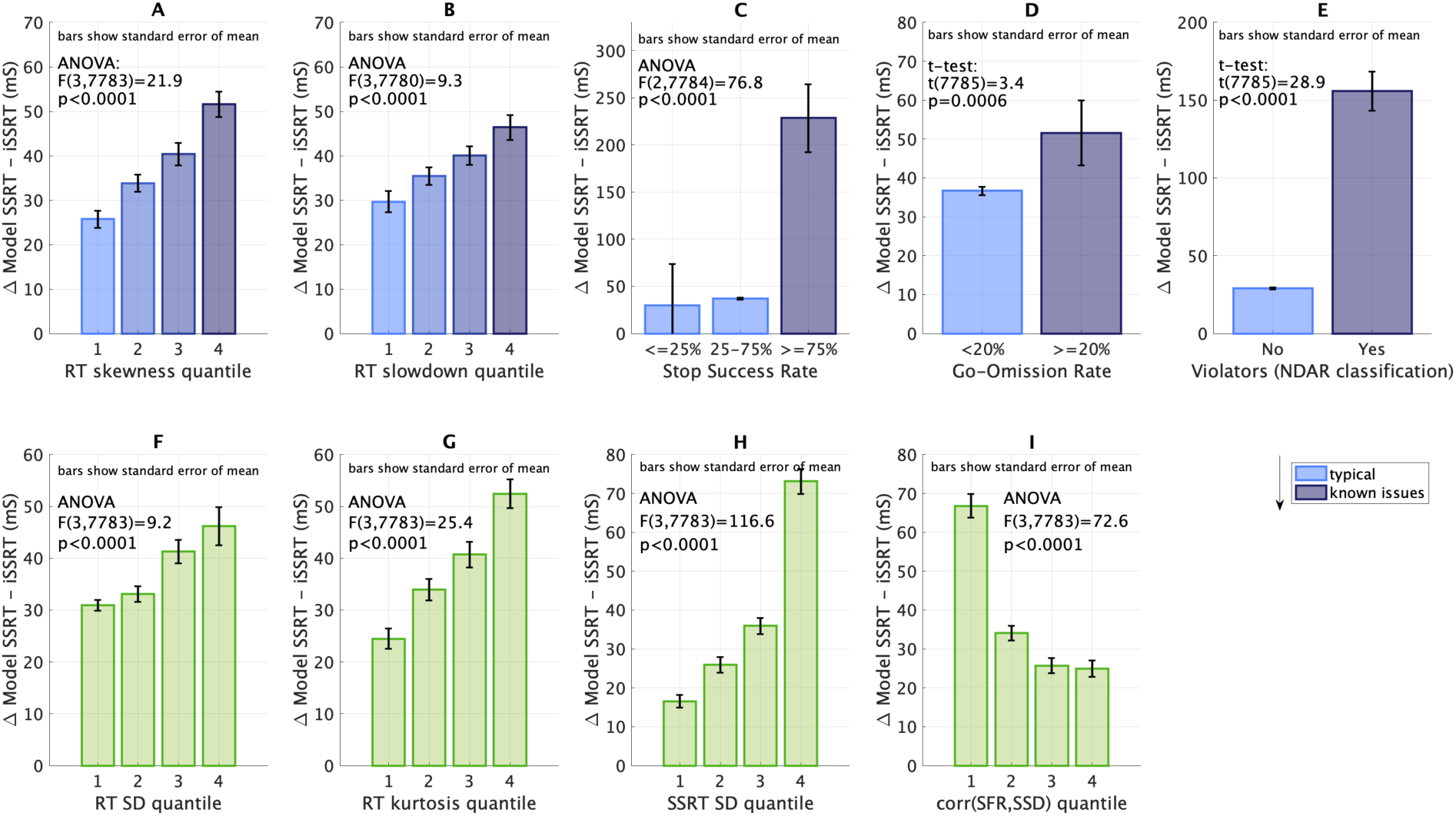
Debiasing of stop-signal reaction time (SSRT) estimates. Comparison of PRAD-inferred SSRT with traditional non-parametric integrated SSRT (iSSRT) across various conditions. (A-E) Conditions known to bias SSRT estimates, including high RT skewness, RT slowdown, stop success rates, go omission rates, and context independence violations. (F-I) Additional conditions revealed by PRAD, such as high RT variability and kurtosis, high SSRT variability, and low SSD-SFR correlations. Higher PRAD SSRT estimates in these conditions demonstrate the model’s ability to compensate for known biases and reveal new insights into SSRT estimation.

For each of these, participants were split into groups based on the relevant measure, and the difference between the PRAD model inferred median SSRT and non-parametric integrated SSRT (iSSRT) was assessed for group differences using ANOVA or t-tests as appropriate. For RT skewness and RT slowdown, participants were split into four groups (quantiles) based on these measures. For stop success rates, participants were split into three groups based on stop success being < =25%, 25%-75% and >=75%. Previous literature has recommended assessing nonparametric SSRT only when stopping success ranges in the 25%-75% range, with underestimation reported for high success rates (Frederick Verbruggen et al., 2019; Verbruggen et al., 2013). For go omission rates, participants were split into two groups based on go-omission rates being < or > 20%. Previous literature has recommended avoiding non-parametric SSRT when go-omission rates are high (Frederick Verbruggen et al., 2019; Verbruggen et al., 2013). For Violators (NDAR classification), participants were split into two groups based on whether they were classified as violators as per the ABCD NDAR classification, which infers violators as children who seem to violate traditional assumptions of context independence. Significant group differences *and* increasing SSRT difference as skewness / slowdown increased (Frederick Verbruggen et al., 2019; Verbruggen et al., 2013), stop success rates / go-omission rates were higher, or for violators, were indicative of debiased PRAD SSRT.

We confirmed that the relative quality of PRAD model fit under these conditions remained stronger compared to control models, thus validating these inferences (**SI Table S16**). PRAD robustly identifies and characterizes response inhibition even when assumptions of the race model and context independence are violated, effectively compensating for known, systematic biases in conventional measures. Thus, PRAD effectively characterizes and rectifies biases inherent in traditional cognitive control models. This is especially important when investigating neurodevelopmental disorders and older adults who exhibit diminished performance and greater variability.

Additionally, PRAD revealed systematic differences in previously unexamined conditions, such as high RT variability (F(3,7783) = 9.2, p<0.0001), high RT kurtosis (F(3,7783) = 25.4, p<0.0001), high SSRT variability (F(3,7783) = 116, p<0.0001), and low correlations between SFR and SSD (F(3,7783) = 72.6, p<0.0001) (Figures 15F-I). This detection of other systematic patterns of differences from non-parametric methods, while exploratory, is indicative that there may be other previously unexamined specific patterns of behavior that lead to biases in traditional non-parametric iSSRT estimates, that can be detected by the PRAD models.

These results demonstrate PRAD’s ability to provide more accurate and unbiased SSRT estimates across a wide range of performance patterns, particularly for individuals who deviate from typical performance profiles, when context independence is violated, or for behavioral patterns that are not ideal for non-parametric estimation.

#### Multiple PRAD mechanisms underlie bias reduction and differentiate behavioral subgroups

Biases in traditionally measured SSRT or the ability of models to explain behavior is linked to behavioral patterns such as violations of context or stochastic independence, high skew, variability, or slowdown in RT, extreme patterns of SFR or go-omissions, and sequential dependencies. We analyzed which PRAD model parameters differentiated behavioral subgroups related to these behavioral patterns (Figure 16), and find that parameters related to proactive mechanisms (factor 1), basic reactive mechanisms (factor 2), and the go-process show strong effects sizes in terms of differences between behaviorally defined sub-groups.

**Figure 16.**
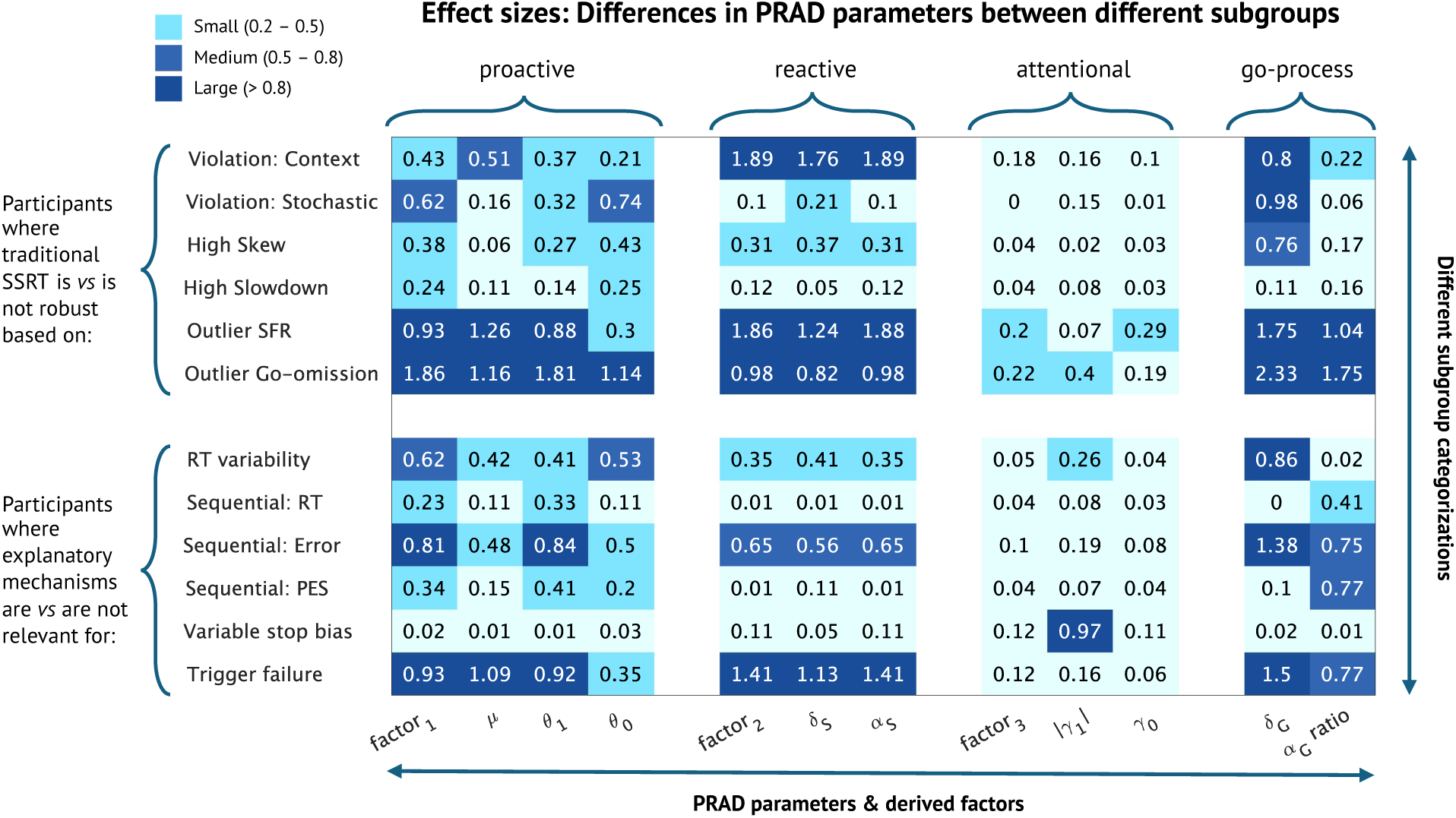
Effect sizes for differences in PRAD parameters between different subgroups. Each row reports the effect size for difference in the PRAD parameters and derived factors between two subgroups. For instance, the first row depicts the effects sizes for differences in parameters between participants who were identified as demonstrating violations of context independence, and those who were not. These subgroups reflect the groups depicted in Figure 2.

### Reducing typicality biases and improving explanations of individual differences for atypical profiles

We examined the model fits and residuals for PRAD and the control RVM model on key behavioral measures. PRAD provided improved fits (lower absolute residuals) to data compared to RVM for RT (Cohen’s d 0.24), RT SD (Cohen’s d 1.04), SFR (Cohen’s d 0.85), and correlation between SFR and SSD (Cohen’s d 0.27).

For each measure, we evaluated the mean absolute residuals by population subgroups based on: (i) family income (based on whether income was lower than or greater than $50k annually), (ii) age (median split), (iii) whether children demonstrated externalizing behavior (based on CBCL externalizing t-scores > 64), (iv) cognitive ability (median split using NIH cognitive toolbox scores), (v) brief problem monitoring assessment (BPM scale > 0), (vi) behavioral inhibition score (median split using the BIS subscale from BIS/BAS), and (vii) adverse life events (median split using ALE scores). We then compared these mean absolute residuals between the nontypical and typical subgroups (lower vs higher cognitive ability, younger vs older children, lower vs higher family income, higher vs lower externalizing, ALE, BIS, and BPM scores) to compute a *typicality bias*.

We observed that while for most measures, both PRAD and RVM demonstrated some typicality bias (relatively better fits for typical vs nontypical subgroups), this bias was lower in the PRAD model vs RVM across all measures and for all of the seven typicality subgroups tested above (**SI Table S17**). Figure 17 and **SI Table S17** show the percentage reductions in these measures of typicality bias by using PRAD compared to the RVM model. Across these four measures and seven subgroup comparisons, the percentage reduction ranged from 2%-113% with a mean value of 47%. Importantly, the PRAD model showed lower mean absolute value of residuals for both typical and nontypical subgroups compared to RVM across all comparisons (i.e., the PRAD reduction in typicality bias was because of improvements in fits for atypical groups, not because of poorer fits for typical groups). This reduction in typicality bias compared to the RVM (which is essentially a Bayesian implementation of a standard horse-race model with competing go and stop processes, and random variability in SSRT across trials) demonstrates that the PDR and AMS additions to the PRAD model allow us to more effectively capture the full spectrum of heterogeneity present in both typical and non-typical or under-represented developing populations.

**Figure 17.**
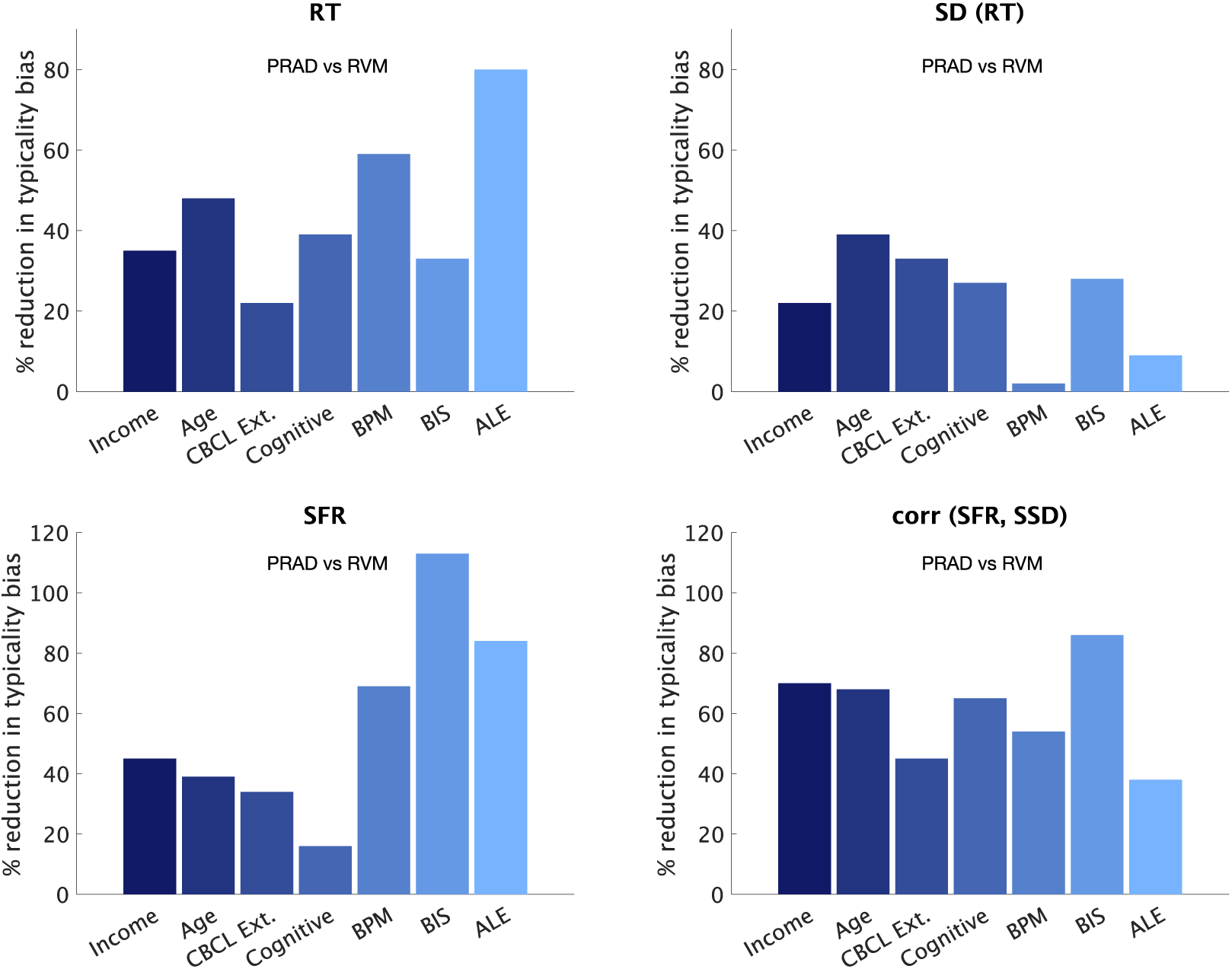
Percentage reduction in typicality bias based on PRAD vs RVM control models. The plots show the percentage reduction in typicality bias, measured as the reduction (from RVM to PRAD) in residual errors in posterior model values of RT, SD(RT), SFR, and correlation between SFR and SSD difference between atypical and typical populations. While PRAD and RVM both demonstrate higher residual errors for atypical populations, PRAD significantly reduces the gap in residual errors between atypical and typical populations, thus debiasing inferences for atypical populations. Atypical populations tested here are based on income (children from poorer families), age (younger children), CBCL externalizing (children with CBCL externalizing t-scores > 64), cognitive abilities (children with lower cognitive scores on the NIH cognitive toolbox), Brief Problem Monitoring survey (children with non-zero scores), BISBAS scale (children with higher inhibition subscale scores), and ALE (children with high bad event scores for adverse life events).

### Relevance to cognitive performance, clinical subgroups, neural mechanisms, and exposomic factors

#### Improved Prediction of Cognitive Performance

PRAD parameters demonstrated superior predictive power for performance on NIH Toolbox cognitive tasks (Akshoomoff et al., 2013; Slotkin et al., 2012) compared to traditional SSRT measures. The NIH-Toolbox (NIH-TB) Cognition Battery subtests were used to measure a diverse set of cognitive domains, including processing speed, executive function, working memory, and language. Uncorrected scores for the Fluid Cognition Composite Score, Crystallized Cognition Composite Score, and Total Cognition Composite Score (Akshoomoff et al., 2013; Slotkin et al., 2012), as well as individual subscores on the following NIH-TB Cognition Battery subtests were analyzed: Dimensional Change Card Sort Test (Executive Function/ Set-Shifting), Picture Vocabulary Test (Language), List Sorting Working Memory Test (Working Memory), Flanker Inhibitory Control and Attention Test (Executive/Attention), Pattern Comparison Processing Speed Test (Processing Speed), Oral Reading Recognition Test (Language)

Using support vector machines (SVM regression model with a Gaussian kernel, to assess the fit as well as 10-fold cross-validated predictions) to fit the overall NIH cognitive toolbox scores, we found that adjusted R² improved from 6.3% to 27.7% when using PRAD parameters instead of traditional iSSRT measures (**Figure 18A)**. The correlations between actual and fitted values increased from 0.25 to 0.53 (**Figure 18B)**. Cross-validated results confirmed this pattern, with PRAD parameters maintaining higher predictive power (adjusted R² = 15.3%; correlations 0.40) compared to iSSRT (adjusted R² = 4.6%; correlations 0.22) (Figure 18C-D).

**Figure 18.**
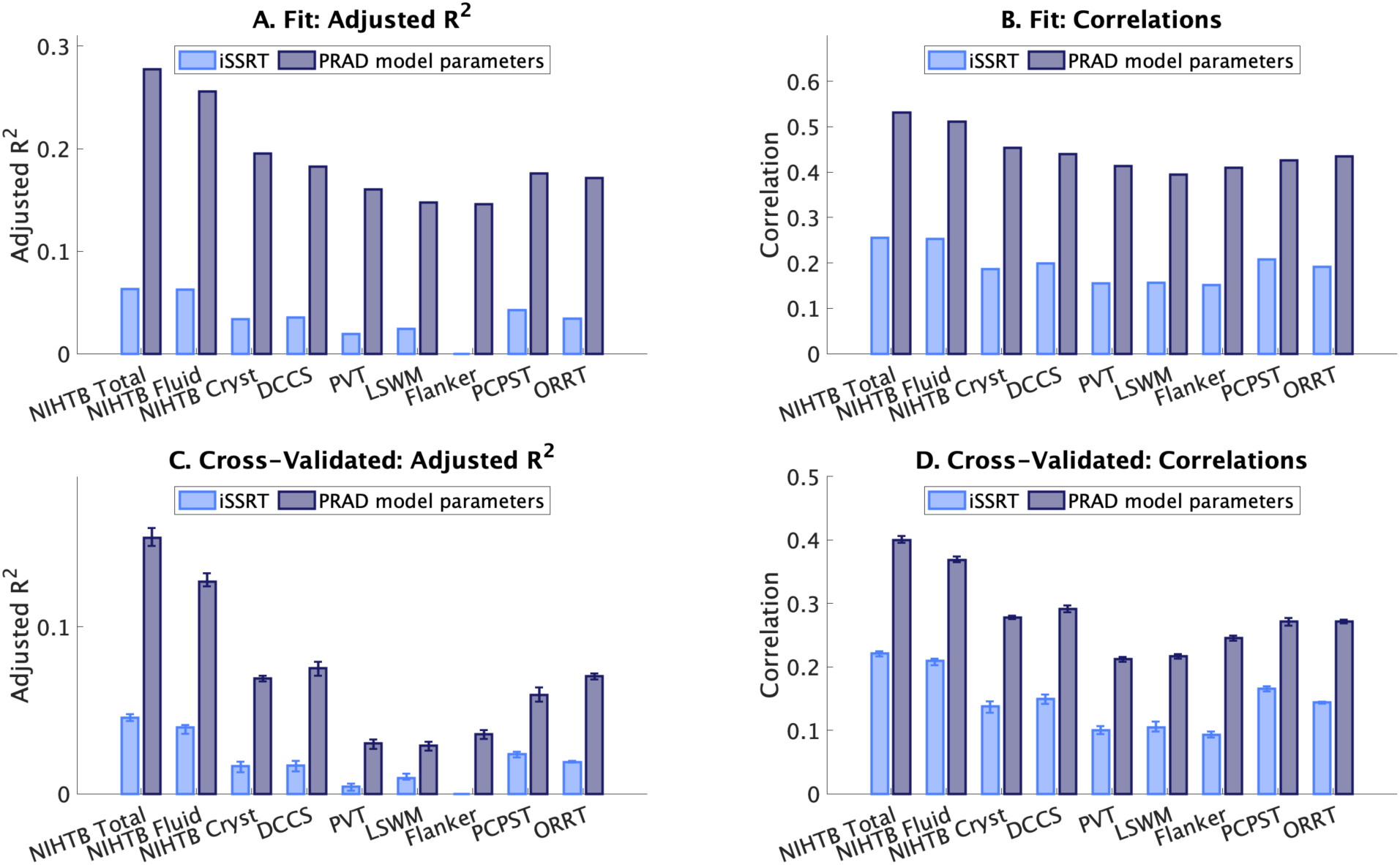
Predictive power of PRAD for NIH Toolbox cognitive tasks. Comparison of model fit and cross-validated performance using traditional SSRT (iSSRT) versus PRAD model parameters. Results show superior predictive power of PRAD for task-distal cognitive measures, with higher adjusted R² and correlations between predicted and actual values in both model fit and cross-validated analyses. Error bars show the 2.5th to 97.5th percentiles across 10-fold crossvalidation.

Similar improvements were observed for individual NIH cognitive toolbox tasks and subscores, including Flanker Inhibitory Control, Dimensional Change Card Sort, and Pattern Comparison Processing Speed tasks (**SI Table S18;** Figure 18). These findings suggest that PRAD parameters capture generalizable aspects of cognitive control that extend beyond the specific context of the stop-signal task, providing a more comprehensive characterization of individual differences in cognitive abilities.

The NIH Toolbox is an integral component within the Research Domain Criteria (RDoC) framework, utilized for assessing a wide range of cognitive functions across various disorders and developmental stages. This comprehensive toolset enables researchers to bridge cognitive performance with underlying neural and psychological mechanisms, making the PRAD model’s predictive power particularly valuable. Our analysis that parameters derived from the PRAD model outperformed traditional SSRT metrics for explaining individual differences in tasks measuring executive function, attention, processing speed, language, and learning/memory, lend substantial external validity to the PRAD model. By aligning PRAD with tasks from the NIH Toolbox, our findings highlight the model’s capacity to capture cognitive processes that are foundational across multiple domains of function and dysfunction. This also sets the stage for future research aimed at integrating cognitive modeling with clinical diagnostics and therapeutic interventions, guided by the principles of the RDoC framework.

### Clinical Relevance

We examined how key PRAD parameters and traditional SSRT measures differed between specific clinical subgroups based on CBCL evaluations and all other individuals. Groups were determined based on a CBCL t-score cutoff of 64, identifying children with attention disorders, rule-breaking, aggression, depression, ADHD, oppositional disorder, and conduct disorder. SSRT shows a significant difference (with effect size > 0.2) only for ADHD and conduct disorder, while the PRAD proactive factor showed significant differences (with effect size > 0.2) for attention, rule-breaking, aggression, ADHD, and conduct disorder, while PRAD go drift rate showed differences for all of these subgroups. Effect sizes for the drift rate were the largest, followed by the proactive factor, and lowest for SSRT (**Figure 19A**). This points towards the emergence of the PRAD drift rate as a transdiagnostic measure, and PRAD proactive control as a stronger measure for differentiating externalizing disorders compared to SSRT.

**Figure 19.**
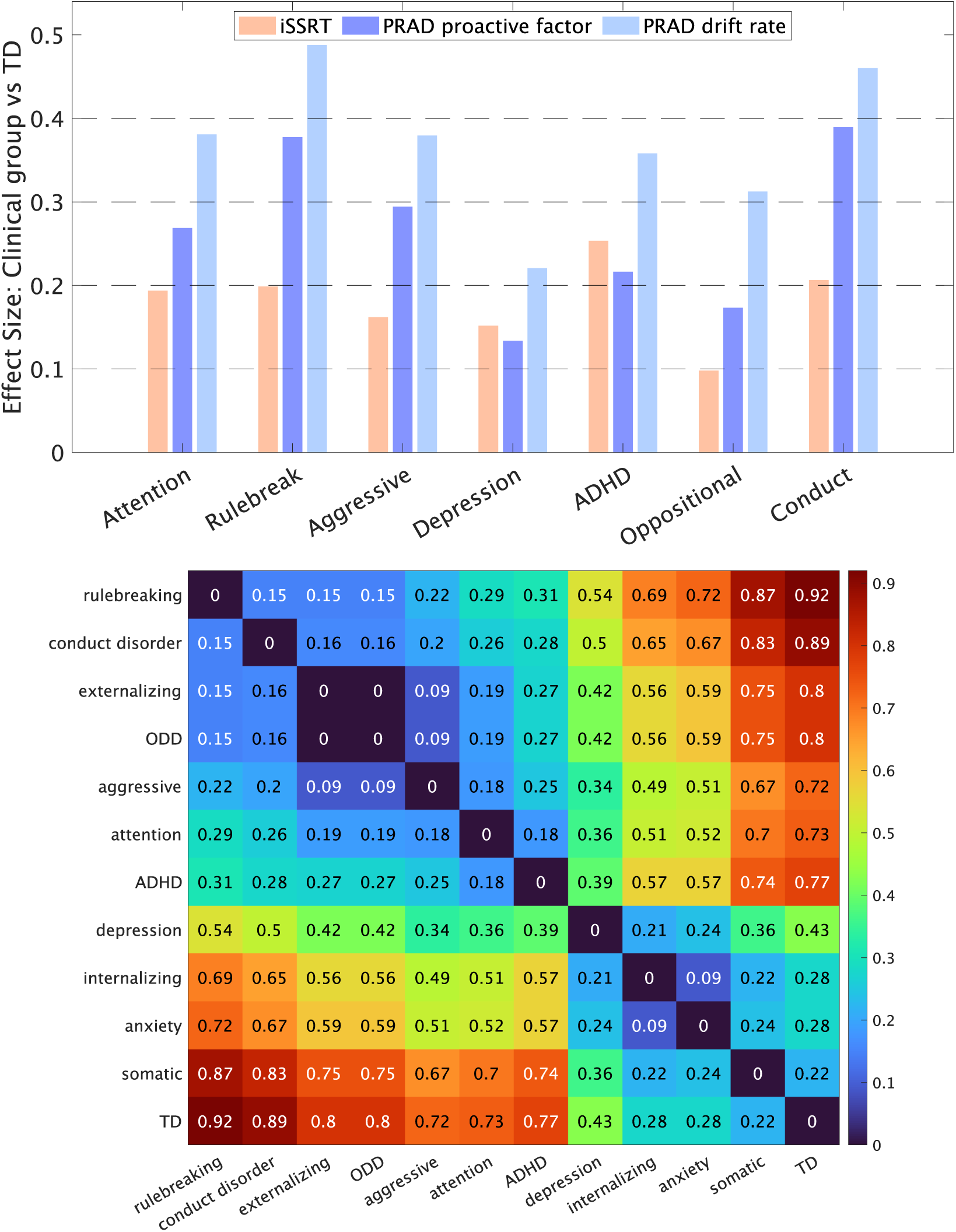
Clinical relevance of PRAD model. (A) The bar charts show effects sizes for the traditional SSRT measure (iSSRT), and key PRAD measures related to proactivity and drift rates, for the effect between specific clinical groups and all other individuals. The clinical groups include attention disorders, rule-breaking, aggression, depression, ADHD, oppositional disorders, and conduct disorders, based on CBCL. In each case other than ADHD, iSSRT effect sizes are less than 0.2 (small). PRAD measures show small to medium (0.2 to 0.5) effect sizes for each of these clinical groups. (B) Pairwise distance between clinical subgroups based on PRAD parameters

Further, we looked at the mean normalized PRAD parameters values for various clinical subgroups (attention disorders, rule-breaking, aggression, depression, ADHD, oppositional disorder, conduct disorder, externalizing, internalizing, somatic, and anxiety disorders) based on the CBCL, as well as typically developing children (as measured by not having membership to any of these 11 clinical subgroups). Computing a pairwise distance between the subgroups based on PRAD parameters yields a hierarchy (**Figure 19B**), with externalizing and internalizing symptoms and diagnosis clustering together, and reveals some interesting patterns, such as externalizing subgroups showing a higher level of separation from TD compared to internalizing, and depression falling somewhere in the middle, slightly distinct from other internalizing subgroups. Importantly, this tells us that the PRAD identifies aspects within the inhibitory control task that are relevant to graded differences based on patterns of clinical symptoms.

In addition, the key factors tackled in our work – debiased SSRT measurements for extreme performers, debiased inferences for non-typical or under-represented subgroups, heterogeneity in intraindividual variability, and heterogeneity in proactive, top-down, and bottom-up regulatory processes contributing to modulation of inhibitory control may become even more important in clinical and neurodiverse populations. The distinction between reliance on proactive versus reactive control mechanisms, as illuminated by the PRAD model, offers a refined lens through which maladaptive behaviors and transdiagnostic symptoms can be understood. Individual differences in these control strategies could account for the wide variability in cognitive performance and behavioral outcomes observed across and within psychiatric disorders. For instance, children with ADHD have been shown to have impaired reactive but not proactive inhibition (Pani et al., 2013; van Hulst et al., 2018). Other studies have also shown that dysfunctional impulsivity is linked to impairments in reactive but not proactive inhibition (Castro-Meneses et al., 2015). This understanding holds significant promise for tailoring interventions to target specific cognitive control deficits, moving towards a more personalized approach in clinical practice. For example, an overreliance on reactive control and a diminished capacity for proactive control may contribute to impulsive behaviors and difficulty with goaldirected planning, which are common features of many psychiatric disorders. By considering individual differences in these control mechanisms, PRAD can help to elucidate the cognitive processes that may underlie common symptoms across different diagnostic categories, in line with the RDoC approach.

### Neural Relevance

To further validate PRAD’s dissociation of reactive and proactive control mechanisms, we examined their neural correlates at the single-trial level. We examined within-subject trial-level relationships between PRAD dynamic parameters and brain fMRI activity. Our analysis focused on core measures of reactive control (trial-level SSRT) and proactive control (trial-level probability of proactivity, and proactive delaying). The results revealed distinct neural signatures corresponding to the dynamics of reactive (SSRT) and proactive (probability of proactivity; proactive delaying) mechanisms within subjects (Figure 20 A-C). Network-level (Shirer, Ryali,

**Figure 20.**
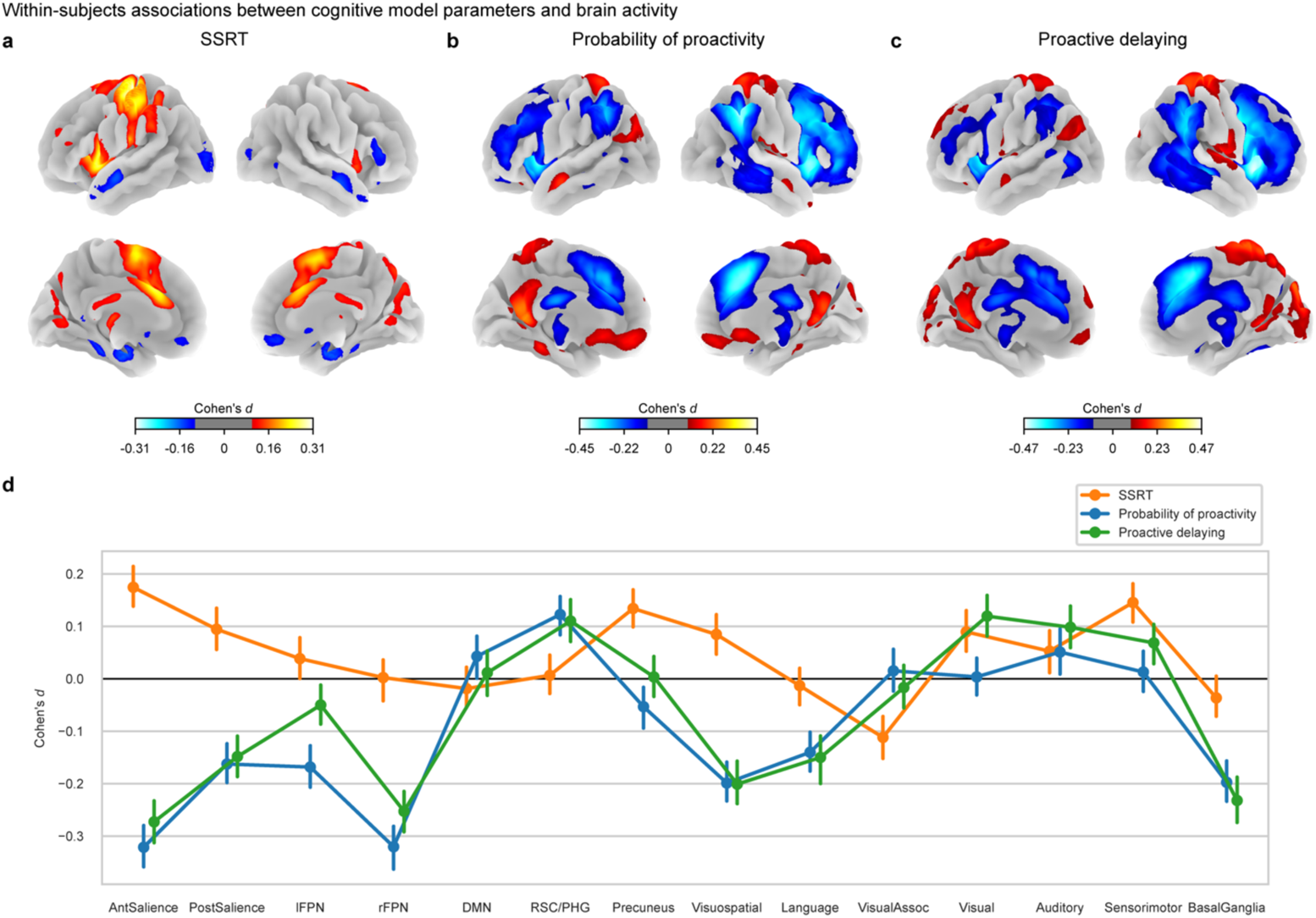
Neural relevance of the PRAD model: Within-subjects analysis with PRAD dynamic parameters reveals distinct neural correlates for reactive and proactive mechanisms. (a-c) Whole-brain Cohen’s d maps showing associations between trial-by-trial brain activity and cognitive model parameters (SSRT, probability of proactivity, and proactive delaying). Thresholded at Cohen’s d ≥ 0.1. SSRT and probability of proactivity N = 4469; proactive delaying N = 4176. (d) Network-level comparison of effect sizes based on the Shirer parcellation(Shirer, Ryali, Rykhlevskaia, Menon, & Greicius, 2012). Error bars show 99% confidence intervals computed using bootstrap resampling.

Rykhlevskaia, Menon, & Greicius, 2012) visualization of the within-subjects associations (Figure 20 **D**) revealed that SSRT showed positive associations in the posterior salience, precuneus, visuospatial, auditory, and sensorimotor networks (all PFDR < 0.01). Probability of proactivity showed negative associations in the salience and frontoparietal networks, and positive associations in the default mode network (all PFDR < 0.01), while proactive delaying showed predominantly negative associations, notably in the salience and frontoparietal networks (all PFDR < 0.01), but positive associations in the retrosplenial cortex and parahippocampal gyrus nodes of the ventral default mode network (PFDR < 0.01). These dissociable neural correlates provide strong validation for PRAD’s separation of reactive and proactive control mechanisms, supporting the model’s neural cognitive architecture. A comprehensive analysis of neural correlates of the PRAD model parameters and demonstration of non-ergodicity in the neurocognitive dynamics of inhibitory control using the PRAD model can be found in a separate paper (Mistry, Branigan, Gao, Cai, & Menon, 2024).

### Relevance to capturing exposomic factors

We examined the correlation of key PRAD parameters (factors relating to proactive control and basic reactive control, and go process drift rate), as well as traditional SSRT measures to exposomic factors including SES (family income category), ALE (adverse life events), and BPM (Behavioral problem monitoring based on teacher inputs). Figure 21 shows how the parameter values change over 4 quantiles of these exposomic factors, along with the underlying correlation. PRAD measures, especially go process drift (*r* 0.1 to 0.21; all p < 0.0001) and factor 1 pertaining to proactive control (r 0.08 to 0.14; all p < 0.0001) generally show stronger correlations compared to SSRT (r 0.03 to 0.1; all p <= 0.011). Lower income (0.07 to 0.2; all p < 0.0001), higher adverse life events (0.03 to 0.1; all p <= 0.011), and higher teacher reported problems (0.1 to 0.21; all p < 0.0001), correspond to lower proactive control, weaker go process drift, weaker reactive control, and higher SSRT.

**Figure 21.**
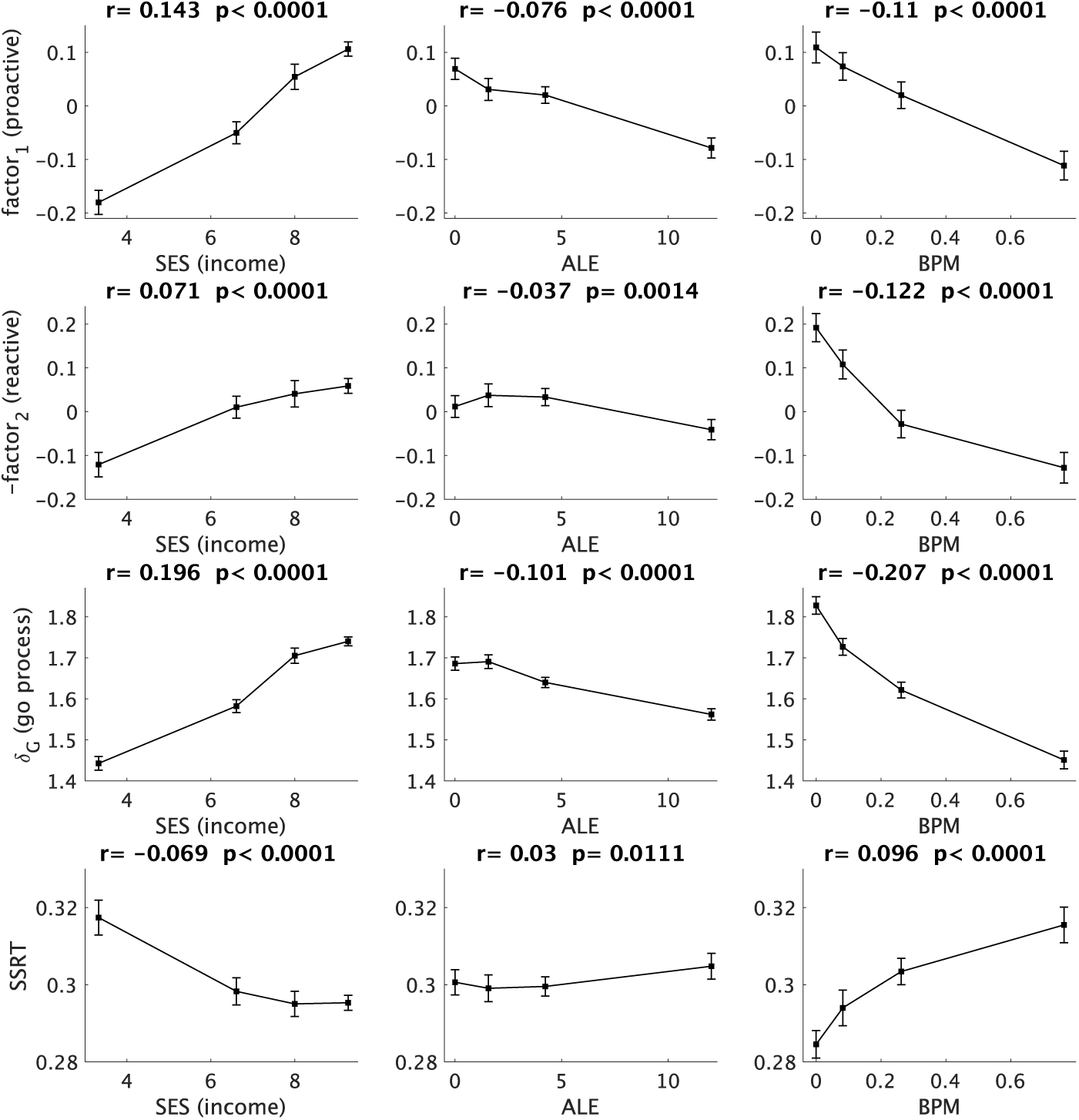
Exposomic relevance of the PRAD model.

Combined with the findings that PRAD reduces typicality bias for subpopulations such as children from lower income families, children with adverse life events, and behavioral problems, these findings demonstrate the added value of using our approach when examining the role of exposomic factors.

### Model Reliability and Sensitivity Analysis

#### Reliability of model measurements

Behavioral and traditional SSRT measures from the SST task have previously been shown to have moderate (ICCs 0.34 to 0.57) levels of reliability (Hedge, Powell, & Sumner, 2018), at least partially attributed to state-dependence of some aspects of the task. Further, these estimates do not typically include a measure of reliability for second-order observed effects, such as the level of within-individual correlations between observed variables and experimental contingencies, which are expected to be even lower.

We first assessed the reliability in behavioral observations across the two runs of SST, yielding *behavioral ICCs* of 0.12 (SFR), 0.57 (go-omissions), 0.85 (RT), -0.01 (correlation between SFR and number of trials since the last stop signal), and 0.10 (correlation between SFR and SSD), with a mean of 0.33 across these measures.

While our primary results are based on jointly fitting data from two separate runs of the SST task, we also fit each run separately and assessed the reliability in PRAD measures across the two runs. *PRAD ICCs* ranged from 0.01 to 0.72, with a mean of 0.34 across the model parameters (**SI Table S19**). Overall, the reliability of PRAD model parameters was within the same range as demonstrated by behavioral and observed measures in the task.

We note here that moderate reliability in cognitive tasks like the SST has previously been attributed to low levels of heterogeneity (the ‘reliability paradox’) in the context of measured task behaviors and state-dependence of these measures (Hedge et al., 2018), rather than measurement noise. This is in line with our results, where the model parameter reliabilities are similar to the reliabilities of different observed measures. Further, the PRAD parameters related to basic go and reactive stopping processes, as well as average dynamic measures of proactive delayed responding (PDR), show higher reliability than the model measured SSRT. Further, the PRAD measures enable dissociating heterogeneity across multiple dimensions, and PRAD measures related to PDR and AMS such as baseline proactivity (θ_0_, 0.69), error-monitoring sensitivity (θ_1_, 1.21), and attention-based adaptivity (γ_1_, 1.29) show higher coefficients of variation compared to SSRT (0.57), which in turn is higher than observed measures like RT (0.16) and SFR (0.18).

These results demonstrate that the reliability of model parameters seem to be limited only by the reliability of task behavior itself, but they provide a more dissociated process perspective of behavior, providing a more nuanced perspective of heterogeneity and individual differences, as well as identify which processes may contribute more strongly to state-dependence (e.g. AMS), without any meaningful loss of reliability when compared to traditional or behavioral measures. Furthermore, with more data collection efforts focused on large datasets with multiple rounds of measurement, the variability in state-dependent processes like AMS over multiple occasions may itself be a potentially relevant diagnostic factor.

### Sensitivity analysis

We performed a comprehensive set of sensitivity analysis, including testing versions of PRAD that were fully generative (PRAD-g vs discriminative drift rate estimation), models with an alternate set of priors for non-decision time (PRAD-n), and subsampling participants based on MCMC sampling patterns. **SITableS19** shows that ICCs between PRAD and PRAD-g parameters are 0.9-1.0 and **SI Table S20** shows that all key results highlighted in this paper are robust to these sensitivity analyses. Details can be found in the **Methods** section.

## Conclusion

We developed and validated a novel computational model of Proactive, Reactive, and Attentional Dynamics (PRAD), that characterizes latent proactive, reactive, and attentional components underlying inhibitory control. We leveraged a very large dataset (N > 7,500) of children ages 9-10 from the NIH ABCD study, which allowed us to probe distinct sources of response intraindividual variability in ways that were previously not possible. The ABCD study provides a unique opportunity to investigate cognitive processes at an unprecedented scale, with a sample size that far exceeds most previous studies in the field. PRAD demonstrates robustness to violations of context and stochastic independence, as well as extreme patterns of behavior, both of which are a limitation for most extant models. PRAD model parameters also provide a nuanced and integrated perspective on the multiple neurocognitive processes that combine to produce inhibitory control behavior. We delineate specific mechanisms of proactive control and attention modulation, demonstrating their interaction and ability to compensate for weak reactive inhibitory control. The strong influence of proactive control processes on behavioral variability suggests that factors previously attributed to reactive inhibitory control failures may reflect breakdowns in proactive control. PRAD as a sophisticated, multicomponent, model offers a dynamic framework for precisely characterizing goal-directed behaviors, meaningfully delineating individual differences in cognitive control processes and their functional consequences, and making theoretical advances in our understanding of cognitive control. PRAD also reduces biases in latent measures, reduces typicality biases, and provides improved fits to a wider range of populations and behaviors. These advances will be critical for examining dynamic neurocognitive mechanisms of inhibitory control in diverse populations.

We have also shown the relevance of PRAD measures to explain individual differences in performance across a range of executive function, attention, processing speed, language, and learning/memory tasks compared to conventional models, to understand and dissociate the neural dynamics underlying proactive and reactive control mechanisms, to identifying differences in clinical subgroups, and to identifying associations between exposomic factors and differences in cognitive control processes. By leveraging computational models like PRAD, we anticipate advancing our understanding of the mechanistic accounts of cognitive disruptions associated with psychopathology. PRAD’s process specificity and explanatory power highlight its potential for elucidating control deficits in psychopathology and informing individualized interventions.

The availability of dynamic trial-level latent cognitive parameters allows for holistic investigation of nonergodic neural processes involved in inhibitory control, supporting the RDoC framework.

More broadly, PRAD exemplifies the utility of sophisticated computational approaches in unraveling the complex dynamics of cognition. It paves the way for developing models that can better capture heterogeneity in cognitive processes across populations and advancing our understanding of mechanisms underlying adaptive and maladaptive behavior.

## Supplementary Information: Methods

### Participants and inclusion criteria

The present study used a sample of N = 7787 from the Adolescent Brain Cognitive Development (ABCD) study. The ABCD study is an ongoing, longitudinal study within the United States that follows a nationally representative sample of children aged 9–10 at baseline (B. J. Casey et al., 2018; Garavan et al., 2018). Data were from the baseline visit of the ABCD study (Volkow et al., 2018) (Collection #2573). We report how we determined our sample size, all data exclusions, and all measures in the study. 9355 participants had two complete runs of the SST task and had data that were successfully fit with the PRAD model, including checks for convergence. We then excluded participants who had a glitch reported in the task presentation (N = 257) and those who had left vs right choice accuracies less than chance level (N = 164). Then, we excluded siblings by randomly keeping one member from each family (using the genetic_paired_subjectid variables from gen_y_pihat; 𝑁 = 1147 excluded). Applying these inclusion criteria left us with a sample of 𝑁 = 7787. See **SI Table S21** for participant demographics.

### Stop-Signal Task

Participants completed the SST (Logan, Schachar, & Tannock, 1997) task during fMRI acquisition. Left and right facing arrows were presented serially as “go” stimuli. Participants indicated the direction of the arrows using a button box and were instructed to respond as quickly and accurately as possible, but to withhold their response on a small subset of trials (“stop” trials) when they saw an upward facing arrow (the “stop” signal), which only appeared after a brief delay (stop signal delay – SSD). Participants completed two runs of 180 trials each, with each run including 30 “stop” trials and 150 “go” trials. On stop trials, the time delay between the “go” and “stop” signals (SSD) was dynamically adjusted by 50 milliseconds increments – increasing after successful stopping and decreasing after unsuccessful stopping, targeted to modulate the difficulty levels so that each participant would be able to successfully inhibit their responses approximately 50% of the time.

### Estimating behavioral patterns

#### Violations of stochastic independence

Based on simulations (Band et al., 2003), a slope of 3.5 corresponds to estimated correlation of 0.2 between RT and SSRT, and we use this benchmark.

**High skew.**benchmarked by inferring the mean / SD of the exponential component of an ex-Gaussian distribution fit to reaction times, being greater than 150ms (F. Verbruggen et al., 2019).

**High slowdown.** benchmarked by fitting a linear slope of RT over trials that results in RT increasing by more than 50% from the first to last trials, or a slowing factor of 1.5 (Verbruggen et al., 2013).

**High RT variability.** This was measured by first inferring the SD of the Gaussian component of an ex-Gaussian distribution fit to reaction times. Then this measure of SD across all participants was fit to a mixture of two Gaussian distributions, and high variability was identified as participants with SD higher than the mean of the larger of the two mixture distributions.

### Computational Modeling – additional details

**Hierarchical Bayesian implementation.** The following MCMC sampling hyperparameters were used: Total 4 chains with 30000 samples each. For each chain, we had a burn-in of 10000 samples, and a thinning factor of 10 was used to record 2000 of the remaining 20000 samples, based on which final inferences were made. The Bayesian priors used in the model are described in **SI Table S22**. Convergence was assessed using R-hat <= 1.1 across all chains. For some participants where convergence across 4 chains was difficult to achieve, we assessed and used only convergent samples based on whether 3 or 2 chains converged. Model inferences for 4195 participants (54%) included convergent samples based on 4 chains, for 2409 (31%) included convergent samples based on 3 chains, and for 1183 (15%) included convergent samples based on 2 chains. To ensure that sampling hyperparameters did not have an effect on the results, we ran a sensitivity analysis (**SI Table S20**) to ensure that all major results replicate, when considering only participants based on selective sampling hyperparameters (convergent samples based on 4 chains; convergent samples based on 3 or more chains). For model fitting, reaction-times less than 0.05s and greater than 1s (the experimental response window) were censored.

### Parameter Recovery

Additional analysis demonstrated that the model shows strong parameter recovery (**SI Figure S1**). To test parameter recovery, we sampled 750 combinations of parameters inferred from actual data, then generated new simulated data (RT, stopping success, choice accuracy, at a trial level) using the PRAD model and these combinations of parameters. Finally, we fit this simulated data using the PRAD model and compared the inferred parameters (recovered) to the parameters used to simulate the data. Parameter recovery is not an assessment of the validity of the PRAD model or its assumptions, nor a measure of effectiveness of the Bayesian methods used to make inferences. It does provide a way to check the implementation of the model, diagnose any potential identifiability issues (Matzke, Logan, & Heathcote, 2020b), and understand the adequacy of the ABCD SST experimental designs for making useful model inferences.

**Sensitivity analysis - PRAD-n model.** Note that 𝜏_-_ is a fixed component of non-decision time, and not strictly comparable to non-decision times in standard DDM models. The priors on 𝜏_-_ in the primary analysis are bounded at 0.1s, smaller than typically assumed for parameters that represent the full non-decision time. For 47% of the participants, the shortest RT is less than 0.1s, so this upper bound would not make an impact (𝜏_-_ < shortest RT) for these participants. The posterior mean and median inferred values of 𝜏_-_ are 0.05s. Recent modeling of the SST process (Weigard et al., 2023), which does *not* consider a separation of non-decision time into fixed and strategic proactive components found overall mean non-decision time of 0.137s in a similar sample. Our selected priors for a partial component of this are thus in line with previous observations. This choice of priors also results in reliable parameter recovery. To ensure that our results are not dependent on this choice of priors for non-decision time, we ran a sensitivity analysis allowing the upper bound on the fixed component 𝜏_-_ of non-decision time to extend up to 0.3s (**SI Table S20,** NDT prior), which shows that all major results replicate regardless of this choice of prior.

**Sensitivity analysis - PRAD-g model.** In the PRAD model, the dynamic drift rate on each trial depends on the duration of the go stimulus (𝑠𝑡𝑖𝑚_!_). On go trials, since this is confounded with the go RT, it can be argued that this formulation is discriminative rather than a strictly generative model. To test the robustness of PRAD, we also test a fully generative model (PRAD-g) where the dynamic drift rate is only applied on stop trials (where the duration of go stimulus is censored by the onset of the stop signal) and 𝑠𝑡𝑖𝑚_!_is not confounded with RT. On go trials, it is retained as 𝛿_"_ 𝕊_45,!_, similar to previous work (Weigard et al., 2023). **SI Table S20** (PRAD-g) shows that all major results replicate for this model.

**Control Models – RVM.** Random variability model is a simplification of the PRAD model without the dynamic hierarchical components and is equivalent to a full Bayesian implementation of the traditional horse-race model(Band et al., 2003), but with the addition of allowing SSRT to vary randomly across trials, based on the parameters of a stopping drift diffusion process. It can be considered a nested version of PRAD with the following constraints applied, plus a change in some priors (**SI table S22):** Constraints: 𝛼_-#_ = 𝛼_-3_, 𝛿_-,!_ = 𝛿_"_ 𝕊_45,!_, 𝜆_!_ = 0, 𝜌_!_ = 0, 𝛽_/,!_ = 0.5.

**Control Models – FSM:** Fixed SSRT model is a further simplification of RVM, where the stopping process is not explicitly modeled, but a constant SSRT value is inferred for each individual which applies to all trials.

### Statistical Testing

Unless stated otherwise, correlations refer to Pearson’s linear correlation coefficient, p-values are based on two-tailed tests.

**A note on effect sizes.** To put some of the effect sizes in this paper in context, note that at our sample size > 7000, there are considerable differences between coefficients of 0.02, 0.05, 0.1, or 0.2. Using random conclusions estimators (Davis-Stober, Dana, Kellen, McMullin, & Bonifay, 2024) the ratio of MSE of a random conclusions estimator to that using the estimator in question with effect sizes of 0.02, 0.05, 0.1, and 0.2 at a sample size of N ∼ 7000 would correspond to ratios obtained from effect sizes of 0.17, 0.42, 0.84, and 1.67 using a sample size ∼ 100.

**Factor Analysis.** Factor analysis was carried out using the lavaan(Rosseel, 2012) package in R(Team, 2020) with the following choice of SEM (structural equation model) hyperparameters: (rotation = "oblimin", estimator = "ML", likelihood = "normal", auto.var = TRUE, auto.efa = TRUE). To compare and evaluate the adequacy of factor analysis models, we used the following criteria: CFI (comparative fit index; threshold 0.95), TLI (Tucker-Lewis index; threshold 0.95), and RMSEA (threshold < 0.08).

## Software

Data were processed and analyzed using R (Team, 2020), MATLAB, and JAGS (Plummer, 2003) (version 4.3.0).

## Acknowledgements

Funding: This work was supported by: The National Institutes of Health (MH121069, MH124816) The National Science Foundation (2024856) The Stanford Maternal and Child Health Research Institute.

## Data availability

Data used in the preparation of this article were obtained from the Adolescent Brain Cognitive

Development^SM^ (ABCD) Study (https://abcdstudy.org), held in the NIMH Data Archive (NDA). This is a multisite, longitudinal study designed to recruit more than 10,000 children age 9-10 and follow them over 10 years into early adulthood. The ABCD Study® is supported by the National Institutes of Health and additional federal partners under award numbers U01DA041048, U01DA050989, U01DA051016, U01DA041022, U01DA051018, U01DA051037, U01DA050987, U01DA041174, U01DA041106, U01DA041117, U01DA041028, U01DA041134, U01DA050988, U01DA051039, U01DA041156, U01DA041025, U01DA041120, U01DA051038, U01DA041148, U01DA041093, U01DA041089, U24DA041123, U24DA041147. A full list of supporters is available at https://abcdstudy.org/federal-partners.html. A listing of participating sites and a complete listing of the study investigators can be found at https://abcdstudy.org/consortium_members/. ABCD consortium investigators designed and implemented the study and/or provided data but did not necessarily participate in the analysis or writing of this report. This manuscript reflects the views of the authors and may not reflect the opinions or views of the NIH or ABCD consortium investigators. These data are publicly shared with eligible researchers.

## Code availability

All code used in this study will be made available upon publication (https://github.com/scsnl).

## Preregistration

This study was not preregistered.

