## Supplementary material for "Proactive, Reactive, and Attentional Dynamics: An Integrative Model of Cognitive Control": SI

Percy K. Mistry et al.

Corresponding authors: Percy K. Mistry, Ph.D. & Vinod Menon, Ph.D.


**This PDF file includes:**

Figures SI1 to SI5
Tables S1 to S22

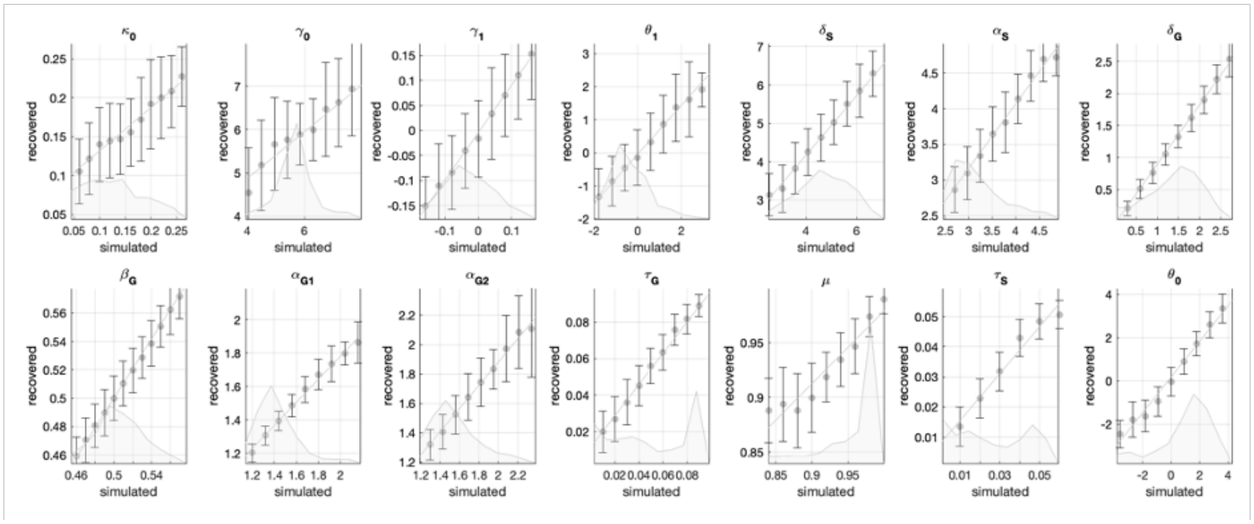

**Figure S1. Parameter Recovery Study.** Figures show the recovered parameters vs binned simulated values, tested for 750 combinations of the 14 PRAD model parameters. The distributions for the simulated values are also shown. Error bars show standard deviations.

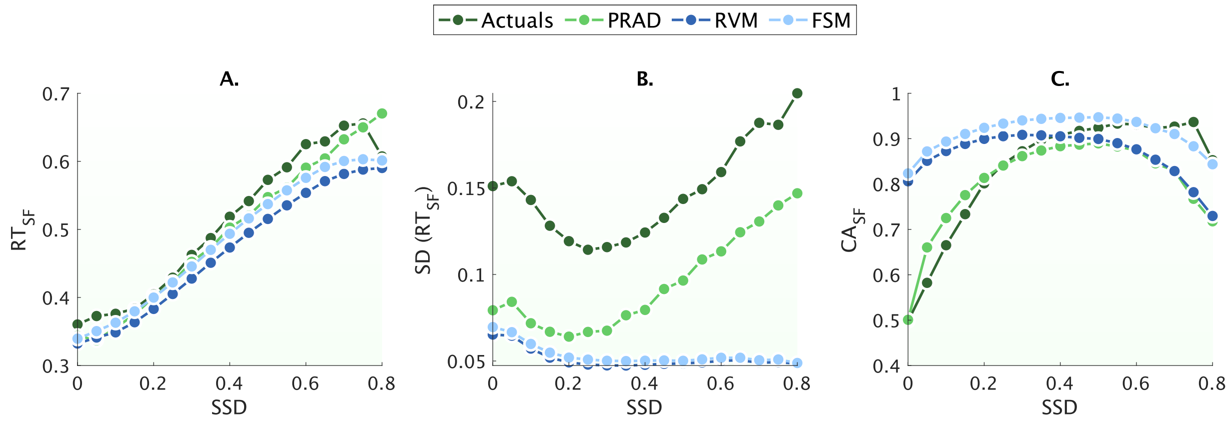

**Figure S2. Model fit comparisons.** Comparing aggregate patterns of actual data with predictions from PRAD and control models (RVM and FSM). Aggregate behavior patterns for (A) RT on stop-failure, (B) SD of RT, and (C) choice accuracies on stop failure across different stop-signal delays (SSD). Dark green – actuals, Light green – PRAD, dark blue – RVM, light blue – FSM. RVM = Random Variability control model; FSM = Fixed SSRT control model.

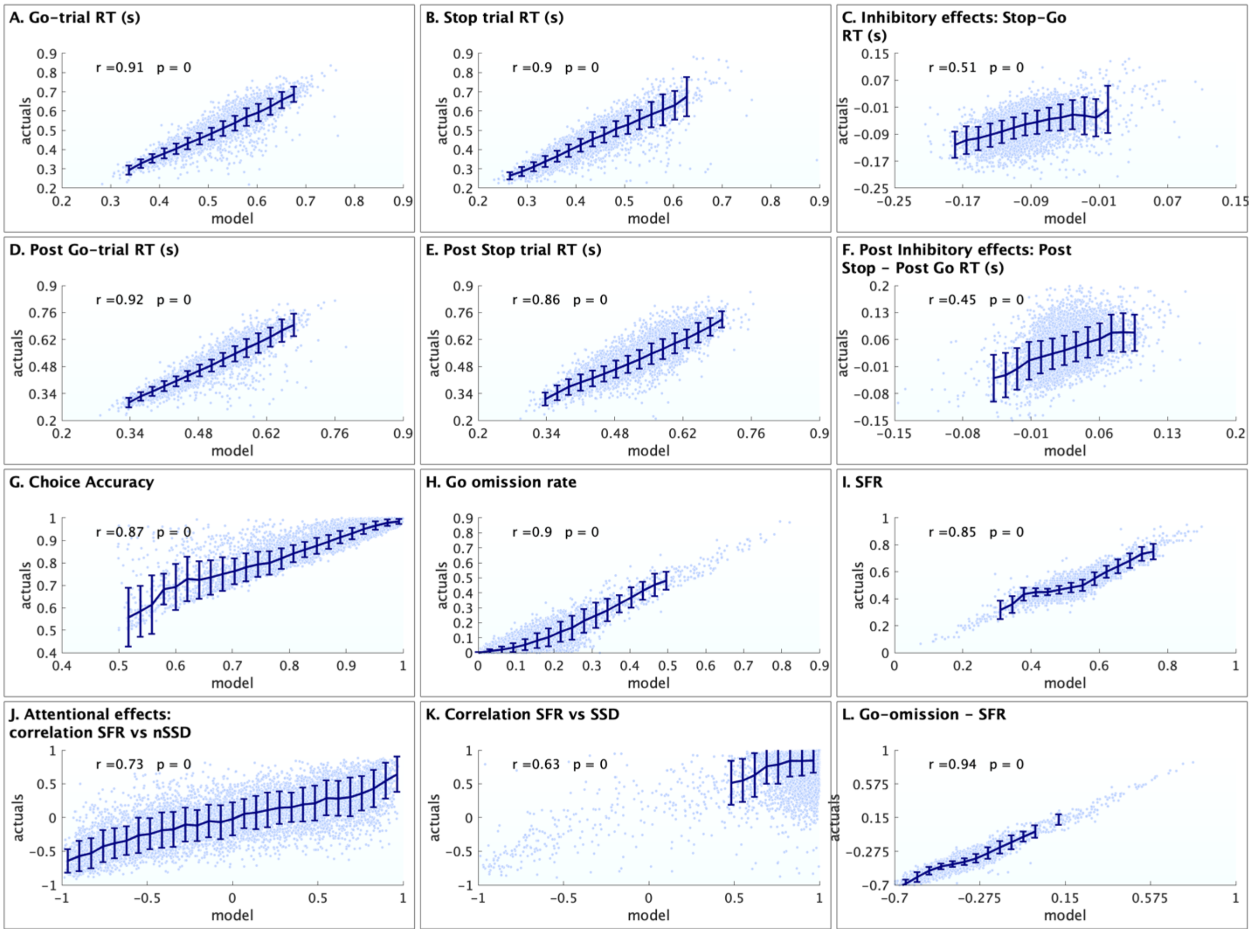

**Figure S3. Individual level fits to data for the PRAD model.** Actuals versus model inferred values for observed measures. (**A-F**) RT related measures: Go-trial RT, stop trial RT, RT difference between stop and go trials, post go-trial RTs, post stop-trial RTs, difference between post stop and post go RTs. (**G-I**) Choice accuracy, go omission rates, stop-failure rates. **(J-K)** Correlations between stop-failure rate and number of trials since last stop signal (nSSD), stop-failure rate and SSD. **(L)** Difference between go-omission rate and stop-failure rate. Error bars reflect standard deviation.

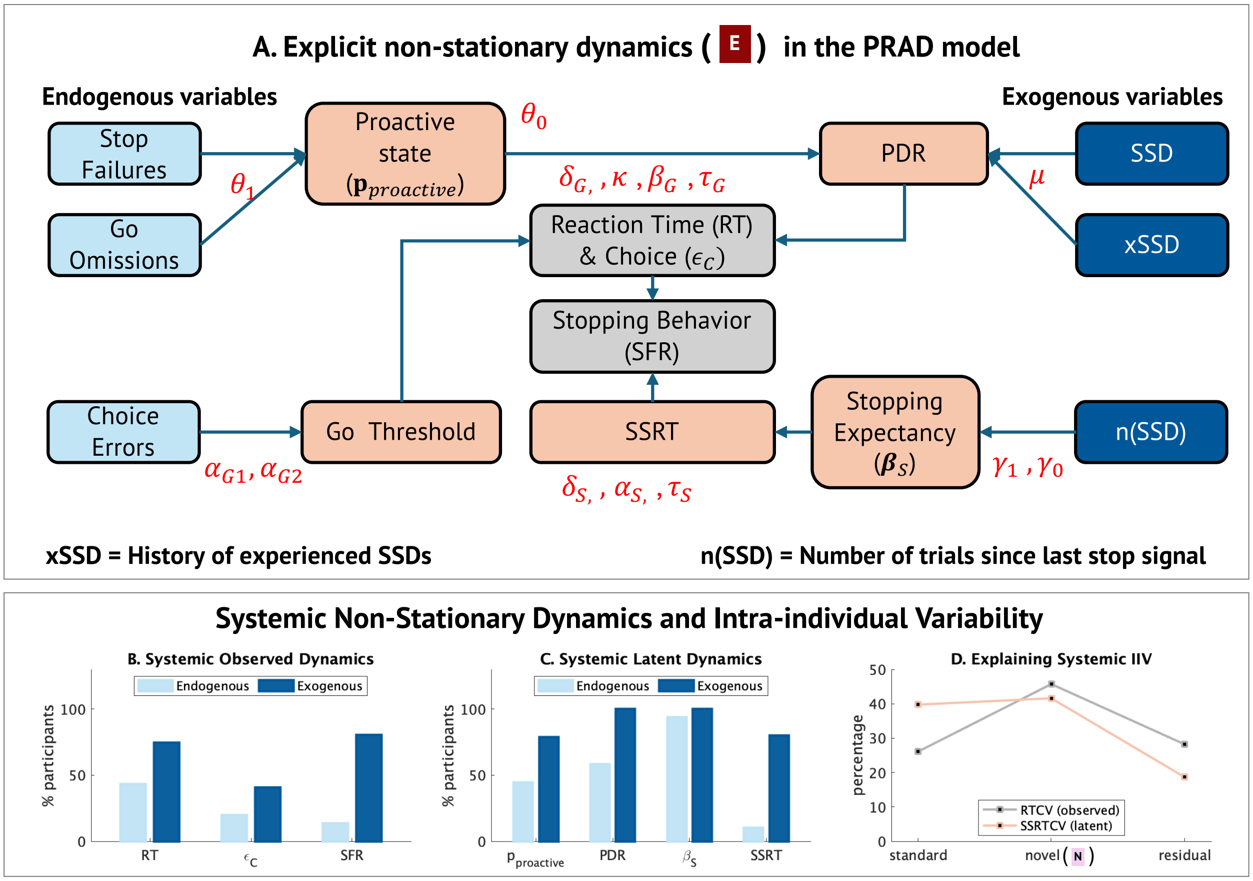

**Figure S4. Non-stationary dynamic and intra-individual variability. (A)** Model pathways for non-stationary dynamics. The PRAD model incorporates latent dynamics that respond to endogenous (aspects that are manifested internally, such as internal cognitive states and performance or error monitoring that is based on implicit or internal assessments) and exogenous (aspects that trigger responses based on explicit or experimental contingencies, such as the experimental stop-signal delays or trial sequence types) variables. Trait measures may govern the interaction of such endogenous and exogenous variables with latent processes, giving rise to non-stationary dynamics. **(B-C)** Proportion of participants who show significant within-subject correlations between (**B**) observed measures and endogenous or exogenous variables; (**C**) latent measures and endogenous or exogenous variables. **(D)** Proportion of variance in IIV (measured by RTCV and SSRTCV) that is explained by standard model parameters, novel model parameters, and residuals.

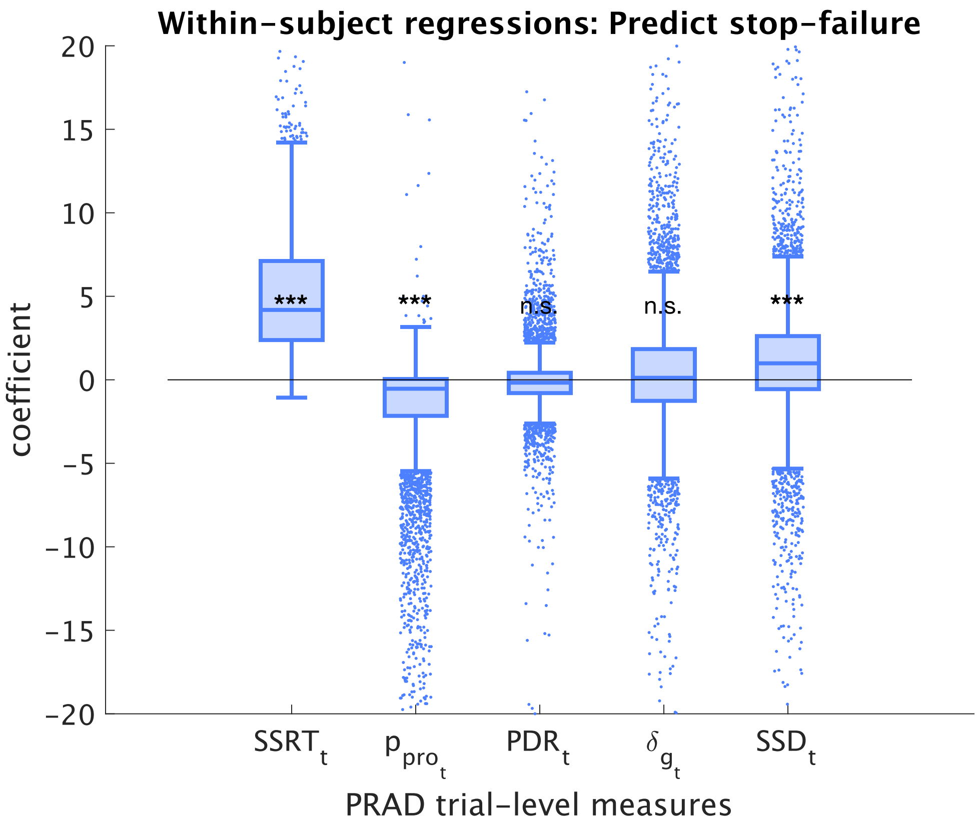

**Figure S5. Within-subject regressions predicting stop failure**. Distribution of standardized coefficients across individuals, showing that higher SSRT and lower probability of proactivity are the strongest contributors to stop-failure rates within individuals.

### Supplementary Tables

#### Table S1. Key PRAD model features.

| Model features | PRAD |
|---|---|
| Process model of stopping | Yes |
| Attentional effects | Yes |
| Modulation of stopping expectancy | Yes |
| Cognitive state switching | Yes |
| Error / performance monitoring | Yes |
| Proactive (strategic) delayed responding | Yes |
| Stimulus duration effects | Yes |
| Adaptive decision threshold | Yes |
| Systemic account of IIV | Yes |
| Assumptions of context / stochastic independence | Not required |
| Trial level SSRT estimates | Yes |
| Trial level measures of proactivity and attention | Yes |
| Accounting for trigger failures | Implicitly |

### Table S2. Summary of key model parameters

| | Interpretation | Process | Dynamic | Type | Mean | SD | 95% CI | |
|---|---|---|---|---|---|---|---|---|
| $\delta_G$ | Go process drift rate baseline | GO | | | 1.629 | 0.627 | 0.279 | 2.79 |
| $\alpha_{G1}$ | Go process decision threshold | GO | | | 1.484 | 0.254 | 1.157 | 2.175 |
| $\alpha_{G2}$ | Go process threshold post-error | GO | Y | new | 1.618 | 0.302 | 1.176 | 2.384 |
| $\kappa_0$ | Duration sensitivity of go drift rate | GO | Y | new | 0.141 | 0.067 | 0.041 | 0.273 |
| $\beta_G$ | Choice bias | GO | | | 0.512 | 0.034 | 0.454 | 0.575 |
| $\delta_S$ | Stop process drift rate | Reactive | | | 5.017 | 1.118 | 2.691 | 7.046 |
| $\alpha_S$ | Stop process decision threshold | Reactive | | | 3.3 | 0.605 | 2.438 | 4.849 |
| $\theta_0$ | Proclivity for proactive state | PDR | Y | new | 1.637 | 1.918 | -3.854 | 4.262 |
| $\theta_1$ | Error modulation of cognitive state | PDR | Y | new | -0.135 | 1.123 | -2.008 | 2.776 |
| $\mu$ | Belief updating persistence | PDR | Y | new | 0.966 | 0.045 | 0.84 | 0.999 |
| $\gamma_1$ | Attention-based adaptivity | AMS | Y | new | -0.01 | 0.084 | -0.171 | 0.168 |
| $\gamma_0$ | Attention length | AMS | Y | new | 5.894 | 0.831 | 4.002 | 7.882 |
| $\tau_G$ | Go process non-decision time* | non-dec | | | 0.053 | 0.035 | 0.002 | 0.098 |
| $\tau_S$ | Stop process non-decision time | non-dec | | | 0.028 | 0.02 | 0.001 | 0.06 |

* partial fixed component of non-decision time

**Table S3. Key observed values and corresponding PRAD model posterior values.**

| Observed | Mean | SD | 95% CI | |
|---|---|---|---|---|
| Mean RT (mS) | 515 | 81 | 365 | 673 |
| Mean Go RT (mS) | 522 | 81 | 371 | 680 |
| Mean Stop-failure RT (mS) | 449 | 86 | 304 | 635 |
| Mean Go - Stop RT (mS) | 74 | 37 | -4 | 138 |
| Mean post-Go RT (mS) | 509 | 82 | 358 | 671 |
| Mean post-Stop RT (mS) | 544 | 85 | 380 | 708 |
| Mean post-Go - post-Stop RT (mS) | -35 | 40 | -117 | 43 |
| Stop failure rate (SFR) | 48% | 7% | 38% | 68% |
| Go omission rate | 7% | 10% | 0% | 38% |
| Choice errors on successful Go | 9% | 8% | 1% | 30% |
| Choice errors on unsuccessful Stop | 20% | 15% | 0% | 52% |
| Choice errors Go - Stop | -11% | 10% | -33% | 4% |
| Correlation (SFR, trials since last Stop signal) | 0 | 0.36 | -0.69 | 0.7 |
| Correlation (go-stop error, trials since last Stop signal) | 0.12 | 0.3 | -0.5 | 0.59 |

| PRAD posterior mean values | Mean | SD | 95% CI | |
|---|---|---|---|---|
| Mean RT (mS) | 500 | 63 | 377 | 617 |
| Mean Go RT (mS) | 515 | 63 | 391 | 631 |
| Mean Stop-failure RT (mS) | 420 | 69 | 302 | 568 |
| Mean Go - Stop RT (mS) | 95 | 30 | 27 | 144 |
| Mean post-Go RT (mS) | 495 | 64 | 374 | 616 |
| Mean post-Stop RT (mS) | 527 | 65 | 395 | 646 |
| Mean post-Go - post-Stop RT (mS) | -32 | 24 | -77 | 19 |
| Stop failure rate (SFR) | 50% | 7% | 39% | 68% |
| Go omission rate | 12% | 10% | 2% | 40% |
| Choice errors on successful Go | 12% | 10% | 1% | 39% |
| Choice errors on unsuccessful Stop | 21% | 11% | 5% | 46% |
| Choice errors Go - Stop | -9% | 6% | -23% | -1% |
| Correlation (SFR, trials since last Stop signal) | 0.02 | 0.53 | -0.88 | 0.87 |
| Correlation (go-stop error, trials since last Stop signal) | 0.27 | 0.26 | -0.41 | 0.66 |

**Table S4. PRAD improvement in aggregate fits. RMSE computed based on aggregate (binned) curves.**

| % reduction in RMSE based on aggregate curves | PRAD vs FSM | PRAD vs RVM |
|---|---|---|
| SFR by unique SSD values | 77% | 66% |
| SFR by unique nSSD values | 80% | 58% |
| Mean RT on stop failure by unique SSD values | 14% | 45% |
| SD RT on stop failure by unique SSD values | 45% | 46% |
| Mean RT on stop failure by unique nSSD values | 13% | 50% |
| Choice accuracy on stop failure by unique SSD values | 48% | 48% |
| Choice accuracy on stop failure by unique nSSD values | 61% | 51% |

### Table S5. Individual Model Fit – correlations and model comparison.

| Correlations | RVM | | FSM | | PRAD | | PRAD % Improvement | |
|---|---|---|---|---|---|---|---|---|
| | r | p | r | p | r | p | vs RVM | vs FSM |
| **RT related measures** | | | | | | | | |
| Go RT | 0.88 | 0.000 | 0.89 | 0.000 | 0.91 | 0.000 | 3 | 2 |
| Stop RT | 0.86 | 0.000 | 0.84 | 0.000 | 0.90 | 0.000 | 5 | 7 |
| Stop RT – Go RT | 0.40 | 0.000 | 0.30 | 0.000 | 0.51 | 0.000 | 28 | 70 |
| Post-Go RT | 0.87 | 0.000 | 0.89 | 0.000 | 0.92 | 0.000 | 6 | 3 |
| Post-Stop RT | 0.79 | 0.000 | 0.80 | 0.000 | 0.86 | 0.000 | 9 | 7 |
| Post-Stop RT–Post-Go RT | 0.02 | 0.077 | 0.03 | 0.014 | 0.45 | 0.000 | 2,150 | 1,400 |
| Within-sub SD RT | -0.09 | 0.000 | -0.12 | 0.000 | 0.13 | 0.000 | 244 | 208 |
| **Error related measures** | | | | | | | | |
| SFR | 0.79 | 0.000 | 0.65 | 0.000 | 0.85 | 0.000 | 8 | 31 |
| Go omission rate | 0.84 | 0.000 | 0.86 | 0.000 | 0.90 | 0.000 | 7 | 5 |
| Go Omission – SFR | 0.91 | 0.000 | 0.85 | 0.000 | 0.94 | 0.000 | 3 | 11 |
| **Within-subject correlation** | | | | | | | | |
| SFR : SSD | 0.17 | 0.000 | 0.23 | 0.000 | 0.63 | 0.000 | 271 | 174 |
| SFR : nSSD | 0.52 | 0.000 | 0.26 | 0.000 | 0.73 | 0.000 | 40 | 181 |

### Table S6. Individual Model Fit – RMSE and model comparison.

| RMSE | RVM | FSM | PRAD | PRAD % Improvement | |
|---|---|---|---|---|---|
| | | | | vs RVM | vs FSM |
| **RT related measures** | | | | | |
| Go RT | 0.044 | 0.043 | 0.041 | 7 | 5 |
| Stop RT | 0.057 | 0.052 | 0.043 | 25 | 17 |
| Stop RT – Go RT | 0.055 | 0.052 | 0.050 | 9 | 4 |
| Post-Go RT | 0.047 | 0.047 | 0.042 | 11 | 11 |
| Post-Stop RT | 0.059 | 0.056 | 0.048 | 19 | 14 |
| Post Stop RT – Post Go RT | 0.057 | 0.057 | 0.044 | 23 | 23 |
| Within SD RT | 0.112 | 0.115 | 0.085 | 24 | 26 |
| **Error related measures** | | | | | |
| SFR | 0.074 | 0.140 | 0.043 | 42 | 69 |
| Go omission rate | 0.091 | 0.086 | 0.065 | 29 | 24 |
| Go Omission – SFR | 0.061 | 0.097 | 0.059 | 3 | 39 |
| **Within-subject correlation** | | | | | |
| %SFR : SSD | 0.368 | 0.363 | 0.266 | 28 | 27 |
| SFR : nSSD | 0.350 | 0.432 | 0.363 | -4 | 16 |

**Table S7.** Divergence in distribution of RT related individual differences, measured by KL-divergence between the distribution of observed values across individuals and corresponding model posterior values.

| KL divergence | PRAD | RVM | FSM | PRAD % Improvement | |
|---|---|---|---|---|---|
| | | | | vs RVM | vs FSM |
| Go RT | 0.52 | 0.73 | 0.61 | 29 | 14 |
| Stop RT | 0.28 | 0.42 | 0.29 | 34 | 4 |
| Stop RT – Go RT | 0.89 | 2.94 | 1.89 | 70 | 53 |
| Post-Go RT | 0.66 | 1.04 | 0.95 | 36 | 30 |
| Post-Stop RT | 0.65 | 1.28 | 0.81 | 49 | 19 |
| Post Stop RT – Post Go RT | 1.22 | 11.56 | 7.32 | 89 | 83 |
| Within SD RT | 8.06 | 46.77 | 46.49 | 83 | 83 |
| SFR | 1.16 | 2.44 | 3.30 | 52 | 65 |
| Go omission rate | 1.00 | 1.23 | 1.20 | 18 | 17 |
| Go Omission – SFR | 0.24 | 0.28 | 0.51 | 14 | 53 |

**Table S8.** We evaluated whether for each individual, observed (RT, choice errors CE, Go-Stop errors GSR, SFR) and latent (SSRT, PDR, stopping bias, probability of proactivity) trial-by-trial behaviors showed significant correlations with one or more trial-level covariates such as SSD, number of trials since the last stop signal was encountered (nSSD), the average SSD encountered so far (xSSD), whether a left versus right error was made on the previous trial (postCE), whether the previous trial was a successful go versus stop trial (postGSE), and whether the previous trial was an inhibitory trial (postS).

| | **Observed** | | | | **Latent** | | | |
|---|---|---|---|---|---|---|---|---|
| | **RT** | **CE** | **GSR** | **SFR** | **SSRT** | **PDR** | **Bias** | **P PRO** |
| SSD | 31% | 15% | 71% | 71% | 9% | 94% | 4% | 11% |
| nSSD | 23% | 15% | 11% | 11% | 76% | 27% | 100% | 27% |
| xSSD | 24% | 10% | 48% | 49% | 7% | 100% | 23% | 54% |
| postCE | 15% | 14% | 6% | 5% | 6% | 13% | 10% | 19% |
| postGSE | 34% | 7% | 22% | 10% | 4% | 51% | 93% | 36% |
| postS | 42% | 9% | 47% | 0% | 0% | 0% | 100% | 42% |
| At least 1 | 81% | 50% | 95% | 85% | 82% | 100% | 100% | 84% |
| At least 2 | 53% | 16% | 73% | 51% | 18% | 98% | 100% | 55% |

**Table S9. PRAD latent dynamic measures.**

| Across individuals | Mean | SD | 95% CI | |
|---|---|---|---|---|
| SSRT (mS) | 350 | 155 | 196 | 749 |
| Average $\beta_S$ | 0.515 | 0.055 | 0.432 | 0.664 |
| PDR (mS) | 150 | 86 | 8 | 321 |
| P(PDR) | 0.776 | 0.253 | 0.03 | 0.984 |
| **Coefficient of Variation** | **Mean** | **SD** | **95% CI** | |
| SSRTCV | 0.173 | 0.079 | 0.072 | 0.371 |
| Average $\beta_S$ CV | 0.083 | 0.06 | 0.005 | 0.229 |
| PDRCV | 0.599 | 0.339 | 0.276 | 1.567 |
| P(PDR) CV | 0.288 | 0.288 | 0.029 | 1.195 |
| **Within individual 95% CI Range** | **Mean** | **SD** | **95% CI** | |
| SSRT (mS) | 238 | 151 | 73 | 626 |
| Average $\beta_S$ | 0.163 | 0.12 | 0.007 | 0.449 |
| PDR (mS) | 268 | 145 | 36 | 618 |
| P(PDR) | 0.564 | 0.371 | 0.026 | 0.988 |

**Table S10. Factor Analysis – 3 factor model.**

| | | Factor Loadings | | | Factor Weights | | |
|---|---|---|---|---|---|---|---|
| | | **f1** | **f2** | **f3** | **f1** | **f2** | **f3** |
| $\theta_0$ | PDR | 0.679 | 0.110 | -0.003 | 0.474 | 0.006 | 0.003 |
| $\theta_1$ | PDR | -0.574 | 0.083 | 0.019 | -0.327 | -0.001 | 0.002 |
| $\mu$ | PDR | 0.469 | -0.204 | 0.030 | 0.241 | -0.002 | 0.006 |
| $\delta_S$ | Reactive | -0.017 | -0.767 | -0.079 | -0.022 | -0.027 | -0.036 |
| $\alpha_S$ | Reactive | -0.008 | 0.997 | -0.031 | -0.022 | 0.969 | -0.001 |
| $\gamma_1$ | AMS | -0.029 | -0.076 | -0.686 | -0.025 | -0.008 | -0.184 |
| $\gamma_0$ | AMS | -0.014 | -0.029 | 0.922 | -0.018 | 0.022 | 0.795 |

**Table S11. Factor Analysis – comparison of models.**

| | $\chi^2$ | **df** | **p** | **RMSEA** | **CFI** | **TLI** | **BIC** |
|---|---|---|---|---|---|---|---|
| Criteria | | | < 0.05 | < 0.08 | > 0.95 | > 0.95 | lower is better |
| 3-factor | 50.5 | 3 | **0** | **0.045** | **0.997** | **0.977** | 140.5k |
| 2-factor | 2466 | 8 | **0** | 0.199 | 0.83 | 0.55 | 142.8k |
| 1-factor | 6571 | 14 | **0** | 0.245 | 0.55 | 0.32 | 146.9k |

**Table S12. Between-subject regressions show a significant relationship between almost all PRAD parameters and observed (SFR, xSSD, RT, RTCV) and latent measures (SSRT, SSRTCV) related to inhibitory control.**

| *Dependent variable* | | **SSRT** | **SFR** | **xSSD** | **SSRTCV** | **RT** | **RTCV** |
|---|---|---|---|---|---|---|---|
| $R^2$ | | 0.86 | 0.65 | 0.77 | 0.89 | 0.67 | 0.68 |
| F | | 3349 | 1053 | 1831 | 4442 | 1117 | 1194 |
| p | | < 0.0001 | < 0.0001 | < 0.0001 | < 0.0001 | < 0.0001 | < 0.0001 |
| $\delta_G$ | | $0.10^{***}$ | $0.30^{***}$ | $-0.30^{***}$ | $0.08^{***}$ | $-0.36^{***}$ | $-0.21^{***}$ |
| $\alpha_{G1}$ | | $0.11^{***}$ | $-0.20^{***}$ | $0.29^{***}$ | $-0.06^{***}$ | $0.56^{***}$ | $-0.32^{***}$ |
| $\alpha_{G2}$ | | $-0.02^{**}$ | $-0.42^{***}$ | $0.22^{***}$ | $-0.03^{***}$ | $0.13^{***}$ | $-0.14^{***}$ |
| $\kappa_0$ | | $-0.05^{***}$ | $-0.16^{***}$ | $0.37^{***}$ | $-0.03^{***}$ | $0.54^{***}$ | $-0.30^{***}$ |
| $\beta_G$ | | 0 | $-0.02^{*}$ | $0.04^{***}$ | $0.01^{*}$ | $-0.03^{***}$ | 0 |
| $\delta_S$ | Factor 2 | $-0.35^{***}$ | $-0.08^{***}$ | $0.22^{***}$ | $-0.86^{***}$ | $0.06^{***}$ | $-0.19^{***}$ |
| $\alpha_S$ | Factor 2 | $0.70^{***}$ | $0.80^{***}$ | $-0.59^{***}$ | $-0.66^{***}$ | $-0.38^{***}$ | $0.22^{***}$ |
| $\theta_0$ | Factor 1 | $-0.03^{***}$ | $-0.45^{***}$ | $0.10^{***}$ | -0.01 | $0.47^{***}$ | $-0.54^{***}$ |
| $\lvert\theta_1\rvert$ | Factor 1 | 0 | $-0.11^{***}$ | $0.07^{***}$ | $-0.02^{***}$ | $-0.03^{***}$ | $0.03^{***}$ |
| $\mu$ | Factor 1 | $0.02^{**}$ | $0.07^{***}$ | $-0.05^{***}$ | 0 | $-0.03^{***}$ | $0.05^{***}$ |
| $\lvert\gamma_1\rvert$ | Factor 3 | $-0.16^{***}$ | $-0.19^{***}$ | $0.18^{***}$ | $0.71^{***}$ | $0.11^{***}$ | $-0.07^{***}$ |
| $\gamma_0$ | Factor 3 | $-0.10^{***}$ | $-0.10^{***}$ | $0.09^{***}$ | $0.17^{***}$ | $0.05^{***}$ | $-0.04^{***}$ |
| $\tau_G$ | | $-0.30^{***}$ | $-0.55^{***}$ | $0.41^{***}$ | -0.05 | $0.29^{***}$ | $-0.58^{***}$ |
| $\tau_S$ | | $0.40^{***}$ | $0.52^{***}$ | $-0.55^{***}$ | $-0.09^{***}$ | $-0.22^{***}$ | $0.37^{***}$ |

**Table S13. PDR Regressions.** Regressions of individual level SFR and xSSD, measures of regression fit and standardizes correlation coefficients are reported. N/A indicates that variable was not included in a particular regression.

| | SFR | xSSD | SFR |
|---|---|---|---|
| $R^2$ | 0.48 | 0.46 | 0.65 |
| F | 3600 | 3270 | 3630 |
| p-value | < 0.0001 | < 0.0001 | < 0.0001 |
| PDR | $-0.44^{***}$ | $0.48^{***}$ | -0.02 |
| SSRT | $0.39^{***}$ | $-0.32^{***}$ | $0.20^{***}$ |
| xSSD | n/a | n/a | $-0.70^{***}$ |
| p (Proactive) | n/a | n/a | $-0.15^{***}$ |

**Table S14. AMS Regressions.** Regressions of individual level SSRT, SSRTCV, and log ratio of SSRT to RT: measures of regression fit and standardizes correlation coefficients are reported. N/A indicates that variable was not included in a particular regression.

| | **SSRT** | **SSRTCV** | **log(SSRT/RT)** |
|---|---|---|---|
| $R^2$ | 0.84 | 0.83 | 0.74 |
| F | 10300 | 9380 | 11000 |
| p-value | < 0.0001 | < 0.0001 | < 0.0001 |
| $\delta_S$ | $-0.38^{***}$ | $-0.79^{***}$ | $-0.27^{***}$ |
| $\alpha_S$ | $0.64^{***}$ | $-0.62^{***}$ | $0.63^{***}$ |
| $\tau_S$ | $0.15^{***}$ | n/a | n/a |
| Average $\beta_S$ | $-0.21^{***}$ | $0.36^{***}$ | n/a |
| $\beta_S$ CV | n/a | $0.55^{***}$ | n/a |

**Table S15.** Nonergodicity. Comparison of within-subject correlations over trials (average correlation coefficients across individuals reported) with the between-subject correlations over individuals based on mean individual level values. Differences, especially directional differences (positive vs negative) are indicative of nonergodicity.

| Correlations | Average Within | %Positive | Between | p-value (Between) |
|---|---|---|---|---|
| xSSD : RT | 0.055 | 66 | 0.737 | < 0.0001 |
| **xSSD : SFR** | **0.222** | **95** | **-0.768** | **< 0.0001** |
| xSSD : $p_{pro}$ | -0.100 | 14 | -0.308 | < 0.0001 |
| xSSD : PDR | 0.674 | 99 | 0.607 | < 0.0001 |
| **xSSD : SSRT** | **0.049** | **65** | **-0.510** | **< 0.0001** |
| **RT : $\boldsymbol{p_{pro}}$** | **0.421** | **99.8** | **-0.052** | **< 0.0001** |
| RT : PDR | 0.360 | 97 | 0.584 | < 0.0001 |
| **RT : SSRT** | **0.559** | **99** | **-0.129** | **< 0.0001** |
| **SFR : $\boldsymbol{p_{pro}}$** | **-0.313** | **2** | **0.065** | **< 0.0001** |
| **SFR : PDR** | **-0.074** | **45** | **-0.592** | **< 0.0001** |
| SFR : SSRT | 0.583 | 99.9 | 0.562 | < 0.0001 |
| SSRT : $p_{pro}$ | -0.036 | 37 | 0.018 | 0.122 |
| **SSRT : PDR** | **0.010** | **55** | **-0.388** | **< 0.0001** |
| $p_{pro}$ : PDR | 0.503 | 99.8 | 0.492 | < 0.0001 |

**Table S16. SSRT Bias.** The table shows the difference between PRAD model inferred SSRT and traditional non-parametric iSSRT (integrated method), grouped by various measures (4 quantiles of RT skewness, 4 quantiles of RT slowdown, 3 groups based on stop success rate, 2 groups based on go omission rates, 2 groups based on violations of context independence). In each case, monotonic increases across the group along with significant group differences based on ANOVA and t-tests indicate that PRAD can overcome systemic biases in traditional parametric iSSRT estimates.

| | | | | | **ANOVA** | | **t-test** | |
|---|---|---|---|---|---|---|---|---|
| **RT Skewness** | **Q1** | **Q2** | **Q3** | **Q4** | **F** | **p** | | |
| $\Delta\,(SSRT_{PRAD} - iSSRT)$ | 26 | 34 | 40 | 52 | 21.9 | <0.0001 | | |
| **RT Slowdown** | **Q1** | **Q2** | **Q3** | **Q4** | **F** | **p** | | |
| $\Delta\,(SSRT_{PRAD} - iSSRT)$ | 30 | 35 | 40 | 46 | 9.3 | <0.0001 | | |
| **Stop Success Rate** | **<=25%** | **25-75%** | **>=75%** | | **F** | **p** | | |
| $\Delta\,(SSRT_{PRAD} - iSSRT)$ | 30 | 37 | 229 | | 76.8 | <0.0001 | | |
| **Go Omission rate** | **<20%** | **>=20%** | | | | | **t** | **p** |
| $\Delta\,(SSRT_{PRAD} - iSSRT)$ | 37 | 52 | | | | | 3.4 | 0.0006 |
| **Violators (NDAR)** | **No** | **Yes** | | | | | **t** | **p** |
| $\Delta\,(SSRT_{PRAD} - iSSRT)$ | 29 | 156 | | | | | 28.9 | <0.0001 |

**Table S17. PRAD debiases inferences by reducing the difference in quality of inferences between typical and non-typical subgroups.** The table shows the effect sizes (Cohen's d) for differences between typical and atypical population subgroups, in the absolute values of model fit residuals for key measures (RT, RT SD, SFR and correlation between SFR and SSD), based on the PRAD and RVM models, as well as the percentage reduction in typicality bias (measured as the % reduction in difference between typical and atypical subgroups in absolute values of residuals from the RVM to PRAD). Positive values for the %reduction indicate that PRAD reduces typicality bias. Typicality is defined in terms of multiple variables including (i) family income (based on whether income was lower than or greater than $50k annually), (ii) age (median split), (iii) whether children demonstrated externalizing behavior (Ext., based on CBCL externalizing t-scores > 64), (iv) cognitive ability (Cog., median split using NIH cognitive toolbox scores), (v) brief problem monitoring assessment (BPM scale > 0), (vi) behavioral inhibition score (median split using the BIS subscale from BIS/BAS), and (vii) adverse life events (median split using ALE scores).

| | **Income** | **Age** | **Ext.** | **Cog.** | **BPM** | **BIS** | **ALE** |
|---|---|---|---|---|---|---|---|
| N (total) | 7139 | 7787 | 7782 | 7627 | 3477 | 7774 | 7366 |
| N (atypical) | 2081 | 3796 | 387 | 3587 | 2754 | 3954 | 4437 |
| N (typical) | 5058 | 3991 | 7395 | 4040 | 723 | 3820 | 2929 |
| **RT** | | | | | | | |
| Effect size (d) of typicality bias - PRAD | 0.15 | 0.21 | 0.12 | 0.33 | 0.08 | 0.04 | 0.01 |
| Effect size (d) of typicality bias - RVM | 0.19 | 0.33 | 0.12 | 0.44 | 0.16 | 0.05 | 0.04 |
| % Reduction in typicality bias (PRAD vs RVM) | 35% | 48% | 22% | 39% | 59% | 33% | 80% |
| **RTSD** | | | | | | | |
| Effect size (d) of typicality bias - PRAD | 0.25 | 0.17 | 0.20 | 0.36 | 0.24 | 0.07 | 0.10 |
| Effect size (d) of typicality bias - RVM | 0.38 | 0.34 | 0.35 | 0.61 | 0.30 | 0.12 | 0.13 |
| % Reduction in typicality bias (PRAD vs RVM) | 22% | 39% | 33% | 27% | 2% | 28% | 9% |
| **SFR** | | | | | | | |
| Effect size (d) of typicality bias - PRAD | 0.06 | 0.09 | 0.02 | 0.08 | 0.01 | -0.01 | 0.01 |
| Effect size (d) of typicality bias - RVM | 0.07 | 0.10 | 0.02 | 0.06 | 0.03 | 0.03 | 0.06 |
| % Reduction in typicality bias (PRAD vs RVM) | 45% | 39% | 34% | 16% | 69% | 113% | 84% |
| **Correlation (SFR, SSD)** | | | | | | | |
| Effect size (d) of typicality bias - PRAD | 0.09 | 0.03 | 0.17 | 0.10 | 0.10 | 0.01 | 0.05 |
| Effect size (d) of typicality bias - RVM | 0.22 | 0.07 | 0.21 | 0.20 | 0.16 | 0.07 | 0.06 |
| % Reduction in typicality bias (PRAD vs RVM) | 70% | 68% | 45% | 65% | 54% | 86% | 38% |

**Table S18.** NIH Cognitive Toolbox predictions using SVM regressions. The table shows the adjusted $R^2$ as well as correlations between model predictions and actual values.

| NIH Cognitive Toolbox | Fit | | Cross-validated | |
|---|---|---|---|---|
| **Predict using iSSRT** | **Adjusted $R^2$** | **Correlation** | **Adjusted $R^2$** | **Correlation** |
| Total score | 6.3% | $0.25^{***}$ | 4.6% | $0.22^{***}$ |
| Fluid sub-score | 6.2% | $0.25^{***}$ | 4.0% | $0.21^{***}$ |
| Crystallized sub-score | 3.4% | $0.19^{***}$ | 1.7% | $0.14^{***}$ |
| DCCS | 3.5% | $0.20^{***}$ | 1.7% | $0.15^{***}$ |
| PVT | 1.9% | $0.15^{***}$ | 0.4% | $0.10^{***}$ |
| LSWM | 2.4% | $0.16^{***}$ | 0.9% | $0.10^{***}$ |
| Flanker | 0.0% | $0.15^{***}$ | -1.5% | $0.09^{***}$ |
| PCPST | 4.2% | $0.21^{***}$ | 2.4% | $0.17^{***}$ |
| ORRT | 3.4% | $0.19^{***}$ | 1.9% | $0.14^{***}$ |
| **Predict using PRAD parameters** | **Adjusted $R^2$** | **Correlation** | **Adjusted $R^2$** | **Correlation** |
| Total score | 27.7% | $0.53^{***}$ | 15.3% | $0.40^{***}$ |
| Fluid sub-score | 25.6% | $0.51^{***}$ | 12.7% | $0.37^{***}$ |
| Crystallized sub-score | 19.5% | $0.45^{***}$ | 6.9% | $0.28^{***}$ |
| DCCS | 18.2% | $0.44^{***}$ | 7.5% | $0.29^{***}$ |
| PVT | 16.0% | $0.41^{***}$ | 3.0% | $0.21^{***}$ |
| LSWM | 14.7% | $0.39^{***}$ | 2.9% | $0.22^{***}$ |
| Flanker | 14.6% | $0.41^{***}$ | 3.6% | $0.25^{***}$ |
| PCPST | 17.6% | $0.43^{***}$ | 5.9% | $0.27^{***}$ |
| ORRT | 17.2% | $0.44^{***}$ | 7.0% | $0.27^{***}$ |

**Table S19. Reliability & Sensitivity Analysis**

| ICC | Across 2 runs (reliability) | PRAD vs PRAD-g (sensitivity) |
|---|---|---|
| N | 6737 | 7625 |
| **Observed Behavior** | | |
| RT | 0.85 | - |
| SFR | 0.12 | - |
| Go omission rate | 0.57 | - |
| Correlation (SFR, SSD) | 0.10 | - |
| Correlation (SFR, nSSD) | -0.01 | - |
| **PRAD individual parameters** | | |
| $\delta_G$ | 0.72 | 1.00 |
| $\alpha_{G1}$ | 0.44 | 0.97 |
| $\alpha_{G2}$ | 0.36 | 0.97 |
| $\kappa_0$ | 0.49 | n/a * |
| $\beta_G$ | 0.56 | 1.00 |
| $\delta_S$ | 0.53 | 0.99 |
| $\alpha_S$ | 0.59 | 0.96 |
| $\theta_0$ | 0.26 | 0.98 |
| $\theta_1$ | 0.25 | 1.00 |
| $\mu$ | 0.08 | 0.99 |
| $\gamma_1$ | 0.02 | 0.98 |
| $\gamma_0$ | 0.01 | 0.91 |
| $\tau_G$ | 0.22 | 0.98 |
| $\tau_S$ | 0.22 | 0.98 |
| **PRAD average of dynamic measures** | | |
| SSRT | 0.42 | 0.99 |
| $p_{PRO}$ | 0.22 | 0.97 |
| PDR | 0.46 | 0.98 |

* $\boldsymbol{\kappa_0}$ has a different connotation under PRAD-g and is hence not comparable

**Table S20. Sensitivity Analysis.** Comparison of key measures and inferences from primary analysis to three sets of control analysis: (1) NDT prior: Broader prior on fixed component ($\boldsymbol{\tau_G}$) of PRAD non-decision time (see **SI Table S20**). (2) Sampling 1: Analysis restricted to subset of participants with three or more converged chains (MCMC sampling). (3) Sampling 2: Analysis restricted to subset of participants with four converged chains (MCMC sampling). Results from the primary analysis are robust and not sensitive to sampling hyperparameters or choice of NDT prior. (4) PRAD-g: Full generative model that does not include stimulus duration effects on go trials.

| Key Measures / Inferences | SI Table | Primary Analysis | Sensitivity Analysis | | | |
|---|---|---|---|---|---|---|
| | | | NDT prior | Sampling 1 | Sampling 2 | PRAD-g |
| N | | 7787 | 7381 | 6604 | 4195 | 7625 |
| **%RMSE reduction - Aggregate vs RVM** | | **PRAD** | **NDT** | **Sampling 1** | **Sampling 2** | **PRAD-g** |
| SFR by unique SSD values | S4 | 66 | 66 | 68 | 76 | 68 |
| SFR by unique nSSD values | S4 | 58 | 58 | 60 | 65 | 53 |
| Mean RT on stop failure by unique SSD values | S4 | 45 | 47 | 31 | 0 | 46 |
| SD RT on stop failure by unique SSD values | S4 | 46 | 45 | 48 | 48 | 45 |
| Mean RT on stop failure by unique nSSD values | S4 | 50 | 57 | 50 | 54 | 60 |
| Choice accuracy on stop failure by unique SSD values | S4 | 48 | 44 | 39 | 30 | 47 |
| Choice accuracy on stop failure by unique nSSD values | S4 | 51 | 52 | 50 | 50 | 27 |
| **Individual Fit Correlations** | | **PRAD** | **NDT** | **Sampling 1** | **Sampling 2** | **PRAD-g** |
| Go RT | S5 | 0.91 | 0.91 | 0.91 | 0.91 | 0.91 |
| Stop RT | S5 | 0.90 | 0.90 | 0.90 | 0.90 | 0.90 |
| Stop RT – Go RT | S5 | 0.51 | 0.50 | 0.49 | 0.47 | 0.51 |
| Post-Go RT | S5 | 0.92 | 0.92 | 0.92 | 0.92 | 0.92 |
| Post-Stop RT | S5 | 0.86 | 0.86 | 0.86 | 0.86 | 0.86 |
| Post Stop RT – Post Go RT | S5 | 0.45 | 0.45 | 0.43 | 0.40 | 0.45 |
| Within-sub SD RT | S5 | 0.13 | 0.16 | 0.13 | 0.12 | 0.13 |
| Within-sub SD RT by SSD | S5 | 0.26 | 0.25 | 0.27 | 0.29 | 0.25 |
| SFR | S5 | 0.85 | 0.85 | 0.85 | 0.88 | 0.83 |
| Go omission rate | S5 | 0.90 | 0.92 | 0.90 | 0.91 | 0.91 |
| Go Omission – SFR | S5 | 0.94 | 0.94 | 0.94 | 0.95 | 0.93 |
| SFR : SSD | S5 | 0.63 | 0.64 | 0.64 | 0.67 | 0.63 |
| SFR : nSSD | S5 | 0.73 | 0.73 | 0.73 | 0.73 | 0.72 |
| **Individual Fit Corr improvement vs RVM** | | **PRAD** | **NDT** | **Sampling 1** | **Sampling 2** | **PRAD-g** |
| Go RT | S5 | 3 | 3 | 3 | 3 | 3 |
| Stop RT | S5 | 5 | 5 | 5 | 3 | 5 |
| Stop RT – Go RT | S5 | 28 | 28 | 29 | 27 | 31 |
| Post-Go RT | S5 | 6 | 6 | 5 | 6 | 5 |
| Post-Stop RT | S5 | 9 | 9 | 8 | 8 | 9 |
| Post Stop RT – Post Go RT | S5 | 2150 | 2150 | 2050 | Inf | 2150 |
| Within-sub SD RT | S5 | 244 | 260 | 230 | 186 | 244 |
| Within-sub SD RT by SSD | S5 | 0 | 0 | 4 | 7 | -3 |
| SFR | S5 | 8 | 6 | 6 | 7 | 5 |
| Go omission rate | S5 | 7 | 10 | 7 | 7 | 8 |
| Go Omission – SFR | S5 | 3 | 3 | 3 | 2 | 2 |
| SFR : SSD | S5 | 271 | 276 | 237 | 148 | 250 |
| SFR : nSSD | S5 | 40 | 40 | 43 | 43 | 39 |
| **Individual Fit RMSE** | | **PRAD** | **NDT** | **Sampling 1** | **Sampling 2** | **PRAD-g** |
| Go RT | S6 | 0.041 | 0.041 | 0.043 | 0.046 | 0.040 |
| Stop RT | S6 | 0.043 | 0.042 | 0.044 | 0.046 | 0.041 |

| | | | | | | |
|---|---|---|---|---|---|---|
| Stop RT – Go RT | S6 | 0.050 | 0.049 | 0.050 | 0.051 | 0.045 |
| Post-Go RT | S6 | 0.042 | 0.041 | 0.043 | 0.046 | 0.041 |
| Post-Stop RT | S6 | 0.048 | 0.048 | 0.049 | 0.051 | 0.048 |
| Post Stop RT – Post Go RT | S6 | 0.044 | 0.043 | 0.044 | 0.046 | 0.043 |
| Within-sub SD RT | S6 | 0.085 | 0.087 | 0.086 | 0.087 | 0.086 |
| Within-sub SD RT by SSD | S6 | 0.049 | 0.050 | 0.049 | 0.051 | |
| SFR | S6 | 0.043 | 0.044 | 0.043 | 0.044 | 0.044 |
| Go omission rate | S6 | 0.065 | 0.056 | 0.065 | 0.066 | 0.062 |
| Go Omission – SFR | S6 | 0.059 | 0.050 | 0.058 | 0.058 | 0.069 |
| SFR : SSD | S6 | 0.266 | 0.270 | 0.274 | 0.288 | 0.265 |
| SFR : nSSD | S6 | 0.363 | 0.361 | 0.365 | 0.368 | 0.367 |
| **Individual Fit RMSE improvement vs RVM** | | **PRAD** | **NDT** | **Sampling 1** | **Sampling 2** | **PRAD-g** |
| Go RT | S6 | 7 | 9 | 7 | 8 | 9 |
| Stop RT | S6 | 25 | 26 | 23 | 22 | 27 |
| Stop RT – Go RT | S6 | 9 | 11 | 9 | 6 | 17 |
| Post-Go RT | S6 | 11 | 15 | 12 | 13 | 13 |
| Post-Stop RT | S6 | 19 | 19 | 20 | 19 | 19 |
| Post Stop RT – Post Go RT | S6 | 23 | 25 | 23 | 21 | 25 |
| Within-sub SD RT | S6 | 24 | 23 | 25 | 25 | 23 |
| Within-sub SD RT by SSD | S6 | 14 | 14 | 16 | 15 | |
| SFR | S6 | 42 | 40 | 42 | 41 | 41 |
| Go omission rate | S6 | 29 | 39 | 29 | 30 | 32 |
| Go Omission – SFR | S6 | 3 | 18 | 5 | 8 | -15 |
| SFR : SSD | S6 | 28 | 28 | 28 | 31 | 28 |
| SFR : nSSD | S6 | -4 | -4 | -4 | -3 | -5 |
| **KLD** | | **PRAD** | **NDT** | **Sampling 1** | **Sampling 2** | **PRAD-g** |
| Go RT | S7 | 0.52 | 0.43 | 0.62 | 0.75 | 0.54 |
| Stop RT | S7 | 0.28 | 0.25 | 0.28 | 0.45 | 0.30 |
| Post-Go RT | S7 | 0.66 | 0.54 | 0.79 | 1.00 | 0.65 |
| Post-Stop RT | S7 | 0.65 | 0.49 | 0.60 | 0.82 | 0.60 |
| Stop RT – Go RT | S7 | 0.89 | 0.84 | 0.86 | 0.99 | 0.62 |
| Post Stop RT – Post Go RT | S7 | 1.22 | 1.14 | 1.23 | 1.31 | 1.10 |
| **KLD % improvement vs RVM** | | **PRAD** | **NDT** | **Sampling 1** | **Sampling 2** | **PRAD-g** |
| Go RT | S7 | 29 | 42 | 27 | 33 | 25 |
| Stop RT | S7 | 34 | 39 | 40 | 18 | 23 |
| Post-Go RT | S7 | 36 | 50 | 35 | 55 | 38 |
| Post-Stop RT | S7 | 49 | 62 | 61 | 59 | 53 |
| Stop RT – Go RT | S7 | 70 | 71 | 71 | 67 | 79 |
| Post Stop RT – Post Go RT | S7 | 89 | 90 | 90 | 92 | 90 |
| ***Factor Loadings*** | | **PRAD** | **NDT** | **Sampling 1** | **Sampling 2** | **PRAD-g** |
| $f1: \theta_0$ | S10 | 0.679 | 0.618 | 0.686 | 0.677 | 0.665 |
| $f1: \theta_1$ | S10 | -0.574 | -0.518 | -0.593 | -0.656 | -0.581 |
| $f1: \mu$ | S10 | 0.469 | 0.507 | 0.459 | 0.460 | 0.490 |
| $f2: \delta_S$ | S10 | -0.767 | -0.747 | -0.772 | -0.762 | -0.784 |
| $f2: \alpha_S$ | S10 | 0.997 | 1.022 | 0.994 | 1.012 | 1.008 |
| $f3: \gamma_1$ | S10 | -0.686 | -0.699 | -0.692 | -0.675 | -0.641 |
| $f3: \gamma_0$ | S10 | 0.922 | 1.008 | 0.947 | 0.991 | 0.915 |
| ***Factor Weights*** | | **PRAD** | **NDT** | **Sampling 1** | **Sampling 2** | **PRAD-g** |
| $f1: \theta_0$ | S10 | 0.474 | 0.449 | 0.471 | 0.422 | 0.451 |
| $f1: \theta_1$ | S10 | -0.327 | -0.297 | -0.342 | -0.403 | -0.336 |
| $f1: \mu$ | S10 | 0.241 | 0.300 | 0.228 | 0.216 | 0.258 |
| $f2: \delta_S$ | S10 | -0.027 | 0.073 | -0.039 | 0.033 | 0.016 |

| | | | | | | |
|---|---|---|---|---|---|---|
| $f2{:}\,\alpha_S$ | S10 | 0.969 | 1.080 | 0.955 | 1.033 | 1.014 |
| $f3{:}\,\gamma_1$ | S10 | -0.184 | 0.017 | -0.136 | -0.028 | -0.169 |
| $f3{:}\,\gamma_0$ | S10 | 0.795 | 1.017 | 0.848 | 0.967 | 0.805 |
| **Factor Model Criteria** | | **PRAD** | **NDT** | **Sampling 1** | **Sampling 2** | **PRAD-g** |
| 3-factor model significant; 2-f and 1-f not significant | S11 | Yes | Yes | Yes | Yes | Yes |
| 3-factor: RMSEA | S11 | 0.045 | 0.027 | 0.051 | 0.045 | 0.048 |
| 2-factor: RMSEA | S11 | 0.199 | 0.181 | 0.203 | 0.214 | 0.202 |
| 1-factor: RMSEA | S11 | 0.245 | 0.265 | 0.253 | 0.263 | 0.232 |
| 3-factor: CFI | S11 | 0.997 | 0.999 | 0.996 | 0.997 | 0.996 |
| 2-factor: CFI | S11 | 0.83 | 0.87 | 0.83 | 0.819 | 0.826 |
| 1-factor: CFI | S11 | 0.547 | 0.514 | 0.535 | 0.52 | 0.596 |
| 3-factor: TLI | S11 | 0.977 | 0.992 | 0.972 | 0.979 | 0.974 |
| 2-factor: TLI | S11 | 0.554 | 0.658 | 0.553 | 0.524 | 0.543 |
| 1-factor: TLI | S11 | 0.32 | 0.271 | 0.302 | 0.281 | 0.394 |
| **Regressions ($R^2$)** | | **PRAD** | **NDT** | **Sampling 1** | **Sampling 2** | **PRAD-g** |
| SSRT | S12 | 0.86 | 0.84 | 0.87 | 0.87 | 0.86 |
| SFR | S12 | 0.65 | 0.65 | 0.67 | 0.7 | 0.64 |
| xSSD | S12 | 0.77 | 0.75 | 0.77 | 0.79 | 0.73 |
| SSRTCV | S12 | 0.89 | 0.85 | 0.89 | 0.89 | 0.89 |
| RT | S12 | 0.67 | 0.67 | 0.69 | 0.71 | 0.57 |
| RTCV | S12 | 0.68 | 0.68 | 0.69 | 0.7 | 0.65 |
| **PDR Regressions ($R^2$)** | | **PRAD** | **NDT** | **Sampling 1** | **Sampling 2** | **PRAD-g** |
| SFR 1 | S13 | 0.48 | 0.45 | 0.49 | 0.51 | 0.47 |
| xSSD | S13 | 0.46 | 0.47 | 0.48 | 0.51 | 0.45 |
| SFR 2 | S13 | 0.65 | 0.64 | 0.66 | 0.67 | 0.65 |
| **AMS Regressions ($R^2$)** | | **PRAD** | **NDT** | **Sampling 1** | **Sampling 2** | **PRAD-g** |
| SSRT | S14 | 0.84 | 0.82 | 0.85 | 0.86 | 0.85 |
| SSRTCV | S14 | 0.83 | 0.78 | 0.83 | 0.83 | 0.83 |
| log(SSRT/RT) | S14 | 0.74 | 0.62 | 0.73 | 0.72 | 0.73 |
| **Within Correlations** | | **PRAD** | **NDT** | **Sampling 1** | **Sampling 2** | **PRAD-g** |
| xSSD : SFR | S15 | 0.22 | 0.22 | 0.22 | 0.21 | 0.22 |
| xSSD : SSRT | S15 | 0.05 | 0.05 | 0.05 | 0.05 | 0.05 |
| RT : p_pro | S15 | 0.42 | 0.46 | 0.42 | 0.42 | 0.42 |
| RT : SSRT | S15 | 0.56 | 0.56 | 0.56 | 0.56 | 0.55 |
| SFR : p_pro | S15 | -0.31 | -0.36 | -0.31 | -0.32 | -0.32 |
| SFR : PDR | S15 | -0.07 | -0.13 | -0.07 | -0.08 | -0.08 |
| SSRT : PDR | S15 | 0.01 | 0.00 | 0.01 | 0.01 | 0.01 |
| **Between Correlations** | | **PRAD** | **NDT** | **Sampling 1** | **Sampling 2** | **PRAD-g** |
| xSSD : SFR | S15 | -0.77 | -0.77 | -0.77 | -0.79 | -0.77 |
| xSSD : SSRT | S15 | -0.51 | -0.51 | -0.52 | -0.52 | -0.52 |
| RT : p_pro | S15 | -0.05 | 0.01 | -0.04 | -0.03 | -0.06 |
| RT : SSRT | S15 | -0.13 | -0.13 | -0.16 | -0.21 | -0.14 |
| SFR : p_pro | S15 | 0.06 | 0.05 | 0.06 | 0.07 | 0.07 |
| SFR : PDR | S15 | -0.59 | -0.54 | -0.60 | -0.62 | -0.58 |
| SSRT : PDR | S15 | -0.39 | -0.36 | -0.40 | -0.42 | -0.39 |
| **Change in sign** | | **PRAD** | **NDT** | **Sampling 1** | **Sampling 2** | **PRAD-g** |
| xSSD : SFR | S15 | Yes | Yes | Yes | Yes | Yes |
| xSSD : SSRT | S15 | Yes | Yes | Yes | Yes | Yes |
| RT : p_pro | S15 | Yes | No | Yes | Yes | Yes |
| RT : SSRT | S15 | Yes | Yes | Yes | Yes | Yes |
| SFR : p_pro | S15 | Yes | Yes | Yes | Yes | Yes |
| SFR : PDR | S15 | No | No | No | No | No |
| SSRT : PDR | S15 | Yes | Yes | Yes | Yes | Yes |

| SSRT Bias | | PRAD | NDT | Sampling 1 | Sampling 2 | PRAD-g |
|---|---|---|---|---|---|---|
| RT skewness - F | S16 | 21.9 | 17.0 | 22.2 | 14.5 | 26.2 |
| RT slowdown - F | S16 | 9.3 | 8.8 | 4.7 | 2.9 | 8.91 |
| Stop success rate - F | S16 | 76.8 | 66.7 | 57.5 | 35.4 | 59.7 |
| Go Omission - t | S16 | 3.4 | 3.3 | 2.1 | 1.1 | 1.9 |
| Violators - t | S16 | 28.9 | 24.6 | 21.7 | 11.2 | 27.7 |
| RT skewness - Monotonically increasing | S16 | Yes | Yes | Yes | Yes | Yes |
| RT slowdown - Monotonically increasing | S16 | Yes | Yes | Yes | Yes | Yes |
| Stop success rate - Monotonically increasing | S16 | Yes | Yes | Yes | Yes | Yes |
| Go Omission - Monotonically increasing | S16 | Yes | Yes | Yes | Yes | Yes |
| Violators - Monotonically increasing | S16 | Yes | Yes | Yes | Yes | Yes |
| **% Reduction in Typicality Bias (PRAD vs RVM)** | | **PRAD** | **NDT** | **Sampling 1** | **Sampling 2** | **PRAD-g** |
| RT (Family income) | S17 | 35 | 33 | 31 | 20 | 35 |
| RTSD (Family income) | S17 | 22 | 23 | 24 | 28 | 22 |
| Corr [SFR,SSD] (Family income) | S17 | 70 | 70 | 68 | 72 | 70 |
| RT (Age) | S17 | 48 | 54 | 48 | 50 | 47 |
| RTSD (Age) | S17 | 39 | 40 | 46 | 55 | 45 |
| Corr [SFR,SSD] (Age) | S17 | 68 | 69 | 85 | 120 | 68 |
| RT (Cognitive) | S17 | 39 | 42 | 40 | 36 | 40 |
| RTSD (Cognitive) | S17 | 27 | 28 | 27 | 31 | 29 |
| Corr [SFR,SSD] (Cognitive) | S17 | 65 | 66 | 66 | 76 | 66 |
| RT (BPM) | S17 | 59 | 64 | 65 | 102 | 60 |
| RTSD (BPM) | S17 | 2 | -4 | 7 | -16 | 8 |
| Corr [SFR,SSD] (BPM) | S17 | 54 | 53 | 64 | 62 | 53 |
| RT (BIS) | S17 | 33 | 29 | 31 | 23 | 41 |
| RTSD (BIS) | S17 | 28 | 23 | 28 | 12 | 25 |
| Corr [SFR,SSD] (BIS) | S17 | 86 | 85 | 87 | 98 | 91 |
| RT (ALE) | S17 | 80 | 95 | 65 | 80 | 85 |
| RTSD (ALE) | S17 | 9 | 14 | 8 | 7 | 11 |
| Corr [SFR,SSD] (ALE) | S17 | 38 | 39 | 40 | 6 | 44 |
| **NIH Cognitive Toolbox: CV Correlations** | | **PRAD** | **NDT** | **Sampling 1** | **Sampling 2** | **PRAD-g** |
| PRAD > iSSRT on all measures | S18 | Yes | Yes | Yes | Yes | Yes |
| iSSRT: Total | S18 | 0.22 | 0.22 | 0.21 | 0.21 | 0.22 |
| iSSRT: Fluid | S18 | 0.21 | 0.21 | 0.20 | 0.19 | 0.20 |
| iSSRT: Crystallized | S18 | 0.14 | 0.14 | 0.14 | 0.14 | 0.14 |
| iSSRT: DCCS | S18 | 0.15 | 0.15 | 0.14 | 0.14 | 0.15 |
| iSSRT: PVT | S18 | 0.10 | 0.10 | 0.11 | 0.11 | 0.10 |
| iSSRT: LSWM | S18 | 0.10 | 0.11 | 0.09 | 0.08 | 0.09 |
| iSSRT: Flanker | S18 | 0.09 | 0.09 | 0.08 | 0.09 | 0.08 |
| iSSRT: PCPST | S18 | 0.17 | 0.16 | 0.15 | 0.14 | 0.16 |
| iSSRT: ORRT | S18 | 0.14 | 0.14 | 0.14 | 0.12 | 0.14 |
| PRAD: Total | S18 | 0.40 | 0.40 | 0.40 | 0.40 | 0.39 |
| PRAD: Fluid | S18 | 0.37 | 0.36 | 0.37 | 0.38 | 0.36 |
| PRAD: Crystallized | S18 | 0.28 | 0.27 | 0.28 | 0.26 | 0.27 |
| PRAD: DCCS | S18 | 0.29 | 0.29 | 0.29 | 0.28 | 0.28 |
| PRAD: PVT | S18 | 0.21 | 0.21 | 0.20 | 0.19 | 0.20 |
| PRAD: LSWM | S18 | 0.22 | 0.21 | 0.21 | 0.23 | 0.21 |
| PRAD: Flanker | S18 | 0.25 | 0.25 | 0.25 | 0.26 | 0.24 |
| PRAD: PCPST | S18 | 0.27 | 0.27 | 0.27 | 0.26 | 0.26 |
| PRAD: ORRT | S18 | 0.27 | 0.27 | 0.28 | 0.28 | 0.27 |

**Table S21. Participants and demographics.**

| | |
|---|---|
| Total | N = 7787 |
| Female | N = 3810 (48.9%) |
| Age | M = 118.9 months; SD = 7.4 months; 95%CI (108 – 131 months) |
| Family Income < $50000 | N = 2081 (26.7%) |
| NIH Cognitive toolbox scores | M = 86.7 (N 7627); SD = 8.9; 95%CI (68 – 103) |

### Table S22. Bayesian priors.

| Parameter | PRAD / PRAD-g | RVM | FSM |
|---|---|---|---|
| $\delta_G$ | Unif (0.0001, 12) | Unif (0.0001, 12) | Unif (0.0001, 12) |
| $\alpha_{G1}$ | Unif (0.5, 4) | Unif (0.5, 4) | Unif (0.5, 4) |
| $\alpha_{G2}$ | Unif (0.5, 4) | n/a | n/a |
| $\kappa_0$ | Normal (0.1, prec = 0.3) T(0,0.3) | n/a | n/a |
| $\beta_G$ | Beta (1,1) T(0.0001,0.9999) | Beta (1,1) T(0.0001,0.9999) | Beta (1,1) T(0.0001,0.9999) |
| $\delta_S$ | $\delta_S = \frac{p}{\alpha_S}$ ; $p$ ~ Unif (04, 24) | $\delta_S = \frac{p}{\alpha_S}$ ; $p$ ~ Unif (04, 24) | n/a |
| $\alpha_S$ | Unif (0.5, 6) | Unif (0.5, 6) | n/a |
| $\theta_0$ | Normal (0, prec = 0.3) | n/a | n/a |
| $\theta_1$ | Normal (0, prec = 0.3) | n/a | n/a |
| $\mu$ | Beta (0.5, 0.5) T(0.8, 0.9999) | n/a | n/a |
| $\gamma_1$ | Normal (0, prec = 0.3) | n/a | n/a |
| $\gamma_0$ | Unif (0, 12) | n/a | n/a |
| $\tau_G$ | Unif (0.0001, min(0.1**,RT*)) | n/a | n/a |
| $\tau_S$ | Unif (0.0001, $\tau_G$) | Uniform (0.0001, $\tau_G$) | n/a |
| Go NDT | n/a | Unif (0.0001, RT*) | Unif (0.0001, RT*) |
| SSRT | n/a | n/a | Unif (0.01,1) |

*Min RT after censoring for RT<0.05s
** Additional NDT control analysis: Unif (0.0001, min(0.3,RT*))
